# Systematic genome analysis enables long-overdue taxonomic reorganization of genus *Orthoflavivirus*; potential assignment of over 50 new species

**DOI:** 10.64898/2026.10.21.760428

**Authors:** Peter Simmonds, Martin Beer, Nicolas Bejerman, Jens Bukh, Anamarija Butković, Agathe Colmant, Humberto Debat, J. Felix Drexler, Amit Kapoor, Jens H Kuhn, Volker Lohmann, Donald B Smith, Jack T Stapleton, Nikos Vasilakis

**Affiliations:** Department of Biomedicine, University of Turku, Turku, Finland; Institute of Diagnostic Virology, Friedrich-Loeffler-Institut, Federal Research Institute for Animal Health, Greifswald-Insel Riems, Germany; Consejo Nacional de Investigaciones Científicas y Técnicas (CONICET), Buenos Aires CP1033, Argentina; Instituto de Patologia Vegetal (IPAVE)-Instituto Nacional de Tecnología Agropecuaria (INTA), Córdoba CP5019, Argentina; Copenhagen Hepatitis C Program (CO-HEP), Department of Immunology and Microbiology, Faculty of Health and Medical Sciences, University of Copenhagen, Denmark; CO-HEP, Department of Infectious Diseases, Copenhagen University Hospital, Hvidovre, Denmark; Department of Virology and Microbiological Preparedness, Statens Serum Institut, Copenhagen, Denmark; Independent Researcher, Institut Pasteur, Paris, France; Unité Des Virus Emergents (UVE: Aix-Marseille University, Universita Di Corsica, IRD 190, Inserm 1207 IRBA), Marseille, France; Charité-Universitätsmedizin Berlin, corporate member of Freie Universität Berlin and Humboldt-Universität zu Berlin, Institute of Virology, Berlin, Germany; Center for Vaccines and Immunity, The Research Institute at Nationwide Children’s Hospital, Columbus, Ohio, USA; Indepdendent Researcher, Fort Detrick, Frederick, Maryland, USA; Department of Infectious Diseases, Molecular Virology, Heidelberg University, Heidelberg, Germany; Centre for Virus Research, University of Glasgow, Glasgow, UK; Departments of Internal Medicine, Microbiology and Immunology, University of Iowa and Iowa City VA Healthcare, Iowa City, Iowa, USA; Department of Pathology and Center for Vector-Borne and Zoonotic Diseases, University of Texas Medical Branch, Galveston, Texas, USA

## Abstract

Members of the *Orthoflavivirus* genus in the family *Flaviviridae* are primarily vector-transmitted RNA viruses infecting humans, other mammals and birds, including major pathogens such as tick borne encephalitis, West Nile, Zika, yellow fever and dengue viruses. Vector-borne flaviviruses are assigned to the *Euflavivirus* subgenus of orthoflaviviruses, with more divergent largely unclassified insect-specific flaviviruses, including cell fusing agent virus assigned to the second subgenus, *Fusivirus*. The currently classified 57 orthoflavivirus species have traditionally been assigned descriptively based on their genome and antigenic relatedness, ecology and host and vector tropisms, although many more orthoflavi-like viruses remain unassigned. To investigate whether species might be demarcated using a fixed genetic divergence threshold, we examined distributions of pairwise distances in *NS5* and *E* genes (which encode the RNA-dependent RNA polymerase and envelope protein, respectively) of classified orthoflaviviruses. Demarcation thresholds of 0.1 in the two regions would be most parsimonious with current euflavivirus classification. Exceptions include species in the tick-borne encephalitis virus complex with divergences below the threshold, and dengue virus types currently assigned to a single species that are substantially above. Application of the proposed thresholds would provide the means to establish 25 new euflavivirus and 33 new fusivirus species and reconcile several anomalies and inconsistencies in the current species taxonomy. The use of a species demarcation threshold brings consistency with the classification of viruses in other families. A robust and evolutionarily consistent framework furthermore enables taxonomic assignments of the numerous currently unassigned orthoflaviviruses that lack the phenotypic descriptions required for the current species definition.

## Introduction

The *Flaviviridae* is a family of positive-sense, enveloped RNA viruses that infect humans, other mammals and birds and which are primarily transmitted through mosquito and tick vectors (1, 2). The family has been reorganised many times, reflecting the growing knowledge of the evolutionary relationships of its viruses to other positive-sense viruses and increasing data on the genetic diversity of the family. Yellow fever virus (YFV) served as the founding virus for the family and was used to name the family (Latin *flavus*, meaning “yellow”) and its genus *Flavivirus*. Distantly related viruses, causing respiratory and enteric infection in domestic mammals were added as a second genus, *Pestivirus*, followed by the additions of further genera of primarily mammalian viruses, including hepatitis C virus (HCV) in the genus *Hepacivirus* (3) and human pegivirus type 1 in the genus *Pegivirus* (4).

Recently, a major reorganisation of the family added the phylogenetically related tick-vectored flavivirids Jīngmén tick virus (JMTV) and Ālóngshān viruses (ALSV), the mosquito-infecting Guaico Culex virus (GCXV), the mammalian Tamana bat virus (TaBV) and the termite-infecting waxsystermes virus (WXTV) by way of establishment of four genera (*Jingmenvirus*, *Guaicovirus*, *Tamanavirus*, and *Termitovirus*, respectively). The far more divergent hepaciviruses and pestiviruses were reclassified into separate families, *Hepaciviridae* and *Pestiviridae* (2). Mosquito- and tick-borne flavivirids are assigned to genus *Orthoflavivirus* and include multiple human pathogens of significant public health impact, including dengue viruses type 1–4 (DENV-1–4), Japanese encephalitis virus (JEV), tick-borne encephalitis viruses 1 and 2 (TBEV-1/2), West Nile virus (WNV), yellow fever virus (YFV), and Zika virus (ZIKV) (1, 2). In total, the genus *Orthoflavivirus* comprises 57 species divided into three subgenera, *Crangovirus*, *Euflavivirus*, and *Fusivirus* (Table S1) (2), with all vector-borne viruses being euflaviviruses that are further clustered into several informal subgroups reflecting their transmission vectors or absence thereof (no known vector [NKV]).

However, even this rather dramatic reorganization can only be seen as an initial step for modernizing flavivirid taxonomy because numerous viruses have been described that await classification, several of which likely require the establishment of novel genera (2). Here, we focus on potential new species in the subgenera *Fusivirus* and *Euflavivirus* (Table S2).

Despite the rapid pace of virus discovery and description of viruses phylogenetically related to orthoflaviviruses, the criteria used for species-rank classification and the viruses assigned have remained almost unchanged for more than two decades; the ICTV 8^th^ Report from 2005 lists 53 of the current 55 species assigned to the *Euflavivirus* subgenus (1). There has similarly been no update over this period on the criteria used to assign orthoflavivirus species, which are currently based on a polythetic list of properties (nucleotide and deduced amino acid sequence, antigenic characteristics, geographic association, vector association, host association, disease association, and ecological characteristics) without specifying which are essential for classification and which are descriptive. Evolutionary relatedness and metrics of sequence divergence have not been the primary metric for species assignment, with examples provided of genetically similar viruses being assigned to different species because of different ecological distributions (*eg.*, louping ill virus [LIV] distributed in moorlands of the UK, and “tick-borne encephalitis virus [“TBEV”] dwelling in forests of mainland Europe). Conversely, the phylogenetically distinct and genetically divergent DENV-1–4 have all been assigned to a single species; despite their documented antigenic differences (5), they share geographical and ecological habitats, transmission vectors and display similar disease manifestations.

In the current era, when vast numbers of novel viruses are being discovered primarily by high throughput metagenomic screening of environmental and clinical samples, the International Committee on Taxonomy of Viruses (ICTV) has enabled virus taxonomic assignments to be based on nucleotide sequences alone, provided the sequences of viral genomes contain complete coding sequences and are of adequate quality (6, 7). In the absence of information on their phenotypic properties, assignments of these virus sequences at ranks such as order, family, genus, and species can largely or entirely be guided by metrics of sequence relatedness in the absence of information on their phenotypic properties. Indeed, the recent consensus that virus taxonomy should be guided by and be consistent with virus evolutionary relationships (8) places sequence relationships as the primary criteria for virus taxonomic assignments. For newly discovered viruses, these have taken priority over phenotypic features such as disease associations, geographical, ecological distributions, and antigenicity that served as defining criteria in many of the older polythetic species definitions.

In case of *Flaviviridae*, no new species have been established for over two decades (apart from two new assignments of TBEV-related species in 2025), and the ICTV Report has reproduced similar standardised lists of unclassified flavivirids over its many editions in this period. Despite this, many of the 37 unclassified viruses listed in the 10^th^ Report (Table S2) have been extensively described in the virology literature, including Kamiti River virus (*n=*23 publications over the period 2003–2026), Culex flavivirus (*n*=69, 2007–2026), Quảng Bình virus (QBV; *n*=12, 2009–2026), Spondweni virus (SPOV; *n*=17, 1982–2025), Cháoyáng virus (CHAOV; *n*=9, 2013– 2023) and many others. That these unclassified orthoflaviviruses are not obscure or insignificant is attested by being the subject of at least 190 PubMed-indexed papers (Table S3).

Because there are often quite limited phenotypic descriptions of many of the unclassified orthoflaviviruses, it would be problematic to apply the current, polythetic species assignment criteria listed by the ICTV for their taxonomic assignments. To address this problem, we have re-investigated whether a genetic distance threshold might be additionally or alternatively used for species demarcation, as was previously attempted using *NS5* sequences from a wide collection of flavivirus isolates (9) and more widely applied for the classification of other vertebrate and invertebrate viruses. In the current study, we have selected a pairwise distance threshold for *NS5* that largely assigns species parsimoniously with current assignments and would enables over 50 new species to be established.

## Methods

### Orthoflavivirus sequence dataset

The current classification of members of the family *Flaviviridae* was drawn from the ICTV Master Species List (1). This provides a listing of the 55 established species in the subgenus *Euflavivirus* and single established species in the subgenera *Fusivirus* and *Crangovirus* (Table S1).

Orthoflavivirus genome sequences (minimum length 9,000 bases, annotated as coding-complete) were available from most of the currently classified members of the *Orthoflavivirus* genus (Table S1). In addition to the exemplar virus, a random selection of up to 19 additional isolates of each virus, if available, were included in the dataset (Table S2).

Unclassified likely orthoflaviviruses were identified through BLAST searching of GenBank for virus genome sequences related to the founding members of the *Fusivirus* and *Crangovirus* genera, cell-fusing agent virus (CFAV; M91671) and Crangon crangon flavivirus (CCFV; MK473878). A further set of likely orthoflaviviruses (taxid: 3076163) were downloaded from NCBI Taxonomy and filtered to include genome sequences >7,000 bases in length. All analysed sequences were downloaded from 12–14 August 2026. Additional searches for unclassified divergent likely orthoflaviviruses, *i.e.*, Cnidaria flavivirus (CnFV; OX394137), Hángzhōu flavivirus 3 (HzFV3; MZ209680), Harrimaniidae flavivirus (HarFV; OX394149), sea-firefly flavivirus (OX394161) and Tabanus rufidens flavivirus (TrFV; LC540441) were performed. However, as our search strategy was primarily focussed on retrieval of virus sequences from the *Euflavivirus* and *Fusivirus* genera, a small number of divergent invertebrate marine flavi-like viruses (10) were excluded.

### Sequence comparisons

Sequence and gene annotations, alignment and estimation of sequence distances (nucleotide and translated amino acid p-distances) were performed using SSE sequence editor (11) and embedded programs (*e.g.*, MUSCLE; (12)). Phylogenetic analyses were performed using MEGA version 7 (13). Distribution of sequence distances were analysed by SPSS v.29. Genome positions and protein boundaries were numbered as annotated in the DENV-2 reference sequence M19197. FASTA format alignments of euflavivirus and fusivirus genome sequences used for analysis are provided in the Supplemental Materials.

## Results

### *Orthoflavivirus* sequence diversity

An alignment was created of all orthoflavivirus genome sequences for analysis of their genetic diversity. This alignment represented 52 of the 57 classified orthoflaviviruses for which at least coding-complete genome sequences were available (Table S1) and for all unclassified viruses that grouped phylogenetically within the *Orthoflavivirus* genus. This dataset comprised primarily fusiviruses and euflaviviruses (*n*=171 and *n*=554, respectively) but also included sequences of crangoviruses (*n*=3). Phylogenetic analysis of *NS5* sequences (Fig. 1; a labelled tree with bootstrap support values is provided in Fig. S1) revealed a large clade of classified euflaviviruses with a separate subgrouping of tick-vectored flaviviruses from those transmitted by mosquitoes or with NKV. The *Euflavivirus* clade additionally harboured several often well characterised, but currently unclassified, viruses such as Lammi virus (LAMV) and Barkédji virus (BJV or BARKV). The *Fusivirus* clade harboured CFAV as the only taxonomically assigned virus (*Orthoflavivirus junctionis*) and sequences of many currently unclassified viruses forming additional species-level groupings. Finally, the *Crangovirus* clade, harboured CCFV while deeper branches in the tree were occupied by other highly divergent orthoflaviviruses, including the unclassified CnFV, HzFV3, sea-firefly virus, and HarFV. They all grouped separately from the three established orthoflavivirus groups and therefore may represent additional subgenera.

**Fig. 1.**
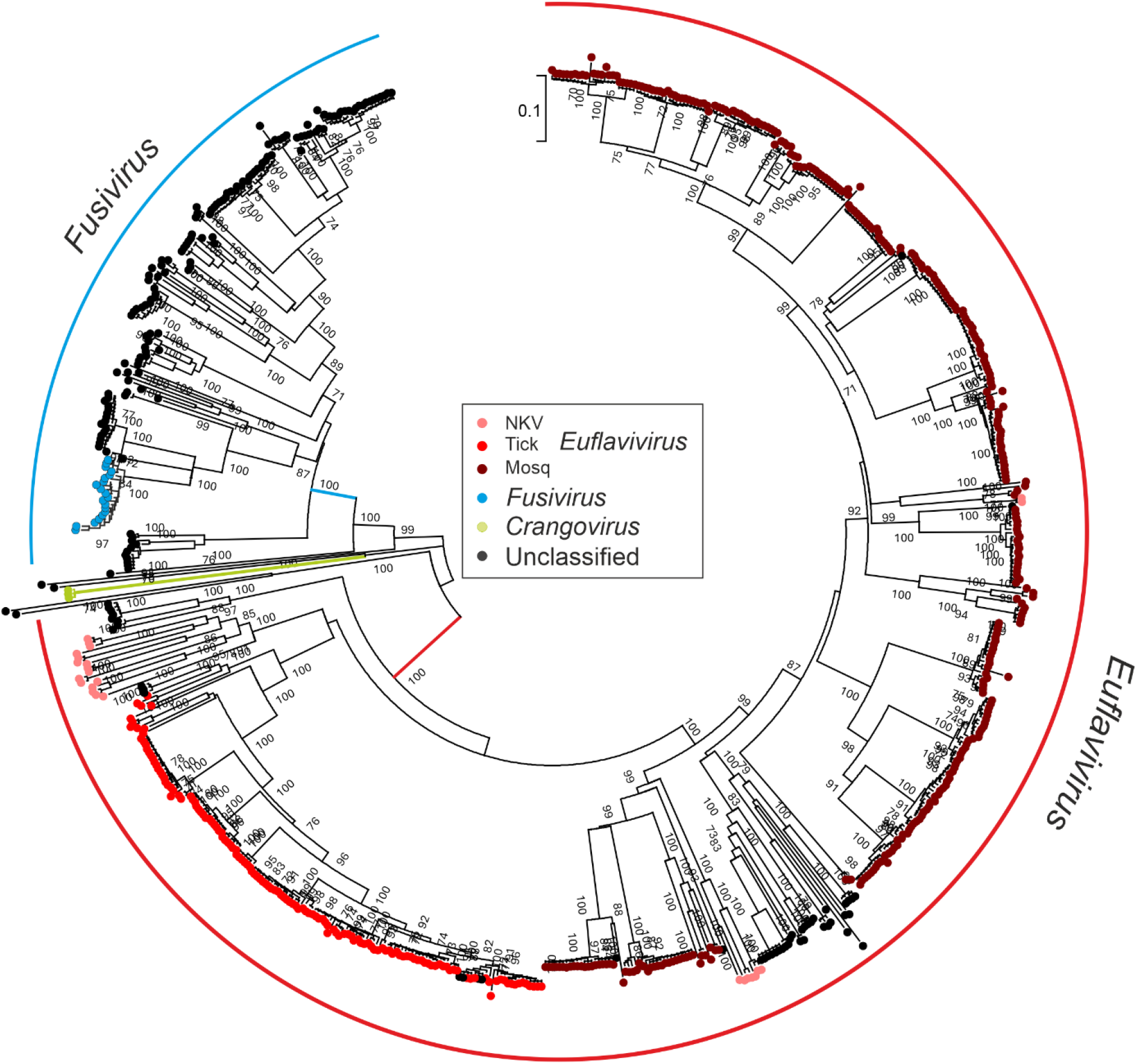
Phylogenetic tree of *NS5* sequences of classified euflaviviruses, fusiviruses (CFAV; *Orthoflavivirus functionis*) and crangon crangon virus (*Crangovirus; Orthoflavivirus aphei*) and a range of currently unclassified orthoflaviviruses. Euflaviviruses were sub-subdivided based on vector use; NKV: no known vector; Tick and Mosq.: mosquito. Bootstrap support (100 replicates of 70% or greater shown on branches). Coloured branches display proposed lineages for the three classified subgenera. A labelled tree is provided as Fig. S1 (Suppl. Data).

### Species rank classification

To investigate whether the current classification of orthoflaviviruses is supported by sequence relationships, sequence alignments of currently classified viruses in the *Euflavivirus* subgenus, available sequences of CFAV and CCFV were retrieved and the distributions of within- and between species genetic divergence compared (Fig. 2; Table 1). Pairwise sequence distances between the 504 classified orthoflavivirus sequences were concordant between *E* and *NS5* nucleotide and deduced *E* and *NS5* amino acid sequences (Fig. 2), although divergence in the latter region was systematically less than observed for *E*. The marked concordance in distances rules out recombination or mosaicism in euflavivirus genomes. For the purposes of making species assignments, sequence distances for the orthoflavivirus polyprotein (*PP*), and for *NS5* and *E* genes should all be equivalent even though the exact demarcation thresholds separating inter-species and intra-species distances may differ between genome regions.

**Fig. 2.**
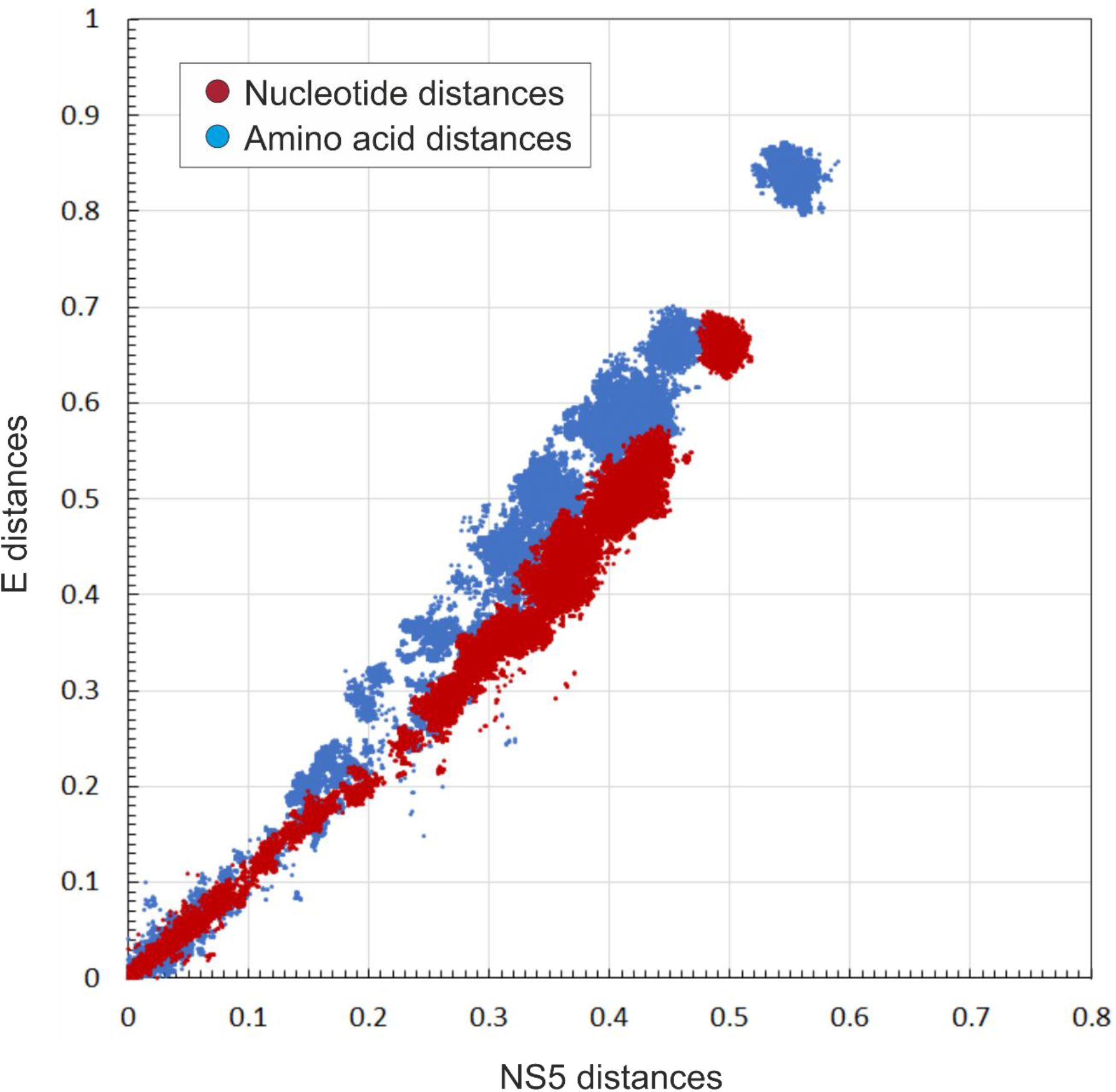
Comparison of nucleotide and translated amino acid pairwise distances of classified orthoflaviviruses (*n*=707; 130,136 datapoints) in *NS5* and *E*.

**Table 1.** Species with p-distances of amino acid (aa) sequences below the proposed demarcation thresholds for the whole virus polyprotein (*PP*), *NS5*, and *E*.

| Species 1 | Species 2 | Count<sup>1</sup> | PP aa distance<sup>2</sup> | NS5 aa distance<sup>2,3</sup> | E aa distance<sup>2,3</sup> |
| --- | --- | --- | --- | --- | --- |
| ***Ntaya virus group*** |  |  |  |  |  |
| *Orthoflavivirus israelense* | *Orthoflavivirus bagazaense* | 114 | 0.015 (0.006–0.030) | 0.013 (0.000–0.039) | 0.016 (0.000–0.086) |
|  | *Orthoflavivirus ntayaense* | 12 | 0.116 (0.114–0.118) | 0.095 (0.095–0.095) | 0.082 (0.074–0.090) |
| *Orthoflavivirus ntayaense* | *Orthoflavivirus bagazaense* | 36 | 0.118 (0.112–0.131) | 0.096 (0.108–0.108) | 0.082 (0.074–0.102) |
| ***Tick-borne encephalitis virus group*** |  |  |  |  |  |
| *Orthoflavivirus neudoerflense* | *Orthoflavivirus loupungi* | 400 | 0.050 (0.000–0.060) | 0.038 (0.000–0.064) | 0.059 (0.000–0.073) |
|  | *Orthoflavivirus mediterraneanense* | 40 | 0.062 (0.058–0.066) | 0.049 (0.042–0.054) | 0.042 (0.036–0.050) |
|  | *Orthoflavivirus zilberi* | 400 | 0.066 (0.061–0.073) | 0.062 (0.054–0.074) | 0.043 (0.034–0.059) |
|  | *Orthoflavivirus omskense* | 400 | 0.102 (0.098–0.106) | 0.075 (0.070–0.084) | 0.075 (0.067–0.083) |
| *Orthoflavivirus zilberi* | *Orthoflavivirus mediterraneanense* | 40 | 0.080 (0.067–0.086) | 0.064 (0.042–0.075) | 0.069 (0.059–0.081) |
|  | *Orthoflavivirus loupungi* | 400 | 0.080 (0.061–0.092) | 0.064 (0.055–0.099) | 0.084 (0.034–0.103) |
|  | *Orthoflavivirus omskense* | 400 | 0.102 (0.098–0.108) | 0.083 (0.075–0.095) | 0.073 (0.059–0.087) |
| *Orthoflavivirus mediterraneanense* | *Orthoflavivirus loupungi* | 40 | 0.075 (0.060–0.086) | 0.057 (0.045–0.088) | 0.063 (0.044–0.069) |
|  | *Orthoflavivirus omskense* | 40 | 0.112 (0.110–0.114) | 0.084 (0.075–0.085) | 0.092 (0.083–0.097) |
| *Orthoflavivirus loupungi* | *Orthoflavivirus omskense* | 400 | 0.115 (0.100–0.125) | 0.084 (0.074–0.106) | 0.107 (0.067–0.117) |
| ***Others*** |  |  |  |  |  |
| *Orthoflavivirus nilense* | *Orthoflavivirus koutangoense* | 80 | 0.109 (0.106–0.114) | 0.094 (0.103–0.103) | 0.120 (0.118–0.120) |
| *Orthoflavivirus wesselsbronense* | *Orthoflavivirus sepikense* | 20 | 0.131 (0.128–0.133) | 0.140 (0.144–0.144) | 0.086 (0.082–0.090) |

<sup>1</sup>Total number of pairwise comparisons
<sup>2</sup>Mean (minimum–maximum) p values.
<sup>3</sup>Grey shaded boxes contain p values greater than the *NS5* or *E* threshold values (0.10).

To determine potential demarcation thresholds for species assignments, distributions of amino acid sequence p-distances of *NS5* (Fig. 3), *E*, and *PP* (Fig. S2) were calculated with inter- and intra-species distances plotted separately. Distributions of distances were comparable although with different scaling reflecting differing degrees of sequence divergence between genome regions. Inter-species distances were substantially greater than intra-species distances for most orthoflaviviruses, although ranges were overlapping and no single threshold value would consistently assign species according to their current classification. However, the proposed thresholds of 0.1 (*NS5*—red dotted line), 0.1 (*E*), and 0.14 (polyprotein) were parsimonious with current assignments. Contrastingly, an *NS5* distance threshold of 0.2 corresponding to the second dip in the distance distribution (Fig. 3) would merge a large number of orthoflaviviruses currently classified into different species (blue bars in same species distribution in Fig. 3).

**Fig. 3.**
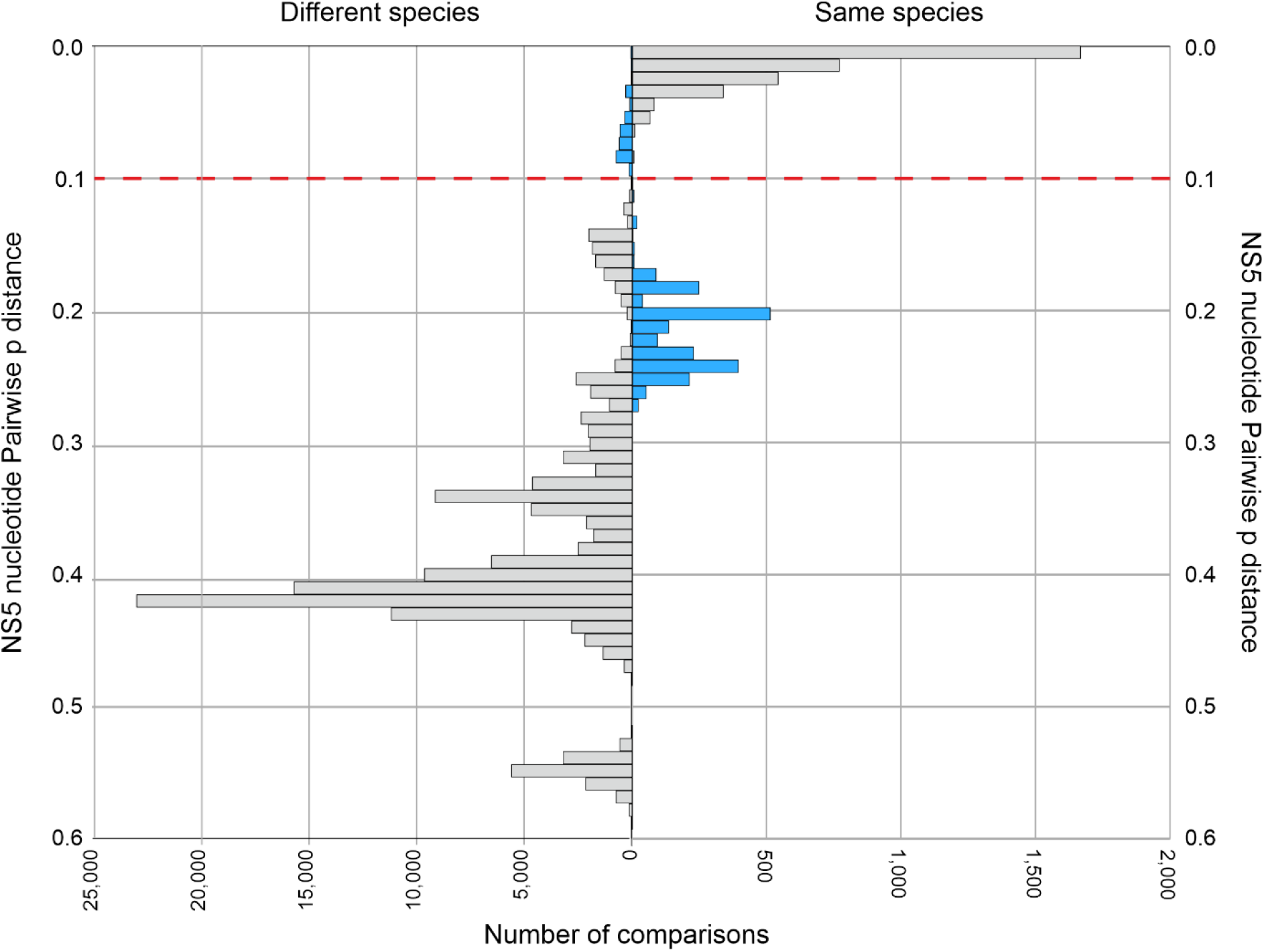
Distributions of *NS5* amino acid p-distances between (left) and within (right) species of classified orthoflaviviruses. Based on a tentative assignment threshold of 0.10 (red dashed line), assignments when sequence distances between species were below or within-species distances above the threshold are shown in blue (listed in Tables 1 and 2 respectively). Values in the range of 0.53–0.57 are from sequence comparisons of cell-fusing agent virus (*Orthflavivirus junctionis*) with viruses assigned to subgenus *Euflavivirus*.

**Table 2.** *NS5* amino acid (aa) sequence p-distances between different orthoflaviviruses.

| Species | Virus 1 | Virus 2 | Count<sup>1</sup> | NS5 aa distance<sup>2,3</sup> |
| --- | --- | --- | --- | --- |
| *Orthoflavivirus aroaense* | Aroa virus | Iguape virus | 4 | 0.087 (0.084–0.090) |
|  |  | Bussuquara virus | 2 | 0.277 (0.276–0.277) |
|  |  | Naranjal virus | 2 | 0.274 (0.274–0.275) |
|  | Naranjal virus | Bussuquara virus | 1 | 0.102 (0.102–0.102) |
|  |  | Iguape virus | 2 | 0.271 (0.268–0.275) |
|  | Iguape virus | Bussuquara virus | 2 | 0.269 (0.266–0.272) |
| *Orthoflavivirus denguei* | dengue virus type 3 | dengue virus type 1 | 400 | 0.184 (0.176–0.191) |
|  |  | dengue virus type 2 | 400 | 0.209 (0.201–0.219) |
|  |  | dengue virus type 4 | 400 | 0.232 (0.226–0.252) |
|  | dengue virus type 2 | dengue virus type 1 | 400 | 0.205 (0.198–0.211) |
|  |  | dengue virus type 4 | 400 | 0.252 (0.243–0.275) |
|  | dengue virus type 4 | dengue virus type 1 | 400 | 0.242 (0.238–0.262) |
| *Orthoflavivirus kokoberaorum* | Kokobera virus | Bainyik virus | 2 | 0.083 (0.081–0.086) |
|  |  | New Mapoon virus | 2 | 0.251 (0.250–0.251) |
|  |  | Stratford virus | 10 | 0.159 (0.156–0.163) |
|  |  | Torres virus | 2 | 0.137 (0.136–0.138) |
|  | Stratford virus | Torres virus | 5 | 0.113 (0.112–0.115) |
|  |  | Bainyik virus | 5 | 0.153 (0.153–0.153) |
|  |  | New Mapoon | 5 | 0.239 (0.237–0.241) |
|  | Torres virus | Bainyik | 1 | 0.120 (0.120–0.120) |
|  |  | New Mapoon | 1 | 0.239 (0.239–0.239) |
|  | New Mapoon virus | Bainyik virus | 1 | 0.241 (0.241–0.241) |
| *Orthoflavivirus* | Sokuluk virus | Entebbe bat virus | 6 | 0.177 (0.175–0.180) |
| *Orthoflavivirus* | Rocio virus | Ilhéus virus | 36 | 0.195 (0.191–0.201) |
| *Orthoflavivirus* | Kyasanur Forest | Alkhumra | 84 | 0.024 (0.021–0.027) |
| *Orthoflavivirus* | Murray Valley | Alfuy virus | 19 | 0.136 (0.133–0.139) |
| *Orthoflavivirus* | Phnom Penh bat | Batu Cave virus | 2 | 0.023 (0.022–0.024) |
| *Orthoflavivirus* | Powassan virus | deer tick virus | 36 | 0.008 (0.001–0.038) |
| *Orthoflavivirus royalense* | Royal Farm virus | Karshi virus | 3 | 0.183 (0.181–0.187) |
| *Orthoflavivirus* | Saboya virus | Potiskum virus | 1 | 0.003 (0.003–0.003) |

<sup>1</sup>Total number of pairwise comparisons<sup>2</sup>Mean (minimum–maximum) p values.<sup>3</sup>Grey shaded boxes contain p values greater than the NS5 threshold value (0.10).

However, several inter-species comparisons had mean pairwise distances below each of the three thresholds, including the tick-borne orthoflaviviruses assigned to *Orthoflavivirus zilberi* (tick-borne encephalitis virus 1 [TBEV-1]), *Orthoflavivirus loupungi* (LIV), *Orthoflavivirus neudoerflense* (TBEV-2), and *Orthoflavivirus omskense* (Omks hemorrhagic fever virus [OHFV]) and viruses in the “Ntaya goup”, *i.e.*, Ntaya virus (NTAV; *Orthoflavivirus ntayaense*), Bagaza virus (BAGV; *Orthoflavivirus bagazaense*), and Israel turkey meningoencephalomyelitis virus (ITV; *Orthoflavivirus israelense*) (Table 1). Pairwise distances between WNV (*Orthoflavivirus nilense*) and Koutango virus (KOUV; *Orthoflavivirus koutangoense*), and between Wesselsbron virus (WESSV; *Orthoflavivirus wesselsbronense*) and Sepik virus (SEPV; *Orthoflavivirus sepikense*) were close to (either above or below) the assignment thresholds in the three regions. All other pairwise distances between species were substantially above the thresholds indicated.

The majority of named orthoflaviviruses showed within-species pairwise distances below the proposed thresholds for all three genome regions (Table S4), the only exception being Yokose virus (YOKV) with the two characterised variants having pairwise distances of 0.180, 0.143, and 0.259 for *PP*, *NS5*, and *E* genome regions respectively.

However, p-distances above proposed thresholds for within-species distances (blue shaded bars in the right-hand histogram in Fig. 3 and Fig. S2) indicate that there were several discrepancies with current species assignments (Table 2). Most prominently, DENV types 1-4 were highly divergent in sequence from each other, well above the intra-species divergence range for *NS5* (0.177–0.256; Table S2; Suppl. Data) and *E* and complete polyprotein sequences (0.215–0.352 and 0.214–0.306 respectively). Mean within-DENV type *NS5* distances ranged from 0.015–0.024 (Table S4) although these were based on the random selection of 20 human-derived variants for each type. Indeed, specifically selected sylvatic isolates of DENV-1, -2 and -4 (14, 15) exhibited greater sequence divergence than human DENV isolates, with distances from homologous types ranging from 0.015 to 0.081 (Table S5), although remaining below the 0.1 threshold. However, DENV isolates previously identified as being divergent such as DKE-121 (16) and QMS-22 (17) had pairwise distances in *NS5* and other regions slightly above the proposed species threshold.

High sequence divergence within a single species is also observed among most viruses assigned to *Orthoflavivirus aroaense* (Table 2; Table S6) with sequences from Iguape virus (IGUV), Bussuquara virus (BSQV) and Naranjal virus (NJLV) having >25% sequence divergence in *NS5* (threshold 0.1), with only Aroa virus (AROAV) and IGUV showing sequence distance consistent with same-species assignment (mean 0.087). Apart from a same-species *NS5* divergence of 0.087 between Kokobera virus (KOKV) and Bainyik virus, all other viruses assigned to *Orthoflavivirus kokoberaorum* (New Mapoon virus [NMV], Stratford virus [STRV] and Torres virus) had *NS5* distances in the inter-species range (0.113–0.51). Table 2 lists four further viruses having sequence divergence above the threshold for species assignment.

### Sequence divergence of currently unclassified orthoflaviviruses

The majority of euflaviviruses have long-established species assignments, but 50 further currently unclassified viruses grouped within the *Euflavivirus* clade while 142 grouped with CFAV in the *Fusivirus* clade (Fig. 1; Fig. S1, Suppl. Data). To establish whether these viruses could be assigned to established or novel species using the thresholds established for the current classified dataset, pairwise distances for sequences were plotted for the *NS5* and *E* regions (Fig. 4). Inclusion of unclassified, likely euflaviviruses produced sequence distance distributions in the *NS5* region that were comparable, although more restricted in range, to the comparison of all classified orthoflaviviruses (Figs. 2, 4). Contrastingly, the set of 159 fusivirus sequences produced inconsistent sequence relationships between *E* and *NS5* specifically for comparisons involving CFAV. Phylogenetic trees for *NS5* and *E* (Fig. 5) showed that the clade of CFAV changed position between the two genome regions, clustering most closely with Aedes flavivirus in the *NS5* tree but with QBV and Culex theileri virus (CTFV) in *E*. Separate comparison of trees for other genes produced topologies similar to *NS5* for all the non-structural genes (*NS1*–*NS5*), while the *pre-M* and *M* genes resembled *E* topology in sequence relationships (data not shown).

**Fig. 4.**
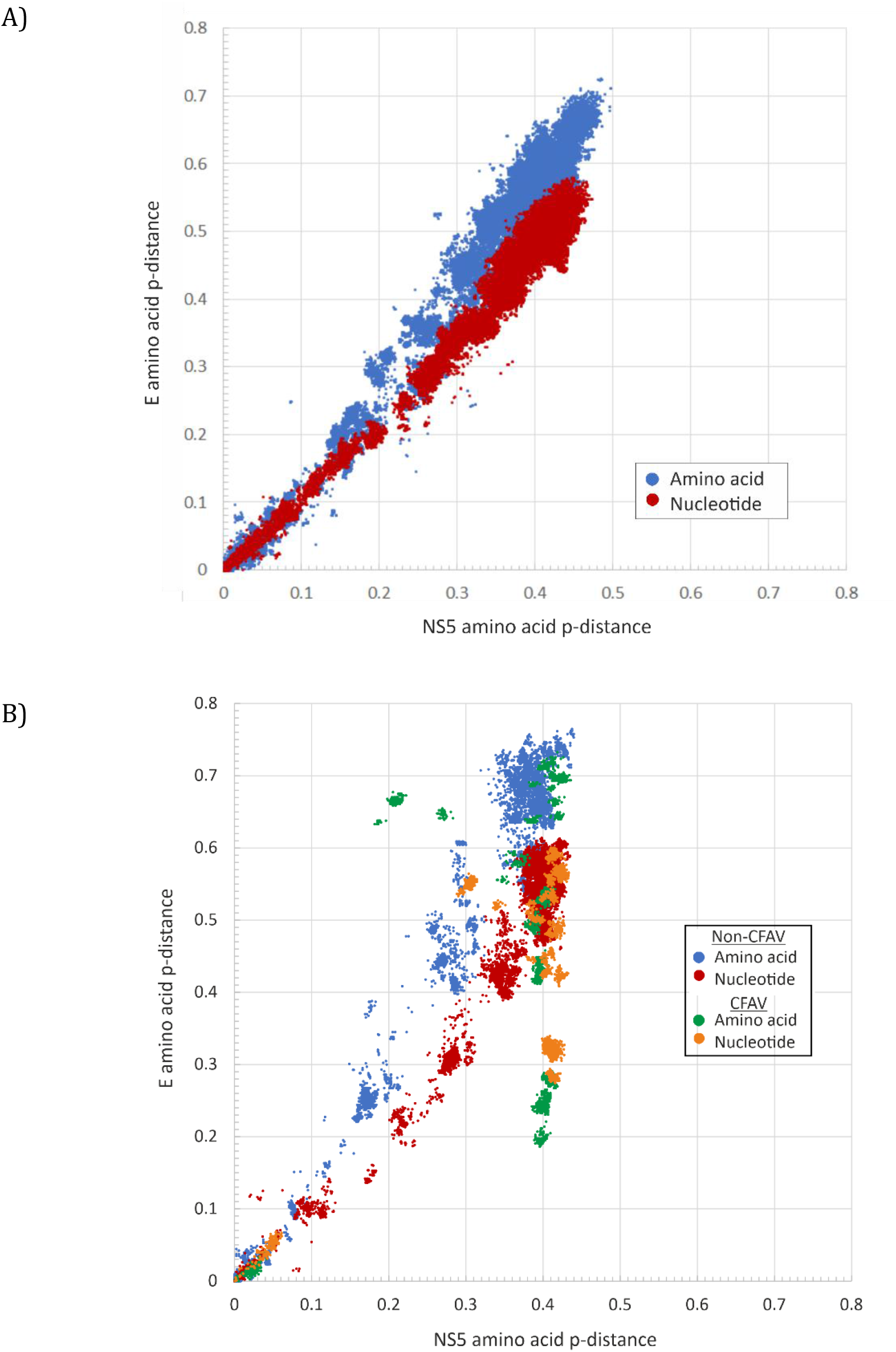
Pairwise nucleotide and amino acid sequence distances in *NS5* and *E* of (A) euflaviviruses and (B) fusiviruses, respectively. Pairwise comparisons involving cell-fusing agent virus (CFAV; *Orthoflavivirus junctionis*) are shaded separately.

**Fig. 5.**
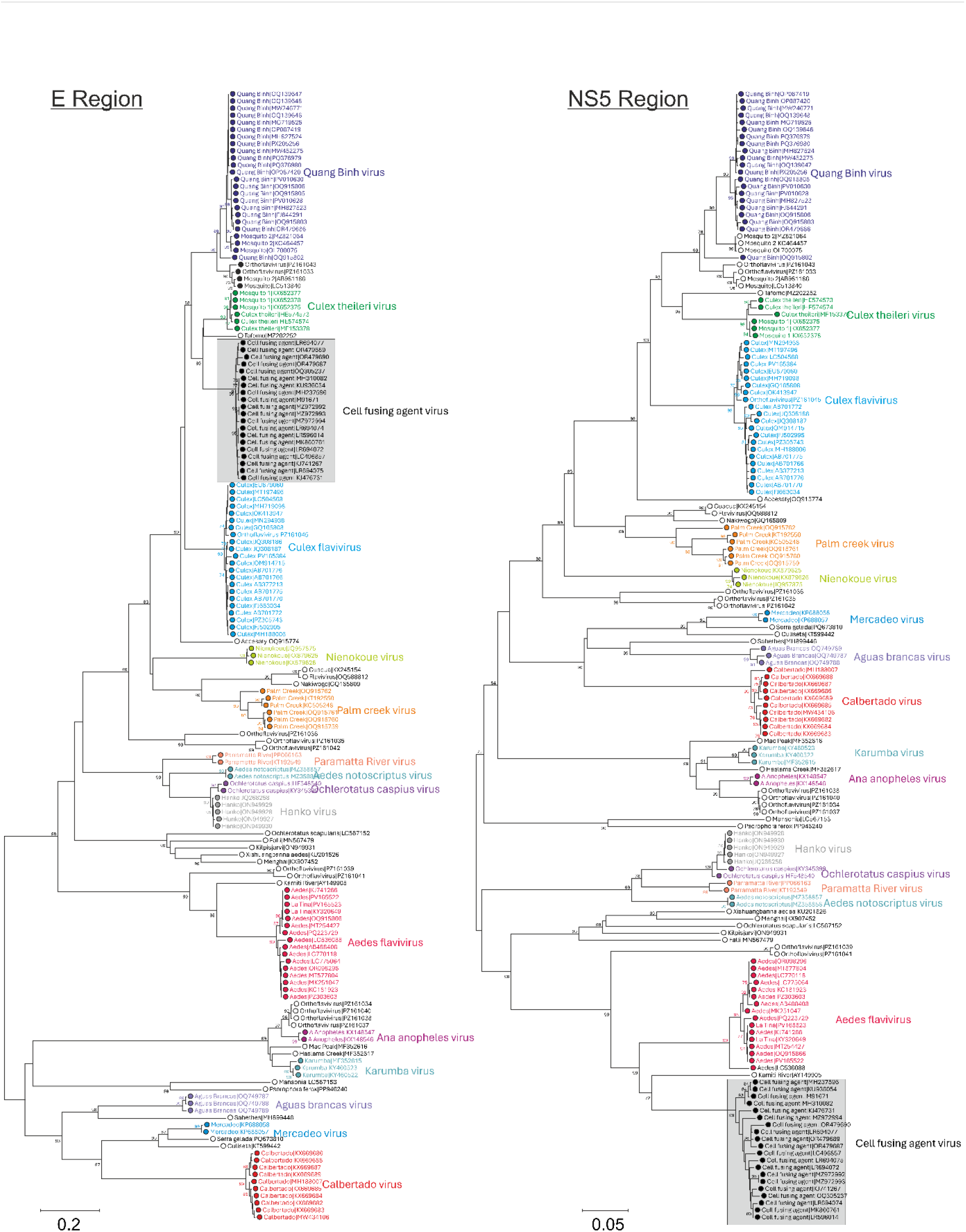
Phylogeny of *E* and *NS5* amino acid sequences of fusiviruses based on neighbour-joining using the JTT substitution model; branches supported by 70% bootstrap replicates are indicated. Virus-level clades are coloured to assist comparison of phylogenies.

### Proposals for the classification of currently unassigned orthoflaviviruses

We propose to use a sequence divergence threshold of 0.1 for *NS5* amino acid sequences to assign currently unclassified orthoflaviviruses. While the *E* gene and the whole polyprotein coding sequence split unclassified and classified orthoflaviviruses equivalently using similar thresholds, the mosaicism of fusiviruses (Figs. 4, 5) indicates that reliance on a single representative gene may be preferable for primary assignments (although it would be important to examine comparable relationships in other genome regions). As the longest gene in the *Orthoflavivirus* genome, and one that contains the fewest indels that might hamper alignment and identification of homologous sequences, *NS5* is the region best suited for species rank classification.

The distribution of sequence distances of the whole set of classified and unclassified euflaviviruses was comparable to classified viruses alone (Fig. 6 compared to Fig. 2). The distribution of fusivirus p-distances was comparable to that of euflaviviruses, featuring pronounced peaks near 0.4 and minor peaks at 0.3 and between 0.11 and 0.2. Minimum values occurred between 0.10 and 0.11, below which the sequence distances corresponded to the typical intra-species distances of classified euflaviviruses. Broadly, the proposed 0.1 amino acid sequence p-distance threshold established for classified euflaviviruses appeared appropriate for fusiviruses as well.

**Fig. 6.**
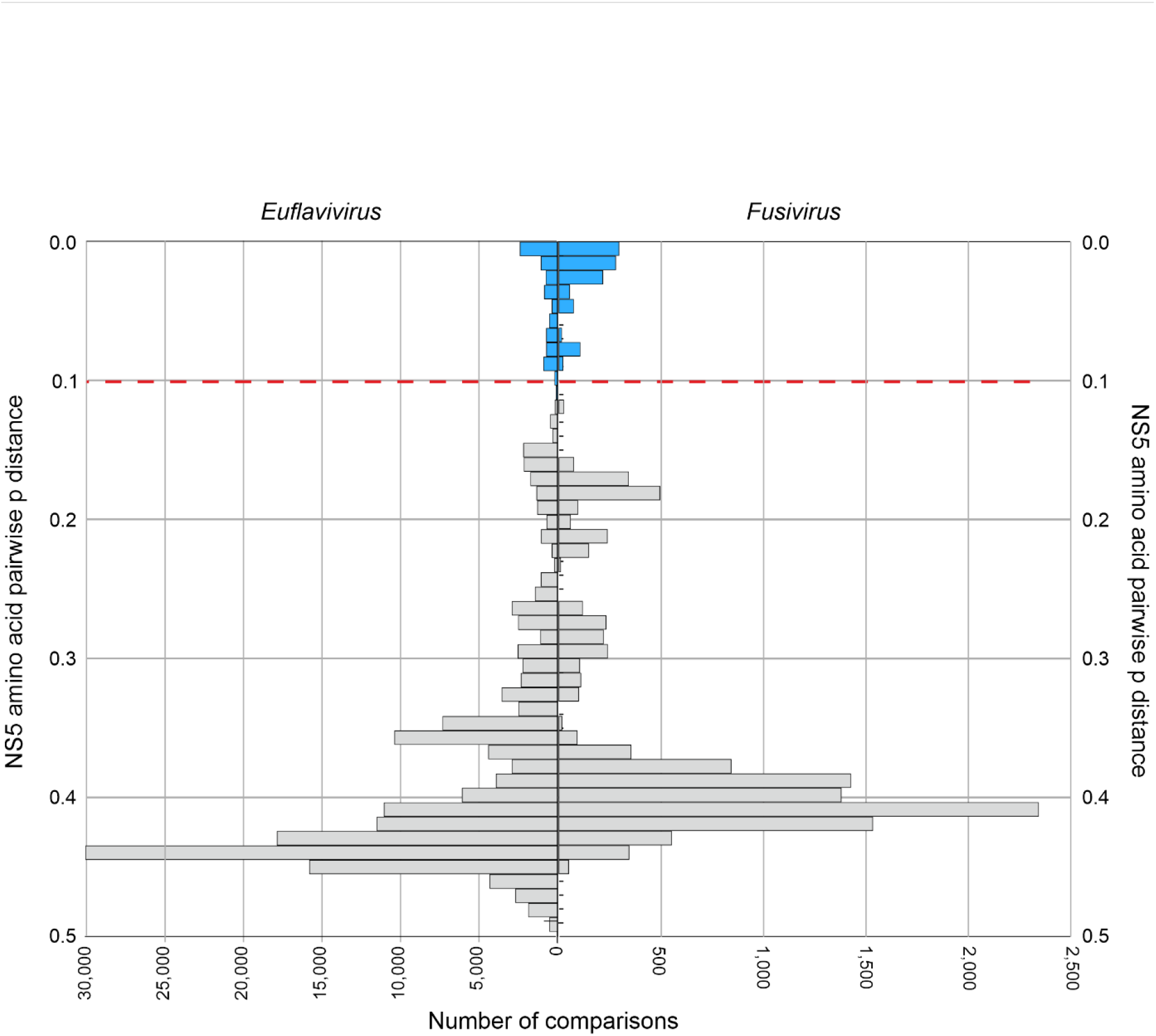
Comparison of pairwise *NS5* amino acid sequence distance distributions of classified and unclassified likely euflaviviruses and fusiviruses. Blue shaded bars depict pairwise distances below the *NS5* species threshold of 0.10.

As a guide towards the future classification of euflaviviruses based on metrics of genetic divergence, mean *NS5* p-distances were calculated for 27 currently unclassified euflaviviruses in the dataset against the set of classified members (Table 3). The closest mean distances associated with classified species were almost invariably in the interspecies range. The only exceptions were Fitzroy River virus (FRV) and WESSV with a divergence equal to the threshold value. Overall, the analysis supports the establishment of at least 24 new euflavivirus species.

**Table 3.** Closest matches of currently unassigned euflaviviruses to classified and unclassified viruses.

| Unclassified virus | Closest classified virus | Species of closest classified virus | NS5 aa p-distance<sup>1</sup> |
| --- | --- | --- | --- |
| Bamaga virus | Banzi virus | *Orthoflavivirus banziense* | 0.309 |
| Barkédji virus | Koutango virus | *Orthoflavivirus koutangoense* | 0.364 (0.363–0.365) |
| Binjari virus | Bussuquara virus | *Orthoflavivirus aroaense* | 0.361 |
| Cháoyáng virus | Bussuquara virus | *Orthoflavivirus aroaense* | 0.357 |
| Chinese soft-shelled turtle flavivirus | Naranjal virus | *Orthoflavivirus aroaense* | 0.378 |
| Dōnggǎng virus | Ilhéus virus | *Orthoflavivirus ilheusense* | 0.352 (0.352–0.356) |
| Fitzroy River virus | Wesselsbron virus | *Orthoflavivirus wesselsbronense* | 0.105 (0.102–0.108) |
| Héběi orthoflavivirus | Yaoundé virus | *Orthoflavivirus yaoundeense* | 0.415 (0.413–0.419) |
| Ilomantsi virus | Usutu virus | *Orthoflavivirus usutuense* | 0.359 (0.358–0.363) |
| Kama virus | Meaban virus | *Orthoflavivirus meabanense* | 0.167 (0.167–0.168) |
| Kampung Karu virus | Usutu virus | *Orthoflavivirus usutuense* | 0.373 (0.372–0.376) |
| Lammi virus | Naranjal virus | *Orthoflavivirus aroaense* | 0.354 (0.354–0.354) |
| Long Pine virus | St. Louis encephalitis virus | *Orthoflavivirus louisense* | 0.356 (0.353–0.358) |
| Lunay virus | Yaoundé virus | *Orthoflavivirus yaoundeense* | 0.375 |
| Marisma mosquito virus | St. Louis encephalitis virus | *Orthoflavivirus louisense* | 0.348 (0.345–0.351) |
| Mpulungu flavivirus | West Nile virus | *Orthoflavivirus nilense* | 0.409 (0.405–0.417) |
| Nanay virus | Yaoundé virus | *Orthoflavivirus yaoundeense* | 0.369 |
| Nhumirim virus | Zika virus | *Orthoflavivirus zikaense* | 0.362 (0.359–0.367) |
| Nounané virus | Torres virus | *Orthoflavivirus kokoberaorum* | 0.387 |
| Paraiso Escondido virus | yellow fever virus | *Orthoflavivirus flavi* | 0.350 (0.344–0.356) |
| Psorophora flavivirus | Bussuquara virus | *Orthoflavivirus aroaense* | 0.362 (0.361–0.362) |
| Saruyama virus | West Nile virus | *Orthoflavivirus nilense* | 0.417 (0.413–0.420) |
| Spondweni virus | Zika virus | *Orthoflavivirus zikaense* | 0.226 (0.219–0.230) |
| T'hó virus | Ilhéus virus | *Orthoflavivirus ilheusense* | 0.212 (0.210–0.214) |
| Trionyx sinensis flavivirus | Naranjal virus | *Orthoflavivirus aroaense* | 0.380 (0.380–0.380) |
| Xīnyáng flavivirus | Yaoundé virus | *Orthoflavivirus yaoundeense* | 0.417 |
| Yanbian tick orthoflavivirus | Yaoundé virus | *Orthoflavivirus yaoundeense* | 0.413 |
| **Unclassified 1** | **Unclassified 2** |  | **NS5 aa p-distance** |
| Chinese soft-shelled turtle flavivirus | Trionyx sinensis flavivirus |  | 0.011 (0.010–0.011) |
| Lammi virus | Cháoyáng virus |  | 0.085 (0.085–0.086) |
| Héběi orthoflavivirus | Yanbian tick orthoflavivirus |  | 0.011 (0.009–0.012) |
| Marisma mosquito virus | Dōnggǎng virus |  | 0.104 (0.104–0.105) |
| Saruyama virus | Xīnyáng flavivirus |  | 0.006 |

<sup>1</sup>Mean (minimum–maximum) amino acid (aa) p values.

Sequence distances between fusiviruses were performed similarly (Table 4), although several viruses with uninformative names, such as “mosquito flavivirus” and “orthoflavivirus” that were often interspersed in clades of named viruses, were excluded (Fig. 5). Twenty-nine viruses exceeded the assigned minimum *NS5* p-distance threshold when compared to other species. An additional four virus pairs diverged by less than 0.1 from each other but exceeded this threshold against all other viruses, representing four further candidate species.

**Table 4.**
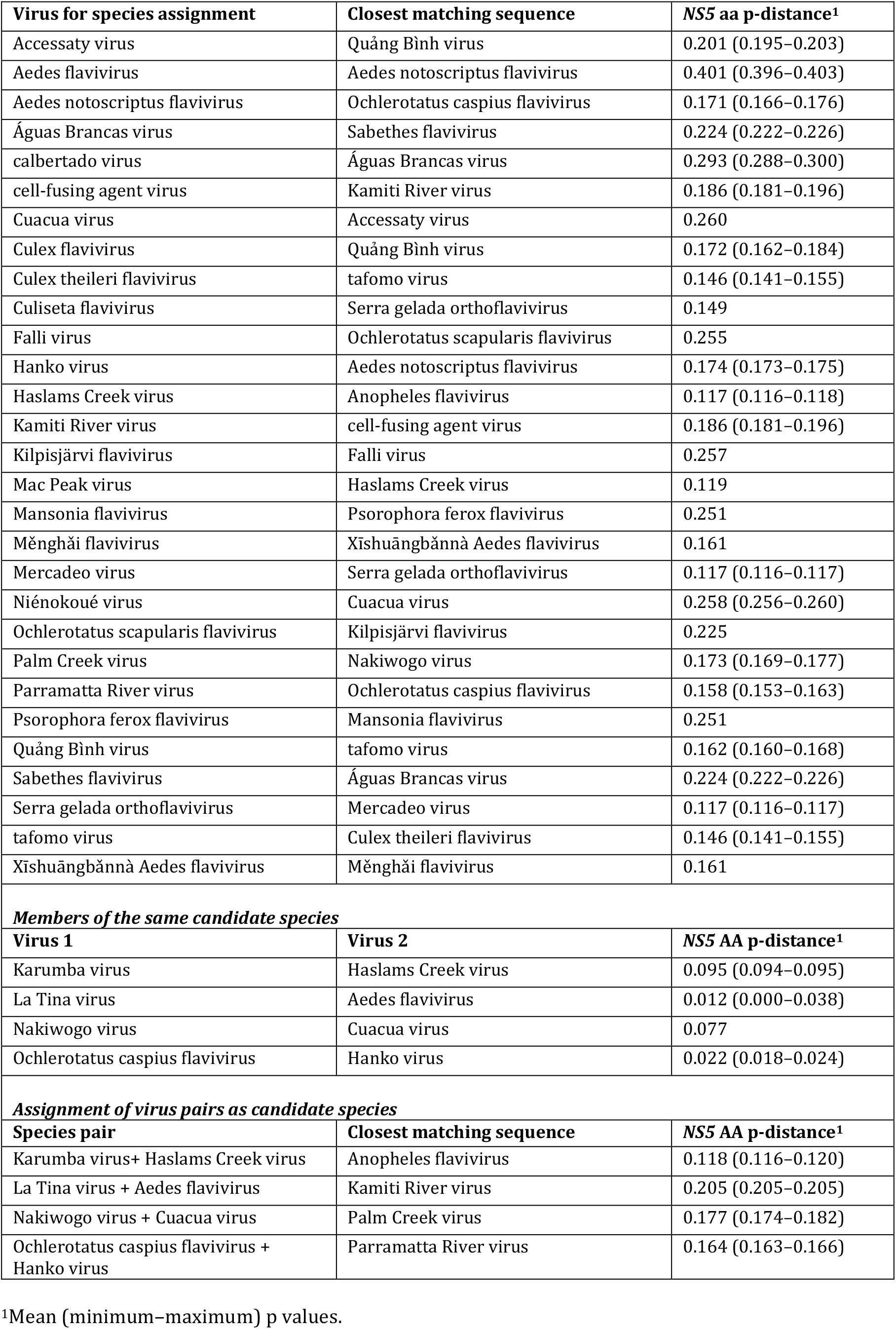
Classification of potential fusiviruses using the 0.10 *NS5* amino acid (aa) p-distance threshold.

| **Virus for species assignment** | **Closest matching sequence** | **NS5 aa p-distance<sup>1</sup>** |
| --- | --- | --- |
| Accessaty virus | Quảng Bình virus | 0.201 (0.195–0.203) |
| Aedes flavivirus | Aedes notoscriptus flavivirus | 0.401 (0.396–0.403) |
| Aedes notoscriptus flavivirus | Ochlerotatus caspius flavivirus | 0.171 (0.166–0.176) |
| Águas Brancas virus | Sabethes flavivirus | 0.224 (0.222–0.226) |
| calbertado virus | Águas Brancas virus | 0.293 (0.288–0.300) |
| cell-fusing agent virus | Kamiti River virus | 0.186 (0.181–0.196) |
| Cuacua virus | Accessaty virus | 0.260 |
| Culex flavivirus | Quảng Bình virus | 0.172 (0.162–0.184) |
| Culex theileri flavivirus | tafomo virus | 0.146 (0.141–0.155) |
| Culiseta flavivirus | Serra gelada orthoflavivirus | 0.149 |
| Falli virus | Ochlerotatus scapularis flavivirus | 0.255 |
| Hanko virus | Aedes notoscriptus flavivirus | 0.174 (0.173–0.175) |
| Haslams Creek virus | Anopheles flavivirus | 0.117 (0.116–0.118) |
| Kamiti River virus | cell-fusing agent virus | 0.186 (0.181–0.196) |
| Kilpisjärvi flavivirus | Falli virus | 0.257 |
| Mac Peak virus | Haslams Creek virus | 0.119 |
| Mansonia flavivirus | Psorophora ferox flavivirus | 0.251 |
| Mễnghải flavivirus | Xīshuāngbǎnnà Aedes flavivirus | 0.161 |
| Mercadeo virus | Serra gelada orthoflavivirus | 0.117 (0.116–0.117) |
| Niênokoué virus | Cuacua virus | 0.258 (0.256–0.260) |
| Ochlerotatus scapularis flavivirus | Kilpisjärvi flavivirus | 0.225 |
| Palm Creek virus | Nakiwogo virus | 0.173 (0.169–0.177) |
| Parramatta River virus | Ochlerotatus caspius flavivirus | 0.158 (0.153–0.163) |
| Psorophora ferox flavivirus | Mansonia flavivirus | 0.251 |
| Quảng Bình virus | tafomo virus | 0.162 (0.160–0.168) |
| Sabethes flavivirus | Águas Brancas virus | 0.224 (0.222–0.226) |
| Serra gelada orthoflavivirus | Mercadeo virus | 0.117 (0.116–0.117) |
| tafomo virus | Culex theileri flavivirus | 0.146 (0.141–0.155) |
| Xīshuāngbǎnnà Aedes flavivirus | Mễnghải flavivirus | 0.161 |
| **Members of the same candidate species** |  |  |
| **Virus 1** | **Virus 2** | **NS5 AA p-distance<sup>1</sup>** |
| Karumba virus | Haslams Creek virus | 0.095 (0.094–0.095) |
| La Tina virus | Aedes flavivirus | 0.012 (0.000–0.038) |
| Nakiwogo virus | Cuacua virus | 0.077 |
| Ochlerotatus caspius flavivirus | Hanko virus | 0.022 (0.018–0.024) |
| **Assignment of virus pairs as candidate species** |  |  |
| **Species pair** | **Closest matching sequence** | **NS5 AA p-distance<sup>1</sup>** |
| Karumba virus+ Haslams Creek virus | Anopheles flavivirus | 0.118 (0.116–0.120) |
| La Tina virus + Aedes flavivirus | Kamiti River virus | 0.205 (0.205–0.205) |
| Nakiwogo virus + Cuacua virus | Palm Creek virus | 0.177 (0.174–0.182) |
| Ochlerotatus caspius flavivirus + Hanko virus | Parramatta River virus | 0.164 (0.163–0.166) |

<sup>1</sup>Mean (minimum–maximum) p values.

## Discussion

### Classification of viruses into species

Species assignments of animals and plants have historically been designed to classify organisms at the most granular level possible, with species corresponding to sets of organisms with interbreeding capacity and shared gene pools. Assignments based on the biological species principle (or equivalents; (18)) obviously cannot apply to viruses (or prokaryotes) in the absence of sexual reproduction and in practice, species have been partitioned based on descriptive properties and patterns of genotypic clustering. For viruses, species assignments are increasingly based on metrics of genetic divergence. For example, enterovirus species are established based on a 30% amino acid sequence divergence threshold over the encoded polyprotein, or 40% in structural genes (19). Orthohepacivirus species are assigned based on a 30% amino acid sequence divergence threshold in *NS5B* (1), whereas most alphavirus species are demarcated at a 10% amino acid sequence threshold for whole genome sequences (20). The 10% amino acid p-distance (and equivalent demarcation of sequences using a 10% threshold for *E* sequences) is not dissimilar to proposed sequence-based species assignments in genera *Jingmenvirus* and *Guaicovirus*, where amino acid p-distance demarcation thresholds range from 12–17% for proteins encoded by their four genomic segments (21). The lowest threshold (12%) was for coding sequences in segment 1 that encode the jingmenvirus RNA-directed RNA polymerase, which is homologous to orthoflavivirus *NS5*. The adoption of a comparable genetic basis for the classification of orthoflaviviruses would therefore not be unprecedented.

### Use of genetic divergence for orthoflavivirus classification

The application of a fixed genetic threshold for the classification of orthoflaviviruses would seem to be advantageous for several reasons. Firstly, ways in which orthoflaviviruses species have been defined has varied over time, with highly variable species assignment criteria used to assign different orthoflaviviruses. Secondly, for vector-borne orthoflaviviruses, there is a secondary classification of serocomplexes based on antigenic cross-reactivity. Neither virus nor serocomplex groupings map naturally to current species assignments. Some serocomplexes (*eg.*, the “Aroa” and “Kokobera” groups) may contain multiple viruses and assigned species, whereas others may correspond to individual viruses (*eg.*, Zika virus of *Orthoflavivirus zikaense*, yellow fever virus of *Orthoflavivirus flavi*). Some species may be subdivided into a series of viruses with distinctive properties, such as STRV, Torres, NMV, and Bainyik viruses of *Orthoflavivirus kokoberaorum*. Finally, with often little information on the properties of insect-specific orthoflaviviruses and a primarily polythetic and descriptive species definitions for species assignments that are more relevant to vectored orthoflaviviruses, it is perhaps unsurprising that fusiviruses have remained almost entirely unclassified over the many decades in which they have been isolated and described.

The use of a fixed genetic divergence threshold based on *NS5* p-distance of 0.1 (and similarly for *E*), for establishment of orthoflavivirus species, would create a classification broadly consistent with current euflavivirus taxonomy. The *NS5* threshold for species identification is indeed analogous to the originally proposed 84% nucleotide sequence identity based on a partial highly conserved region of *NS5* (9). Most species assignments are supported by the proposed demarcation threshold, with a small number of exceptions listed in Table 1 (2 or more species would be re-classified as single species) and conversely, species with member viruses that were substantially more diverse than the threshold (Table 2).

However, many of these discrepancies are already widely recognised and therefore may not be controversial. Notably, ITV and BAGV are interspersed on phylogenetic analysis and minimally diverge from each other (mean 0.013, range 0.000–0.039), reflecting previous descriptions of these viruses as highly genetically related or identical (22, 23) or as proposed a merged species (24). NTAV would also fall into this species, consistent with the description of all three viruses as members of the “Ntaya” group. However, Tembusu virus (TMUV), another member of the group, shows an *NS5* divergence of 0.18 from other members, well above the species demarcation threshold.

The other group of species with divergence substantially below the *NS5* threshold are members of the “tick-borne encephalitis” group (LIV, TBEV-1, TBEV-2, and Turkish sheep encephalitis virus [TSEV],) along with OHFV, which are currently assigned to five separate species. Members of the previously assigned species *Orthoflavivirus encephalitidis* and *Orthoflavivirus loupungi* had different phenotypes (geographical and host range, transmission vector and pathogenicity) but recognition of *Orthoflavivirus loupungi* as a species made *Orthoflavivirus encephalitidis* paraphyletic (25) even though the two assigned species might be justified phenotypically given their systematic differences in geographical and host range and pathogenicities. The two species were recently formally reclassified into four monophyletic species by the ICTV (proposal 2025.007S; (26)). However, pairwise distances between these newly assigned species are all consistently below the proposed *NS5* demarcation threshold (Table 1). Two other pairs of virus species (*Orthoflavivirus nilense* [WNV]/[KOUV] and *Orthoflavivirus wesselsbronense* [Wesselbron virus]/*Orthoflavivirus sepikense* [Sepik virus]) had *NS5*, *E*, and *PP* divergence values close to the demarcation thresholds. Indeed, KOUV was formerly described as a lineage of WNV (27, 28), along with Kunjin virus and several further numbered (unnamed) lineages.

A larger number of discrepancies with the current species classification were observed for species with members that were far more divergent than the *NS5* threshold (Table 2). For example, the species *Orthoflavivirus aroaense* was first described as the “Aroa” group and formalised as the *Aroa virus* species in 2005 (29). While the assigned AROAV, BSQV, IGUV, and NJLV remain partially characterised (30–32), their description as a serocomplex indicates a degree of antibody cross-reactivity between them. Nevertheless, pairwise distances of >25% place BSQV and NJLV into two additional species. There would be a similar split of the species *Orthoflavivirus kokoberaorum*, with most members of the “Kokobera” group (NMV, STRV, Torres virus) being sufficiently divergent from KOV and from each other to constitute new species. Results for viruses of several other species (*Orthoflavivirus entebbeense*, *Orthoflavivirus ilheusense*, *Orthoflavivirus murrayense* and *Orthoflavivirus royalense*) indicate that some or all of these species might be similarly split.

Perhaps more strikingly, the four dengue virus types that are currently all assigned to *Orthoflavivirus denguei* had intertypic pairwise distances of 0.18–0.25, well above the range observed between other orthoflavivirus species. Dengue viruses were first characterised in the 1940s (33), long before the concept of a virus species was developed. Their subsequent designation as four types within a single species was based on their shared geographies, clinical presentations of dengue fever, and induction of partial cross-immunity (34), criteria that have been regarded as reasons for shared species assignments. However, their genetic diversity, matching that observed between different orthoflavivirus species, has been long recognised (5, 9, 35) even though the criteria for species assignment based on shared ecology, pathogenesis, and antigenic cross-reactivity using the current polythetic definition is quite incongruous with the obvious genetic divergence they display. The recent emergence of highly divergent DENV-2 and DENV-4 (16, 17) underscores their continuous evolution and challenges our current system of taxonomic classification. As these and several other highly divergent viruses fall outside the existing phylogenetic boundaries this suggests that the sylvatic dengue reservoir sampled to date represent only a limited fraction of the diversity circulating in nature. Any future reorganisation of *Orthoflavivirus denguei*, such as a potential split into four separate species, should not be controversial and would be unlikely to lead to nomenclatural confusion as the names of the viruses (dengue virus types 1 – 4 [DENV-1–4]) assigned to these species would be unaffected.

### Assignment of new species

Clearly, adoption of a divergence threshold for orthoflavivirus classification would require careful review of its impact on current taxonomy. In some instances, pragmatic decisions would have to be made for virus groups with divergence close to the threshold. For example, what would be most parsimonious way to assign SEPV and WESSV, and what would be the impact of splitting a well-known species such as *Orthoflavivirus denguei* into four species despite these sharing many features in common? However, adoption of the threshold would have enormous advantages for taxonomically assigning the many orthoflaviviruses that have remained unclassified for decades. This would include the primarily arthropod-specific fusiviruses for which information on host range, geography, and pathogenicity, as used in the current species definition, has no real relevance. The demarcation threshold would additionally enable classification of the large number of unclassified euflaviviruses, an important step for these well characterised viruses, many of which lie outside the established serocomplexes. Overall, application of genetic species demarcation threshold would enable around 24 new species to be added to the *Euflavivirus* subgenus and 33 new species to the *Fusivirus* subgenus, almost doubling the number of orthoflavivirus species.

### Taxonomic descriptions of distinct orthoflaviviruses within a species

One effect of applying a 0.1 *NS5* threshold for species demarcation is that several of the newly established species would harbour separately named viruses with distinctive properties that might otherwise be separately taxonomically assigned. One example described above would be the potential re-organization of *Orthoflavivirus zilberi*, *Orthoflavivirus mediterraneanense*, *Orthoflavivirus neudoerflense* and *Orthoflavivirus loupingi* into a single species even though the assigned viruses have only very recently been assigned to four separate monophyletic species based on their host and vectors, geography, and pathogenicity (25, 26).

It may be possible to maintain this differentiation while keeping a shared species assignment through the adoption of taxonomic classification below the species rank through their assignment to subspecies. This addition to virus taxonomy is currently under active discussion by the ICTV (Proposal 2025.001G; (36)) and with the wider virology community (37, 38). The use of subspecies assignments would also provide greater flexibility in virus taxonomic descriptions for other orthoflaviviruses such as LAMV and CHAOV, and the various lineages of WNV, including KOUV.

### Community feedback on potential future changes to *Orthoflavivirus* species taxonomy

As members of the current ICTV *Flaviviridae* Study Group, we believe that there is strong case for modification or replacement of the current rule for the assignment of species in the *Orthoflavivirus* genus through the use of genetic demarcation thresholds. However, feedback from the virological, medical and public health communities most affected by potential future changes to *Orthoflavivirus* taxonomy is vital in informing future formal taxonomic proposals to the ICTV. We have therefore a short survey form that we invite readers of the articles to complete: https://forms.gle/GD8MRGHKPsW4fyg59

Responses received by June 2027 will enable peer opinion to be taken into account before taxonomy proposal submission to the Executive Committee meeting of the ICTV in late summer, 2027.

## Conclusions

This study has analysed the potential impact of applying a uniform genetic assignment threshold for the taxonomic assignment of orthoflaviviruses. This step requires further discussion within the ICTV *Flaviviridae* Study Group and the wider virology community, but its application would be consistent with the increasing use of genetic relationships for virus classification, bring simplicity and consistency to the way species are established, enable orthoflaviviruses to be classified without the type of phenotypic information required by the current polythetic species definition, and finally give the large number of unassigned orthoflaviviruses a taxonomic home.

## Acknowledgements

Research reported in this publication was supported in part by the National Institute of Allergy and Infectious Diseases of the National Institutes of Health under Award Number U01AI151807 awarded to N.V.. The content is solely the responsibility of the authors and does not necessarily represent the official views of the National Institutes of Health. Support for the study was also funded by the U.S. Department of Veterans Affairs Merit Review Grant, Senior Clinical Investigator Award BX000207 and the VA SeqCURE Network funding (both to J.T.S.). A.K. is supported by grants from the National Institutes of Health, USA, award numbers AI185926 and AI183877. J.B. has been awarded the Distinguished Innovator grant from the Novo Nordisk Foundation (NNF23OC0087091) and the 2025 Fritz Kauffmann Award. D.B.S. is supported by a grant from the Microbiology Society, UK.

