## Supplementary Data for "Systematic genome analysis enables long-overdue taxonomic reorganization of genus *Orthoflavivirus*; potential assignment of over 50 new species"

SUPPLEMENTARY TABLES

**Table S1**. List of currently classified *Orthoflavivirus* species and example viruses (n = 57)

| **Subgenus** | **Species_name** | **Example virus** | **Vector** | **Avail**[1] |
|---|---|---|---|---|
| *Euflavivirus* | *Orthoflavivirus apoiense* | Apoi virus | Unknown | |
| *Euflavivirus* | *Orthoflavivirus aroaense* | Aroa virus | Mosquito | |
| *Euflavivirus* | *Orthoflavivirus bagazaense* | Bagaza virus | Mosquito | |
| *Euflavivirus* | *Orthoflavivirus banziense* | Banzi virus | Mosquito | |
| *Euflavivirus* | *Orthoflavivirus bouboulense* | Bouboui virus | Mosquito | |
| *Euflavivirus* | *Orthoflavivirus bravoense* | Rio_Bravo virus | Unknown | |
| *Euflavivirus* | *Orthoflavivirus bukalasaense* | Bukalasa_bat virus | Unknown | |
| *Euflavivirus* | *Orthoflavivirus cacipacoreense* | Cacipacore virus | Mosquito | N/A |
| *Euflavivirus* | *Orthoflavivirus careyense* | Carey Island virus | Unknown | N/A |
| *Euflavivirus* | *Orthoflavivirus cowboneense* | Cowbone Ridge virus | Unknown | |
| *Euflavivirus* | *Orthoflavivirus dakarense* | Dakar_bat virus | Unknown | |
| *Euflavivirus* | *Orthoflavivirus denguei* | Dengue virus type 2 | Mosquito | N/A |
| *Euflavivirus* | *Orthoflavivirus edgehillense* | Edge Hill virus | Mosquito | |
| *Euflavivirus* | *Orthoflavivirus entebbeense* | Entebbe_bat virus | Unknown | |
| *Euflavivirus* | *Orthoflavivirus flavi* | Yellow fever virus | Mosquito | |
| *Euflavivirus* | *Orthoflavivirus gadgetsense* | Gadgets_Gully virus | Tick | |
| *Euflavivirus* | *Orthoflavivirus ilheusense* | Ilheus virus | Mosquito | |
| *Euflavivirus* | *Orthoflavivirus israelense* | Israel turkey meningoencephamyelitis virus | Mosquito | |
| *Euflavivirus* | *Orthoflavivirus japonicum* | Japanese encephalititis virus | Mosquito | |
| *Euflavivirus* | *Orthoflavivirus jugraense* | Jugra virus | Mosquito | |
| *Euflavivirus* | *Orthoflavivirus jutiapaense* | Jutiapa virus | Unknown | |
| *Euflavivirus* | *Orthoflavivirus kadamense* | Kadam virus | Tick | |
| *Euflavivirus* | *Orthoflavivirus kedougouense* | Kedougou virus | Mosquito | |
| *Euflavivirus* | *Orthoflavivirus kokoberaorum* | Kokobera virus | Mosquito | |
| *Euflavivirus* | *Orthoflavivirus koutangoense* | Koutango virus | Mosquito | |
| *Euflavivirus* | *Orthoflavivirus kyasanurense* | Kyasanur Forest disease virus | Tick | |
| *Euflavivirus* | *Orthoflavivirus langatense* | Langat virus | Tick | |
| *Euflavivirus* | *Orthoflavivirus louisense* | St Louis encephalitis virus | Mosquito | N/A |
| *Euflavivirus* | *Orthoflavivirus loupingi* | Louping ill virus | Tick | |
| *Euflavivirus* | *Orthoflavivirus meabanense* | Meaban virus | Tick | |
| *Euflavivirus* | *Orthoflavivirus mediterranense* | Turkish sheep encephalitis virus | Tick | |
| *Euflavivirus* | *Orthoflavivirus modocense* | Modoc virus | Unknown | |
| *Euflavivirus* | *Orthoflavivirus montanaense* | Montana myotis leukoencephalitis virus | Unknown | |
| *Euflavivirus* | *Orthoflavivirus murrayense* | Murray valley encephalitis virus | Mosquito | |
| *Euflavivirus* | *Orthoflavivirus neudoerflense* | Tick-borne encephalititis virus | Tick | |
| *Euflavivirus* | *Orthoflavivirus nilense* | West Nile virus | Mosquito | |
| *Euflavivirus* | *Orthoflavivirus ntayaense* | Ntaya virus | Mosquito | |
| *Euflavivirus* | *Orthoflavivirus omskense* | Omsk haemmorhagic fever virus | Tick | |
| *Euflavivirus* | *Orthoflavivirus perlitaense* | San Perlita virus | Unknown | |
| *Euflavivirus* | *Orthoflavivirus phnompenhense* | Phnom_Penh_bat virus | Unknown | N/A |
| *Euflavivirus* | *Orthoflavivirus powassanense* | Powassan virus | Tick | |
| *Euflavivirus* | *Orthoflavivirus royalense* | Karshi virus | Tick | |
| *Euflavivirus* | *Orthoflavivirus saboyaense* | Saboya virus | Mosquito | |
| *Euflavivirus* | *Orthoflavivirus saumarezense* | Saumarez_Reef virus | Tick | |

| | | | |
|---|---|---|---|
| *Euflavivirus* | *Orthoflavivirus sepikense* | Sepik virus | Mosquito |
| *Euflavivirus* | *Orthoflavivirus tembusu* | Tembusu virus | Mosquito |
| *Euflavivirus* | *Orthoflavivirus tyuleniyense* | Tyuleniy virus | Tick |
| *Euflavivirus* | *Orthoflavivirus ugandaense* | Uganda_S virus | Mosquito |
| *Euflavivirus* | *Orthoflavivirus usutuense* | Usutu virus | Mosquito |
| *Euflavivirus* | *Orthoflavivirus viejaense* | Sal_Vieja virus | Unknown |
| *Euflavivirus* | *Orthoflavivirus wesselsbronense* | Wesselsbron virus | Mosquito |
| *Euflavivirus* | *Orthoflavivirus yaoundeense* | Yaounde virus | Mosquito |
| *Euflavivirus* | *Orthoflavivirus yokoseense* | Yokose virus | Unknown |
| *Euflavivirus* | *Orthoflavivirus zikaense* | Zika virus | Mosquito |
| *Euflavivirus* | *Orthoflavivirus zilberi* | Tick-borne encephalitis virus 1 | Tick |
| *Fusivirus* | *Orthoflavivirus iunctionis.* | Cell fusing agent | Insect |
| *Crangovirus* | *Orthovirus aphei* | Crangon crangon virus | Crustacean |

[1]N/A: not included; only incomplete genome sequence available

**Table S2.** List of sequences analysed in the study

| Genus | Virus_name | Species_name | Accession_number | Description |
|---|---|---|---|---|
| *Classified euflaviviruses* | | | | |
| *Euflavivirus* | Apoi virus | *Orthoflavivirus apoiense* | PZ305740 | Apoi virus strain Apoi SM7, |
| *Euflavivirus* | Aroa virus | *Orthoflavivirus aroaense* | KF917535 | Aroa virus strain Macaray 01809, |
| *Euflavivirus* | Aroa virus | *Orthoflavivirus aroaense* | PZ305741 | Aroa virus strain Macaray 01809, |
| *Euflavivirus* | Bussuquara virus | *Orthoflavivirus aroaense* | AY632536 | Bussuquara virus strain BeAn 4073, |
| *Euflavivirus* | Iguape virus | *Orthoflavivirus aroaense* | AY632538 | Iguape virus strain SPAn 71686, |
| *Euflavivirus* | Iguape virus | *Orthoflavivirus aroaense* | PZ324379 | Iguape virus strain SPAn 71686, |
| *Euflavivirus* | Naranjal virus | *Orthoflavivirus aroaense* | KF917538 | Naranjal virus strain 25008, |
| *Euflavivirus* | Bagaza virus | *Orthoflavivirus bagazaense* | AY632545 | Bagaza virus strain DakAr B209, |
| *Euflavivirus* | Bagaza virus | *Orthoflavivirus bagazaense* | EU684972 | Bagaza virus isolate 96363, |
| *Euflavivirus* | Bagaza virus | *Orthoflavivirus bagazaense* | HQ644144 | Bagaza virus isolate Spain B/2010, |
| *Euflavivirus* | Bagaza virus | *Orthoflavivirus bagazaense* | KR108244 | Bagaza virus isolate BAGV/Spain/RLP-Hcc1/2010, |
| *Euflavivirus* | Bagaza virus | *Orthoflavivirus bagazaense* | KR108246 | Bagaza virus isolate BAGV/Spain/RLP-Hcc3/2010, |
| *Euflavivirus* | Bagaza virus | *Orthoflavivirus bagazaense* | LC318701 | Bagaza virus,: |
| *Euflavivirus* | Bagaza virus | *Orthoflavivirus bagazaense* | MF380425 | Bagaza virus isolate ARD137998_DIAWARA_SEN_2000, |
| *Euflavivirus* | Bagaza virus | *Orthoflavivirus bagazaense* | MF380426 | Bagaza virus isolate ARD138018_DIAWARA_SEN_2000, |
| *Euflavivirus* | Bagaza virus | *Orthoflavivirus bagazaense* | MF380427 | Bagaza virus isolate ARD171075_BARKEDJI_SEN_2004, |
| *Euflavivirus* | Bagaza virus | *Orthoflavivirus bagazaense* | MF380428 | Bagaza virus isolate ARD171102_BARKEDJI_SEN_2004, |
| *Euflavivirus* | Bagaza virus | *Orthoflavivirus bagazaense* | MF380429 | Bagaza virus isolate ARA23139_Dezidougou_CI_1988, |
| *Euflavivirus* | Bagaza virus | *Orthoflavivirus bagazaense* | MF380430 | Bagaza virus isolate |
| *Euflavivirus* | Bagaza virus | *Orthoflavivirus bagazaense* | MF380431 | Bagaza virus isolate ARD99335_BARKEDJI_SEN_1994, |
| *Euflavivirus* | Bagaza virus | *Orthoflavivirus bagazaense* | MF380433 | Bagaza virus isolate ARD152146_DIAWARA_SEN_2001, |
| *Euflavivirus* | Bagaza virus | *Orthoflavivirus bagazaense* | MF380434 | Bagaza virus isolate ARD260266_BARKEDJI_SEN_2014, |
| *Euflavivirus* | Bagaza virus | *Orthoflavivirus bagazaense* | MW463911 | Bagaza virus isolate ZRU96/16/2, |
| *Euflavivirus* | Bagaza virus | *Orthoflavivirus bagazaense* | MW672101 | Bagaza virus strain MP314-NA-2018, |
| *Euflavivirus* | Bagaza virus | *Orthoflavivirus bagazaense* | PP236851 | Bagaza virus isolate BAGV_SPA/E/2021-01/RLP-b/3V, |
| *Euflavivirus* | Bagaza virus | *Orthoflavivirus bagazaense* | PP236852 | Bagaza virus isolate BAGV_SPA/E/2019-03/RLP-b/3V, |
| *Euflavivirus* | Banzi virus | *Orthoflavivirus banziense* | DQ859056 | Banzi virus strain SAH 336 polyprotein gene, |
| *Euflavivirus* | Banzi virus | *Orthoflavivirus banziense* | PZ305742 | Banzi virus strain SAH 336, |
| *Euflavivirus* | Bouboui virus | *Orthoflavivirus bouboulense* | DQ859057 | Bouboui virus strain DAK AR B490 polyprotein gene, |
| *Euflavivirus* | Rio Bravo virus | *Orthoflavivirus bravoense* | JQ582840 | Rio_Bravo virus strain M64, |
| *Euflavivirus* | Rio Bravo virus | *Orthoflavivirus bravoense* | PZ324384 | Rio_Bravo virus strain M64, |
| *Euflavivirus* | Bukalasa bat virus | *Orthoflavivirus bukalasaense* | OL709419 | Bukalasa_bat virus isolate BP111 sequence |
| *Euflavivirus* | Cacipacore virus | *Orthoflavivirus cacipacoreense* | KF917536 | Cacipacore virus strain BeAn 3276000, |
| *Euflavivirus* | Dakar bat virus | *Orthoflavivirus dakarense* | OL709418 | Dakar_bat virus isolate BP180, |
| *Euflavivirus* | Dakar bat virus | *Orthoflavivirus dakarense* | PZ305744 | Dakar_bat virus strain BP180, |
| *Euflavivirus* | Dengue virus type 1 | *Orthoflavivirus denguei* | EU482522 | DENV_1 isolate DENV-1/VN/BID-V978/2006, |
| *Euflavivirus* | Dengue virus type 1 | *Orthoflavivirus denguei* | FJ461323 | DENV_1 isolate DENV-1/VN/BID-V1887/2007, |
| *Euflavivirus* | Dengue virus type 1 | *Orthoflavivirus denguei* | FJ639820 | DENV_1 isolate DENV-1/VE/BID-V2260/2006, |
| *Euflavivirus* | Dengue virus type 1 | *Orthoflavivirus denguei* | FJ850103 | DENV_1 isolate DENV-1/VE/BID-V2468/2008, |
| *Euflavivirus* | Dengue virus type 1 | *Orthoflavivirus denguei* | GQ199797 | DENV_1 isolate DENV-1/VN/BID-V2774/2007, |
| *Euflavivirus* | Dengue virus type 1 | *Orthoflavivirus denguei* | GQ199835 | DENV_1 isolate DENV-1/VN/BID-V2840/2005, |
| *Euflavivirus* | Dengue virus type 1 | *Orthoflavivirus denguei* | GQ199845 | DENV_1 isolate DENV-1/VN/BID-V2881/2006, |
| *Euflavivirus* | Dengue virus type 1 | *Orthoflavivirus denguei* | GQ868509 | DENV_1 isolate DENV-1/MX/BID-V3698/2007, |
| *Euflavivirus* | Dengue virus type 1 | *Orthoflavivirus denguei* | GU131756 | DENV_1 isolate DENV-1/VN/BID-V3983/2008, |
| *Euflavivirus* | Dengue virus type 1 | *Orthoflavivirus denguei* | KJ189304 | DENV_1 isolate DENV-1/CO/BID-V7292/2005, |
| *Euflavivirus* | Dengue virus type 1 | *Orthoflavivirus denguei* | KJ189356 | DENV_1 isolate DENV-1/PR/BID-V7710/2012, |
| *Euflavivirus* | Dengue virus type 1 | *Orthoflavivirus denguei* | KM403613 | DENV_1 isolate SGEHI(D1)48695Y13, |
| *Euflavivirus* | Dengue virus type 1 | *Orthoflavivirus denguei* | KP686070 | DENV_1 isolate ww2014, |
| *Euflavivirus* | Dengue virus type 1 | *Orthoflavivirus denguei* | KX225491 | DENV_1 isolate GZ8_A3795/M/GZ/2006/DEV1 |
| *Euflavivirus* | Dengue virus type 1 | *Orthoflavivirus denguei* | KY057366 | DENV_1 isolate SUB-026A, |
| *Euflavivirus* | Dengue virus type 1 | *Orthoflavivirus denguei* | MN018289 | DENV_1 isolate D16009, |
| *Euflavivirus* | Dengue virus type 1 | *Orthoflavivirus denguei* | MN018297 | DENV_1 isolate D17026, |
| *Euflavivirus* | Dengue virus type 1 | *Orthoflavivirus denguei* | MN018329 | DENV_1 isolate D15033, |
| *Euflavivirus* | Dengue virus type 1 | *Orthoflavivirus denguei* | OZ494870 | dengue virus type 1 isolate 1-02-24-2924 genome |
| *Euflavivirus* | Dengue virus type 1 | *Orthoflavivirus denguei* | PX297359 | DENV_1 isolate DENV1/Bahamas/1977/Yale-DH006/P2, |
| *Euflavivirus* | Dengue virus type 2 | *Orthoflavivirus denguei* | AB479042 | DENV_2, |
| *Euflavivirus* | Dengue virus type 2 | *Orthoflavivirus denguei* | AY702034 | DENV_2 strain Cuba13/97, |
| *Euflavivirus* | Dengue virus type 2 | *Orthoflavivirus denguei* | DQ181805 | DENV_2 strain ThD2_0168_79, |
| *Euflavivirus* | Dengue virus type 2 | *Orthoflavivirus denguei* | EU482467 | DENV_2 isolate DENV-2/VN/BID-V921/2006, |
| *Euflavivirus* | Dengue virus type 2 | *Orthoflavivirus denguei* | EU482544 | DENV_2 isolate DENV-2/US/BID-V1031/2006, |
| *Euflavivirus* | Dengue virus type 2 | *Orthoflavivirus denguei* | EU482621 | DENV_2 isolate DENV-2/NI/BID-V1191/2007, |
| *Euflavivirus* | Dengue virus type 2 | *Orthoflavivirus denguei* | EU482736 | DENV_2 isolate DENV-2/US/BID-V679/1994, |
| *Euflavivirus* | Dengue virus type 2 | *Orthoflavivirus denguei* | EU482740 | DENV_2 isolate DENV-2/US/BID-V683/1994, |
| *Euflavivirus* | Dengue virus type 2 | *Orthoflavivirus denguei* | EU569709 | DENV_2 isolate DENV-2/US/BID-V1376/1996, |
| *Euflavivirus* | Dengue virus type 2 | *Orthoflavivirus denguei* | EU687215 | DENV_2 isolate DENV-2/US/BID-V1436/2004, |
| *Euflavivirus* | Dengue virus type 2 | *Orthoflavivirus denguei* | FJ744721 | DENV_2 isolate DENV-2/TH/BID-V2307/2001, |
| *Euflavivirus* | Dengue virus type 2 | *Orthoflavivirus denguei* | GQ398282 | DENV_2 strain DENV-2/PR/35DN/1994, |
| *Euflavivirus* | Dengue virus type 2 | *Orthoflavivirus denguei* | KC294212 | DENV_2 isolate DENV-2/PE/FPI01777/2011, |
| *Euflavivirus* | Dengue virus type 2 | *Orthoflavivirus denguei* | KM279550 | DENV_2 isolate DC663Y12, |
| *Euflavivirus* | Dengue virus type 2 | *Orthoflavivirus denguei* | KY474310 | DENV_2 isolate TD-00080-P, |
| *Euflavivirus* | Dengue virus type 2 | *Orthoflavivirus denguei* | MK783200 | DENV_2 isolate GZ8H/25, |
| *Euflavivirus* | Dengue virus type 2 | *Orthoflavivirus denguei* | MT180479 | DENV_2 isolate 231/17kandy, |
| *Euflavivirus* | Dengue virus type 2 | *Orthoflavivirus denguei* | MW577815 | DENV_2 isolate 168, |
| *Euflavivirus* | Dengue virus type 2 | *Orthoflavivirus denguei* | OP410990 | DENV_2 isolate SG(EHI)D2/41002Y19, |
| *Euflavivirus* | Dengue virus type 2 | *Orthoflavivirus denguei* | PV752123 | DENV_2 isolate SL_2017_TG41/1-10723, |
| *Euflavivirus* | Dengue virus type 3 | *Orthoflavivirus denguei* | EU081208 | DENV_3 strain D3/SG/05K3900DK1/2005, |
| *Euflavivirus* | Dengue virus type 3 | *Orthoflavivirus denguei* | FJ024466 | DENV_3 isolate DENV-3/US/BID-V1607/2004, |

| Genus | Virus_name | Species_name | Accession_number | Description |
|---|---|---|---|---|
| *Euflavivirus* | Dengue virus type 3 | *Orthoflavivirus denguei* | FJ182007 | DENV_3 isolate DENV-3/US/BID-V1617/2005, |
| *Euflavivirus* | Dengue virus type 3 | *Orthoflavivirus denguei* | FJ639784 | DENV_3 isolate DENV-3/VE/BID-V2217/2003, |
| *Euflavivirus* | Dengue virus type 3 | *Orthoflavivirus denguei* | FJ639805 | DENV_3 isolate DENV-3/VE/BID-V2242/2005, |
| *Euflavivirus* | Dengue virus type 3 | *Orthoflavivirus denguei* | FJ898472 | DENV_3 isolate DENV-3/VE/BID-V2969/2003, |
| *Euflavivirus* | Dengue virus type 3 | *Orthoflavivirus denguei* | GU131858 | DENV_3 isolate DENV-3/BR/BID-V3460/2006, |
| *Euflavivirus* | Dengue virus type 3 | *Orthoflavivirus denguei* | GU131943 | DENV_3 isolate DENV-3/IPC/BID-V4302/2007, |
| *Euflavivirus* | Dengue virus type 3 | *Orthoflavivirus denguei* | HM756278 | DENV_3 isolate DENV-3/NI/BID-V4813/2009, |
| *Euflavivirus* | Dengue virus type 3 | *Orthoflavivirus denguei* | JF808119 | DENV_3 isolate D3BR/BR8/04, |
| *Euflavivirus* | Dengue virus type 3 | *Orthoflavivirus denguei* | KJ189270 | DENV_3 isolate DENV-3/PE/BID-V7053/2007, |
| *Euflavivirus* | Dengue virus type 3 | *Orthoflavivirus denguei* | KJ189298 | DENV_3 isolate DENV-3/PE/BID-V7087/2008, |
| *Euflavivirus* | Dengue virus type 3 | *Orthoflavivirus denguei* | OP410997 | DENV_3 isolate SG(EHI)D3/04530Y19, |
| *Euflavivirus* | Dengue virus type 3 | *Orthoflavivirus denguei* | PP582637 | DENV_3 isolate 07178/DENV3/FG/2021-01-07 |
| *Euflavivirus* | Dengue virus type 3 | *Orthoflavivirus denguei* | PP582662 | DENV_3 isolate 20003/DENV3/FG/2023-07-20 |
| *Euflavivirus* | Dengue virus type 3 | *Orthoflavivirus denguei* | PV918774 | DENV_3 isolate SLS0049, |
| *Euflavivirus* | Dengue virus type 3 | *Orthoflavivirus denguei* | PX480334 | DENV_3 isolate D17-0442, |
| *Euflavivirus* | Dengue virus type 3 | *Orthoflavivirus denguei* | PX480418 | DENV_3 isolate D19-2035, |
| *Euflavivirus* | Dengue virus type 3 | *Orthoflavivirus denguei* | PX480431 | DENV_3 isolate D19-2240, |
| *Euflavivirus* | Dengue virus type 3 | *Orthoflavivirus denguei* | PX480435 | DENV_3 isolate D20-0183, |
| *Euflavivirus* | Dengue virus type 4 | *Orthoflavivirus denguei* | AY618990 | DENV_4 strain ThD4_0348_91, |
| *Euflavivirus* | Dengue virus type 4 | *Orthoflavivirus denguei* | FJ882583 | DENV_4 isolate DENV-4/VE/BID-V2492/2007, |
| *Euflavivirus* | Dengue virus type 4 | *Orthoflavivirus denguei* | GQ199881 | DENV_4 isolate DENV-4/US/BID-V2435/1996, |
| *Euflavivirus* | Dengue virus type 4 | *Orthoflavivirus denguei* | GQ868582 | DENV_4 isolate DENV-4/CO/BID-V3409/2001, |
| *Euflavivirus* | Dengue virus type 4 | *Orthoflavivirus denguei* | GQ868584 | DENV_4 isolate DENV-4/CO/BID-V3411/2004, |
| *Euflavivirus* | Dengue virus type 4 | *Orthoflavivirus denguei* | JF262783 | DENV_4 isolate INDIA G11337, |
| *Euflavivirus* | Dengue virus type 4 | *Orthoflavivirus denguei* | JN559741 | DENV_4 strain H772854, |
| *Euflavivirus* | Dengue virus type 4 | *Orthoflavivirus denguei* | JQ513330 | DENV_4 strain H772846, |
| *Euflavivirus* | Dengue virus type 4 | *Orthoflavivirus denguei* | JQ922558 | DENV_4 isolate DENV-4/IND/624000/1962, |
| *Euflavivirus* | Dengue virus type 4 | *Orthoflavivirus denguei* | KF041260 | DENV_4 isolate D4/Pakistan/150/2009, |
| *Euflavivirus* | Dengue virus type 4 | *Orthoflavivirus denguei* | KF955510 | DENV_4 isolate DENV-4/KH/BID-V2055/2002, |
| *Euflavivirus* | Dengue virus type 4 | *Orthoflavivirus denguei* | KJ160504 | DENV_4 isolate rDENV4, |
| *Euflavivirus* | Dengue virus type 4 | *Orthoflavivirus denguei* | KJ596670 | DENV_4 strain DENV-4/MT/BR76_TVP17953/2012 |
| *Euflavivirus* | Dengue virus type 4 | *Orthoflavivirus denguei* | KP188564 | DENV_4 isolate BR/SJRP/614/2013, |
| *Euflavivirus* | Dengue virus type 4 | *Orthoflavivirus denguei* | KP188566 | DENV_4 isolate BR/SJRP/850/2013, |
| *Euflavivirus* | Dengue virus type 4 | *Orthoflavivirus denguei* | KY672958 | DENV_4 isolate DENV-4/China/YN/15DGR35 (2015), |
| *Euflavivirus* | Dengue virus type 4 | *Orthoflavivirus denguei* | OK605767 | DENV_4 isolate DENV4-1_1229, |
| *Euflavivirus* | Dengue virus type 4 | *Orthoflavivirus denguei* | ON799403 | DENV_4 isolate THSTI-TRC-DENV4-02, |
| *Euflavivirus* | Dengue virus type 4 | *Orthoflavivirus denguei* | PP128486 | DENV_4 isolate 4II_AY762085_Singapur, |
| *Euflavivirus* | Dengue virus type 4 | *Orthoflavivirus denguei* | PX480445 | DENV_4 isolate D19-2229, |
| *Euflavivirus* | Entebbe bat virus | *Orthoflavivirus entebbeense* | DQ837641 | Entebbe_bat virus, |
| *Euflavivirus* | Entebbe bat virus | *Orthoflavivirus entebbeense* | KP233893 | Entebbe_bat virus strain UGA125, |
| *Euflavivirus* | Entebbe bat virus | *Orthoflavivirus entebbeense* | PQ720541 | Entebbe_bat virus clone UGA125, |
| *Euflavivirus* | Sokuluk virus | *Orthoflavivirus entebbeense* | KF917541 | Sokoluk virus strain LEIV-400K, |
| *Euflavivirus* | Sokuluk virus | *Orthoflavivirus entebbeense* | PZ324386 | Sokoluk virus strain LEIV-400K, |
| *Euflavivirus* | Yellow fever virus | *Orthoflavivirus flavi* | AF094612 | YFV strain Trinidad 79A isolate 788379 |
| *Euflavivirus* | Yellow fever virus | *Orthoflavivirus flavi* | JF912189 | YFV strain BeAR646536, |
| *Euflavivirus* | Yellow fever virus | *Orthoflavivirus flavi* | JX898869 | YFV isolate DakArAmt7 from Cote d'Ivoire |
| *Euflavivirus* | Yellow fever virus | *Orthoflavivirus flavi* | MF370531 | YFV strain ESS1115-H844801, |
| *Euflavivirus* | Yellow fever virus | *Orthoflavivirus flavi* | MK333804 | YFV strain GO27, |
| *Euflavivirus* | Yellow fever virus | *Orthoflavivirus flavi* | MK583166 | YFV isolate YFV043FIG, |
| *Euflavivirus* | Yellow fever virus | *Orthoflavivirus flavi* | MK728873 | YFV isolate GO21, |
| *Euflavivirus* | Yellow fever virus | *Orthoflavivirus flavi* | MN506267 | YFV strain MR 1044, |
| *Euflavivirus* | Yellow fever virus | *Orthoflavivirus flavi* | MN506288 | YFV strain VL 2964, |
| *Euflavivirus* | Yellow fever virus | *Orthoflavivirus flavi* | MN506289 | YFV strain VL 3012, |
| *Euflavivirus* | Yellow fever virus | *Orthoflavivirus flavi* | MN643092 | YFV strain RJ139, |
| *Euflavivirus* | Yellow fever virus | *Orthoflavivirus flavi* | MZ604852 | YFV strain SP13/BRA/2019 |
| *Euflavivirus* | Yellow fever virus | *Orthoflavivirus flavi* | MZ604855 | YFV strain SP15/BRA/2019 |
| *Euflavivirus* | Yellow fever virus | *Orthoflavivirus flavi* | MZ604865 | YFV strain SP12/BRA/2018 |
| *Euflavivirus* | Yellow fever virus | *Orthoflavivirus flavi* | OR509732 | YFV strain 17D-204, |
| *Euflavivirus* | Yellow fever virus | *Orthoflavivirus flavi* | PQ602526 | YFV isolate LEIAL3377/BRA/2024, |
| *Euflavivirus* | Yellow fever virus | *Orthoflavivirus flavi* | PQ879115 | YFV strain YFV/LEIAL4065/BRA/2024, |
| *Euflavivirus* | Yellow fever virus | *Orthoflavivirus flavi* | PQ879116 | YFV strain YFV/LEIAL4066/BRA/2024, |
| *Euflavivirus* | Yellow fever virus | *Orthoflavivirus flavi* | PV454340 | YFV strain YFV/LEIAL4700/BRA/2025, |
| *Euflavivirus* | Yellow fever virus | *Orthoflavivirus flavi* | PX379990 | YFV isolate PM435227, |
| *Euflavivirus* | Gadgets Gully virus | *Orthoflavivirus gadgetsense* | DQ235145 | Gadgets Gully virus from Australia |
| *Euflavivirus* | Ilheus virus | *Orthoflavivirus ilheusense* | AY632539 | Ilheus virus strain |
| *Euflavivirus* | Ilheus virus | *Orthoflavivirus ilheusense* | OP947885 | Ilheus virus strain BeH 7445, |
| *Euflavivirus* | Ilheus virus | *Orthoflavivirus ilheusense* | OP947886 | Ilheus virus strain Original, |
| *Euflavivirus* | Ilheus virus | *Orthoflavivirus ilheusense* | OP947889 | Ilheus virus strain ZCM 228, |
| *Euflavivirus* | Ilheus virus | *Orthoflavivirus ilheusense* | PQ553185 | Ilheus isolate An32356, |
| *Euflavivirus* | Ilheus virus | *Orthoflavivirus ilheusense* | PQ553186 | Ilheus isolate Ar9810, |
| *Euflavivirus* | Ilheus virus | *Orthoflavivirus ilheusense* | PQ553187 | Ilheus isolate Ar30600, |
| *Euflavivirus* | Ilheus virus | *Orthoflavivirus ilheusense* | PQ553188 | Ilheus isolate Ar30825, |
| *Euflavivirus* | Ilheus virus | *Orthoflavivirus ilheusense* | PQ553189 | Ilheus isolate Ar31540, |
| *Euflavivirus* | Ilheus virus | *Orthoflavivirus ilheusense* | PQ553190 | Ilheus isolate Ar33443, |
| *Euflavivirus* | Ilheus virus | *Orthoflavivirus ilheusense* | PQ553191 | Ilheus isolate Ar53247, |
| *Euflavivirus* | Ilheus virus | *Orthoflavivirus ilheusense* | PQ553192 | Ilheus isolate Ar54053, |
| *Euflavivirus* | Ilheus virus | *Orthoflavivirus ilheusense* | PQ553194 | Ilheus isolate Ar103409, |
| *Euflavivirus* | Ilheus virus | *Orthoflavivirus ilheusense* | PQ553197 | Ilheus isolate Ar423201, |
| *Euflavivirus* | Ilheus virus | *Orthoflavivirus ilheusense* | PQ553198 | Ilheus isolate Ar423237, |
| *Euflavivirus* | Ilheus virus | *Orthoflavivirus ilheusense* | PQ553199 | Ilheus isolate Ar425358, |
| *Euflavivirus* | Ilheus virus | *Orthoflavivirus ilheusense* | PQ553202 | Ilheus isolate H539894, |

| Genus | Virus_name | Species_name | Accession_number | Description |
|---|---|---|---|---|
| *Euflavivirus* | Ilheus virus | *Orthoflavivirus ilheusense* | PV844895 | Ilheus strain FPI22633, |
| *Euflavivirus* | Rocio virus | *Orthoflavivirus ilheusense* | AY632542 | Rocio virus strain SPH 34675, |
| *Euflavivirus* | Rocio virus | *Orthoflavivirus ilheusense* | MF461639 | Rocio virus strain SPH 34675, |
| *Euflavivirus* | Israel turkey | *Orthoflavivirus israelense* | KC734549 | ITMV, |
| *Euflavivirus* | Israel turkey | *Orthoflavivirus israelense* | KC734550 | ITMV isolate |
| *Euflavivirus* | Israel turkey | *Orthoflavivirus israelense* | KC734551 | ITMV isolate |
| *Euflavivirus* | Israel turkey | *Orthoflavivirus israelense* | KC734552 | ITMV isolate |
| *Euflavivirus* | Israel turkey | *Orthoflavivirus israelense* | KC734553 | ITMV isolate |
| *Euflavivirus* | Israel turkey | *Orthoflavivirus israelense* | KF917537 | ITMV strain ME 30502, |
| *Euflavivirus* | Japanese encephalititis virus | *Orthoflavivirus japonicum* | AF045551 | JEV strain K94P05, |
| *Euflavivirus* | Japanese encephalititis virus | *Orthoflavivirus japonicum* | AF254453 | JEV strain T1P1 |
| *Euflavivirus* | Japanese encephalititis virus | *Orthoflavivirus japonicum* | EF107523 | JEV strain WHe, |
| *Euflavivirus* | Japanese encephalititis virus | *Orthoflavivirus japonicum* | GQ199609 | JEV clone CNU/LP2x, |
| *Euflavivirus* | Japanese encephalititis virus | *Orthoflavivirus japonicum* | GU187972 | JEV strain SC04-17, |
| *Euflavivirus* | Japanese encephalititis virus | *Orthoflavivirus japonicum* | JF706283 | JEV isolate ZMT, |
| *Euflavivirus* | Japanese encephalititis virus | *Orthoflavivirus japonicum* | JN381840 | JEV isolate GSBY0810, |
| *Euflavivirus* | Japanese encephalititis virus | *Orthoflavivirus japonicum* | JN381841 | JEV isolate HN0421, |
| *Euflavivirus* | Japanese encephalititis virus | *Orthoflavivirus japonicum* | JN711459 | JEV strain HN2, |
| *Euflavivirus* | Japanese encephalititis virus | *Orthoflavivirus japonicum* | KT229574 | JEV strain YN09M57, |
| *Euflavivirus* | Japanese encephalititis virus | *Orthoflavivirus japonicum* | KT447437 | JEV strain Anyang-300, |
| *Euflavivirus* | Japanese encephalititis virus | *Orthoflavivirus japonicum* | KU323483 | JEV strain SA14, |
| *Euflavivirus* | Japanese encephalititis virus | *Orthoflavivirus japonicum* | KU871322 | JEV isolate C14B14, |
| *Euflavivirus* | Japanese encephalititis virus | *Orthoflavivirus japonicum* | KU871355 | JEV isolate BJ-1_BCP24, |
| *Euflavivirus* | Japanese encephalititis virus | *Orthoflavivirus japonicum* | KY609173 | JEV isolate BJ-1(P) polyprotein mRNA, |
| *Euflavivirus* | Japanese encephalititis virus | *Orthoflavivirus japonicum* | MF542268 | JEV isolate 10S3, |
| *Euflavivirus* | Japanese encephalititis virus | *Orthoflavivirus japonicum* | OR965960 | JEV isolate QLD_S46716_M2022 |
| *Euflavivirus* | Japanese encephalititis virus | *Orthoflavivirus japonicum* | PV472353 | JEV strain GX1209, |
| *Euflavivirus* | Japanese encephalititis virus | *Orthoflavivirus japonicum* | PV666542 | JEV strain EMAI17/NSW/22, |
| *Euflavivirus* | Japanese encephalititis virus | *Orthoflavivirus japonicum* | PV666560 | JEV strain EMAI40/NSW/22, |
| *Euflavivirus* | Japanese encephalititis virus | *Orthoflavivirus japonicum* | PV666568 | JEV strain EMAI51/NSW/22, |
| *Euflavivirus* | Japanese encephalititis virus | *Orthoflavivirus japonicum* | PV666599 | Japanese encephalitis virus strain P22-01661-20/QLD/22 |
| *Euflavivirus* | Japanese encephalititis virus | *Orthoflavivirus japonicum* | PX400461 | Japanese encephalitis virus strain SD-2025, |
| *Euflavivirus* | Jugra virus | *Orthoflavivirus jugraense* | DQ859066 | Jugra virus strain P-9-314 polyprotein gene, |
| *Euflavivirus* | Jutiapa virus | *Orthoflavivirus jutiapaense* | KJ469371 | Jutiapa virus strain JG-128, |
| *Euflavivirus* | Jutiapa virus | *Orthoflavivirus jutiapaense* | PZ324380 | Jutiapa virus strain JG-128, |
| *Euflavivirus* | Kadam virus | *Orthoflavivirus kadamense* | DQ235146 | Kadam virus from Uganda polyprotein gene, |
| *Euflavivirus* | Kedougou virus | *Orthoflavivirus kedougouense* | AY632540 | Kedougou virus strain DakAar D1470, |
| *Euflavivirus* | Kedougou virus | *Orthoflavivirus kedougouense* | PZ324381 | Kedougou virus strain DakAar D1470, |
| *Euflavivirus* | Bainyik virus | *Orthoflavivirus kokoberaorum* | KM225264 | Bainyik virus strain MK7979, |
| *Euflavivirus* | Kokobera virus | *Orthoflavivirus kokoberaorum* | AY632541 | Kokobera virus, |
| *Euflavivirus* | Kokobera virus | *Orthoflavivirus kokoberaorum* | OL347997 | Kokobera virus isolate A2019-0110, |
| *Euflavivirus* | New Mapoon virus | *Orthoflavivirus kokoberaorum* | KC788512 | New_Mapoon virus, |
| *Euflavivirus* | Stratford virus | *Orthoflavivirus kokoberaorum* | KF917540 | Stratford virus strain C338, |
| *Euflavivirus* | Stratford virus | *Orthoflavivirus kokoberaorum* | KM225263 | Stratford virus strain C338, |
| *Euflavivirus* | Stratford virus | *Orthoflavivirus kokoberaorum* | PP496992 | Stratford virus isolate 22 BrackenRidge, |
| *Euflavivirus* | Stratford virus | *Orthoflavivirus kokoberaorum* | PP496993 | Stratford virus isolate 76 Banyo, |
| *Euflavivirus* | Stratford virus | *Orthoflavivirus kokoberaorum* | PP496996 | Stratford virus isolate 295 Corinda, |
| *Euflavivirus* | Torres virus | *Orthoflavivirus kokoberaorum* | KM225265 | Torres virus strain TS5273, |
| *Euflavivirus* | Koutango virus | *Orthoflavivirus koutangoense* | MN057643 | Koutango virus isolate PM148, |
| *Euflavivirus* | Koutango virus | *Orthoflavivirus koutangoense* | OQ067500 | Koutango virus strain AnD5443, |
| *Euflavivirus* | Koutango virus | *Orthoflavivirus koutangoense* | PP489328 | Koutango virus isolate BAR_S3_S008, |
| *Euflavivirus* | Koutango virus | *Orthoflavivirus koutangoense* | PP885725 | Koutango virus isolate ISL_S6_2050, |
| *Euflavivirus* | Alkhumra virus | *Orthoflavivirus kyasanurense* | JF416954 | Alkhumra hemorrhagic fever virus strain 200300001, |
| *Euflavivirus* | Alkhumra virus | *Orthoflavivirus kyasanurense* | JF416956 | Alkhumra hemorrhagic fever virus strain Zaki #1, |
| *Euflavivirus* | Alkhumra virus | *Orthoflavivirus kyasanurense* | JF416961 | Alkhumra hemorrhagic fever virus strain 200904529, |
| *Euflavivirus* | Alkhumra virus | *Orthoflavivirus kyasanurense* | JF416962 | Alkhumra hemorrhagic fever virus strain 200905926, |
| *Euflavivirus* | Alkhumra virus | *Orthoflavivirus kyasanurense* | JF416966 | Alkhumra hemorrhagic fever virus strain 200905921, |
| *Euflavivirus* | Alkhumra virus | *Orthoflavivirus kyasanurense* | JX271892 | Alkhumra hemorrhagic fever virus strain 1336, |
| *Euflavivirus* | Kyasanur Forest disease virus | *Orthoflavivirus kyasanurense* | HM055369 | KFDV strain KFD P 9605, |
| *Euflavivirus* | Kyasanur Forest disease virus | *Orthoflavivirus kyasanurense* | JF416960 | KFDV strain W-377, |
| *Euflavivirus* | Kyasanur Forest disease virus | *Orthoflavivirus kyasanurense* | MH013227 | KFDV strain NIV16877, |
| *Euflavivirus* | Kyasanur Forest disease virus | *Orthoflavivirus kyasanurense* | OP037814 | KFDV isolate KFDV, |
| *Euflavivirus* | Kyasanur Forest disease virus | *Orthoflavivirus kyasanurense* | OQ866118 | KFDV isolate |
| *Euflavivirus* | Kyasanur Forest disease virus | *Orthoflavivirus kyasanurense* | OR162010 | KFDV isolate MCL-19-H-561-MH_P1_11477, |
| *Euflavivirus* | Kyasanur Forest disease virus | *Orthoflavivirus kyasanurense* | OR162013 | KFDV isolate MCL-19-H-650-MH_P1_11483, |
| *Euflavivirus* | Kyasanur Forest disease virus | *Orthoflavivirus kyasanurense* | OR162014 | KFDV isolate MCL-19-H-686-MH_P1_11486, |
| *Euflavivirus* | Kyasanur Forest disease virus | *Orthoflavivirus kyasanurense* | OR162016 | KFDV isolate MCL-19-H-689-MH_P1_11492, |
| *Euflavivirus* | Kyasanur Forest disease virus | *Orthoflavivirus kyasanurense* | OR162018 | KFDV isolate MCL-19-H-897-MH_P1_11498, |
| *Euflavivirus* | Kyasanur Forest disease virus | *Orthoflavivirus kyasanurense* | OR179910 | KFDV isolate NIV142856_KA_P1_10842, |
| *Euflavivirus* | Kyasanur Forest disease virus | *Orthoflavivirus kyasanurense* | PP597397 | KFDV isolate NIV_164187_plaque_1.1, |
| *Euflavivirus* | Kyasanur Forest disease virus | *Orthoflavivirus kyasanurense* | PP597398 | KFDV isolate NIV_164187_plaque_3.2, |
| *Euflavivirus* | Kyasanur Forest disease virus | *Orthoflavivirus kyasanurense* | PP763477 | KFDV isolate NIV_164187_Plaque-1.1, |
| *Euflavivirus* | Langat virus | *Orthoflavivirus langatense* | AF253419 | Langat virus, |
| *Euflavivirus* | Langat virus | *Orthoflavivirus langatense* | EU790644 | Langat virus strain TP21, |
| *Euflavivirus* | Langat virus | *Orthoflavivirus langatense* | MK680893 | Langat virus strain T 1674, |
| *Euflavivirus* | Langat virus | *Orthoflavivirus langatense* | OP037820 | Langat virus isolate LGT 1, |
| *Euflavivirus* | Langat virus | *Orthoflavivirus langatense* | OP037821 | Langat virus isolate LGT 2, |
| *Euflavivirus* | Langat virus | *Orthoflavivirus langatense* | OP037822 | Langat virus isolate LGT 3, |
| *Euflavivirus* | St Louis encephalitis virus | *Orthoflavivirus louisense* | EU566860 | SLEV strain Hubbard, |
| *Euflavivirus* | St Louis encephalitis virus | *Orthoflavivirus louisense* | FJ753286 | SLEV strain CbaAr-4005, |
| *Euflavivirus* | St Louis encephalitis virus | *Orthoflavivirus louisense* | FJ753287 | SLEV strain 79V-2533, |

| Genus | Virus_name | Species_name | Accession_number | Description |
|---|---|---|---|---|
| *Euflavivirus* | St Louis encephalitis virus | *Orthoflavivirus louisense* | KX258462 | SLEV isolate 39, |
| *Euflavivirus* | St Louis encephalitis virus | *Orthoflavivirus louisense* | PX678853 | SLEV isolate MERC702, |
| *Euflavivirus* | St Louis encephalitis virus | *Orthoflavivirus louisense* | PZ205910 | SLEV isolate 5016, |
| *Euflavivirus* | St Louis encephalitis virus | *Orthoflavivirus louisense* | PZ205912 | SLEV isolate 5193, |
| *Euflavivirus* | St Louis encephalitis virus | *Orthoflavivirus louisense* | PZ205913 | SLEV isolate 5566, |
| *Euflavivirus* | St Louis encephalitis virus | *Orthoflavivirus louisense* | PZ205914 | SLEV isolate 5569, |
| *Euflavivirus* | St Louis encephalitis virus | *Orthoflavivirus louisense* | PZ205915 | SLEV isolate 6033, |
| *Euflavivirus* | St Louis encephalitis virus | *Orthoflavivirus louisense* | PZ205920 | SLEV isolate 6741, |
| *Euflavivirus* | St Louis encephalitis virus | *Orthoflavivirus louisense* | PZ205922 | SLEV isolate 7000, |
| *Euflavivirus* | St Louis encephalitis virus | *Orthoflavivirus louisense* | PZ205932 | SLEV isolate 4992, |
| *Euflavivirus* | St Louis encephalitis virus | *Orthoflavivirus louisense* | PZ205933 | SLEV isolate 5001, |
| *Euflavivirus* | St Louis encephalitis virus | *Orthoflavivirus louisense* | PZ205935 | SLEV isolate 5292, |
| *Euflavivirus* | St Louis encephalitis virus | *Orthoflavivirus louisense* | PZ205940 | SLEV isolate 5987, |
| *Euflavivirus* | St Louis encephalitis virus | *Orthoflavivirus louisense* | PZ205942 | SLEV isolate 6027, |
| *Euflavivirus* | St Louis encephalitis virus | *Orthoflavivirus louisense* | PZ205943 | SLEV isolate 6034, |
| *Euflavivirus* | St Louis encephalitis virus | *Orthoflavivirus louisense* | PZ205949 | SLEV isolate 7204, |
| *Euflavivirus* | St Louis encephalitis virus | *Orthoflavivirus louisense* | PZ205967 | SLEV isolate 140473, |
| *Euflavivirus* | Louping ill virus | *Orthoflavivirus loupingi* | DQ235152 | Spanish sheep encephalitis virus |
| *Euflavivirus* | Louping ill virus | *Orthoflavivirus loupingi* | KF056331 | Louping ill virus isolate Penrith |
| *Euflavivirus* | Louping ill virus | *Orthoflavivirus loupingi* | KJ495985 | Louping ill virus isolate Primorye-185-91 |
| *Euflavivirus* | Louping ill virus | *Orthoflavivirus loupingi* | KP144331 | Louping ill virus isolate LI3/1 |
| *Euflavivirus* | Louping ill virus | *Orthoflavivirus loupingi* | KT224354 | Louping ill virus strain LEIV-7435Tur |
| *Euflavivirus* | Louping ill virus | *Orthoflavivirus loupingi* | KT224355 | Negishi virus |
| *Euflavivirus* | Louping ill virus | *Orthoflavivirus loupingi* | MG243699 | Tick-borne encephalitis virus strain N5- |
| *Euflavivirus* | Louping ill virus | *Orthoflavivirus loupingi* | MK007533 | Louping ill virus isolate ENG_PEN6_2009 |
| *Euflavivirus* | Louping ill virus | *Orthoflavivirus loupingi* | MK007538 | Louping ill virus isolate SCO_K_1980 |
| *Euflavivirus* | Louping ill virus | *Orthoflavivirus loupingi* | MK007543 | Louping ill virus isolate SCO_INV1_1983 |
| *Euflavivirus* | Louping ill virus | *Orthoflavivirus loupingi* | MK007546 | Louping ill virus isolate ENG_DEV4_1995 |
| *Euflavivirus* | Louping ill virus | *Orthoflavivirus loupingi* | MK007547 | Louping ill virus isolate ENG_DEV2_1989 |
| *Euflavivirus* | Louping ill virus | *Orthoflavivirus loupingi* | MK007548 | Louping ill virus isolate ENG_DEV1_1983 |
| *Euflavivirus* | Louping ill virus | *Orthoflavivirus loupingi* | MN844186 | Louping ill virus isolate PRES1 |
| *Euflavivirus* | Louping ill virus | *Orthoflavivirus loupingi* | ON502378 | Tick-borne encephalitis virus strain Salland (NL), |
| *Euflavivirus* | Louping ill virus | *Orthoflavivirus loupingi* | PP782054 | Alpine chamois encephalitis virus strain cds |
| *Euflavivirus* | Louping ill virus | *Orthoflavivirus loupingi* | PP795456 | Alpine chamois encephalitis virus isolate Tick\63100- |
| *Euflavivirus* | Louping ill virus | *Orthoflavivirus loupingi* | PP795457 | Alpine chamois encephalitis virus isolate |
| *Euflavivirus* | Louping ill virus | *Orthoflavivirus loupingi* | PP882705 | Louping ill virus strain L_178 |
| *Euflavivirus* | Louping ill virus | *Orthoflavivirus loupingi* | Y07863 | Louping ill virus |
| *Euflavivirus* | Meaban virus | *Orthoflavivirus meabanense* | DQ235144 | Meaban virus from France polyprotein gene, |
| *Euflavivirus* | Tick-borne encephalitis virus | *O. mediterraneanense* | DQ235151 | Turkish sheep encephalitis virus cds |
| *Euflavivirus* | Tick-borne encephalitis virus | *O. mediterraneanense* | DQ235153 | Greek goat encephalitis virus cds |
| *Euflavivirus* | Modoc virus | *Orthoflavivirus modocense* | AJ242984 | Modoc virus, |
| *Euflavivirus* | Modoc virus | *Orthoflavivirus modocense* | OQ095369 | Modoc virus isolate 83, |
| *Euflavivirus* | Modoc virus | *Orthoflavivirus modocense* | PZ324382 | Modoc virus strain M544, |
| *Euflavivirus* | Montana myotis leukoencephalitis | *Orthoflavivirus montanaense* | AJ299445 | Montana myotis leukoencephalitis virus, |
| *Euflavivirus* | Alfuy virus | *Orthoflavivirus murrayense* | AY898809 | Alfuy virus strain MRM3929, |
| *Euflavivirus* | Murray valley encephalitis virus | *Orthoflavivirus murrayense* | AF161266 | MVEV strain MVE-1-51, |
| *Euflavivirus* | Murray valley encephalitis virus | *Orthoflavivirus murrayense* | KC852189 | MVEV isolate MVE-1-51-P0 |
| *Euflavivirus* | Murray valley encephalitis virus | *Orthoflavivirus murrayense* | KC852192 | MVEV isolate MVE-1-51-CD8 |
| *Euflavivirus* | Murray valley encephalitis virus | *Orthoflavivirus murrayense* | KC852193 | MVEV isolate OR156-P0, |
| *Euflavivirus* | Murray valley encephalitis virus | *Orthoflavivirus murrayense* | KC852196 | MVEV isolate OR156-CD8 |
| *Euflavivirus* | Murray valley encephalitis virus | *Orthoflavivirus murrayense* | OQ418654 | MVEV isolate CY1189, |
| *Euflavivirus* | Murray valley encephalitis virus | *Orthoflavivirus murrayense* | OQ418656 | MVEV isolate CY2692, |
| *Euflavivirus* | Murray valley encephalitis virus | *Orthoflavivirus murrayense* | OQ418657 | MVEV isolate Gu0091, |
| *Euflavivirus* | Murray valley encephalitis virus | *Orthoflavivirus murrayense* | OQ418662 | MVEV isolate K10456, |
| *Euflavivirus* | Murray valley encephalitis virus | *Orthoflavivirus murrayense* | OQ418671 | MVEV isolate K31302, |
| *Euflavivirus* | Murray valley encephalitis virus | *Orthoflavivirus murrayense* | OQ418672 | MVEV isolate K42307, |
| *Euflavivirus* | Murray valley encephalitis virus | *Orthoflavivirus murrayense* | OQ418681 | MVEV isolate K73146, |
| *Euflavivirus* | Murray valley encephalitis virus | *Orthoflavivirus murrayense* | OQ418685 | MVEV isolate K73475, |
| *Euflavivirus* | Murray valley encephalitis virus | *Orthoflavivirus murrayense* | OQ418689 | MVEV isolate K74558, |
| *Euflavivirus* | Murray valley encephalitis virus | *Orthoflavivirus murrayense* | OQ418692 | MVEV isolate K75678, |
| *Euflavivirus* | Murray valley encephalitis virus | *Orthoflavivirus murrayense* | OQ418698 | MVEV isolate K84754, |
| *Euflavivirus* | Murray valley encephalitis virus | *Orthoflavivirus murrayense* | OQ418699 | MVEV isolate K85133, |
| *Euflavivirus* | Murray valley encephalitis virus | *Orthoflavivirus murrayense* | OQ418715 | MVEV isolate P6513, |
| *Euflavivirus* | Murray valley encephalitis virus | *Orthoflavivirus murrayense* | OQ418722 | MVEV isolate PNG6910, |
| *Euflavivirus* | Tick-borne encephalitis virus | *Orthoflavivirus neudoerflense* | GU183381 | Tick-borne encephalitis virus strain Joutseno, |
| *Euflavivirus* | Tick-borne encephalititis virus | *Orthoflavivirus neudoerflense* | PQ442018 | Tick-borne encephalitis virus isolate KME-EU-486 (POLY) |
| *Euflavivirus* | Tick-borne encephalititis virus | *Orthoflavivirus neudoerflense* | LC171402 | Tick-borne encephalitis virus genomic RNA, strain: |
| *Euflavivirus* | Tick-borne encephalititis virus | *Orthoflavivirus neudoerflense* | KJ922513 | Tick-borne encephalitis virus strain Petracova |
| *Euflavivirus* | Tick-borne encephalititis virus | *Orthoflavivirus neudoerflense* | KU884607 | Tick-borne encephalitis virus isolate Leila-BH95/15, |
| *Euflavivirus* | Tick-borne encephalititis virus | *Orthoflavivirus neudoerflense* | MT228628 | Tick-borne encephalitis virus clone Hypr_variant_40_PS |
| *Euflavivirus* | Tick-borne encephalititis virus | *Orthoflavivirus neudoerflense* | PQ442011 | Tick-borne encephalitis virus isolate KME-R2817 (POLY) |
| *Euflavivirus* | Tick-borne encephalititis virus | *Orthoflavivirus neudoerflense* | MK560446 | Tick-borne encephalitis virus strain 172-68 |
| *Euflavivirus* | Tick-borne encephalititis virus | *Orthoflavivirus neudoerflense* | PQ442025 | Tick-borne encephalitis virus isolate 2024-NGS-1663 |
| *Euflavivirus* | Tick-borne encephalititis virus | *Orthoflavivirus neudoerflense* | PZ383778 | MAG: Tick-borne encephalitis virus isolate SOR-F3 |
| *Euflavivirus* | Tick-borne encephalititis virus | *Orthoflavivirus neudoerflense* | ON228432 | Tick-borne encephalitis virus isolate 11599, |
| *Euflavivirus* | Tick-borne encephalititis virus | *Orthoflavivirus neudoerflense* | OQ889245 | Tick-borne encephalitis virus isolate 2020-P71-S6-like |
| *Euflavivirus* | Tick-borne encephalititis virus | *Orthoflavivirus neudoerflense* | MK801804 | Tick-borne encephalitis virus isolate Sipoo-4-Finland- |
| *Euflavivirus* | Tick-borne encephalititis virus | *Orthoflavivirus neudoerflense* | GQ266392 | Tick-borne encephalitis virus isolate AS33 |
| *Euflavivirus* | Tick-borne encephalititis virus | *Orthoflavivirus neudoerflense* | PZ361878 | Tick-borne encephalitis virus isolate |
| *Euflavivirus* | Tick-borne encephalititis virus | *Orthoflavivirus neudoerflense* | HM535610 | Tick-borne encephalitis virus strain KrM 213 |
| *Euflavivirus* | Tick-borne encephalititis virus | *Orthoflavivirus neudoerflense* | MT228627 | Tick-borne encephalitis virus clone |

| Genus | Virus_name | Species_name | Accession_number | Description |
|---|---|---|---|---|
| *Euflavivirus* | Tick-borne encephalititis virus | *Orthoflavivirus neudoerflense* | MN661145 | Tick-borne encephalitis virus isolate UK-Hampshire 2019 |
| *Euflavivirus* | Tick-borne encephalititis virus | *Orthoflavivirus neudoerflense* | OQ889184 | Tick-borne encephalitis virus isolate 2009-P88-S1-like |
| *Euflavivirus* | Tick-borne encephalititis virus | *Orthoflavivirus neudoerflense* | MZ969639 | Tick-borne encephalitis virus isolate NL-RMB2, |
| *Euflavivirus* | West Nile virus | *Orthoflavivirus nilense* | AF260969 | WNV strain RO97-50, |
| *Euflavivirus* | West Nile virus | *Orthoflavivirus nilense* | DQ666449 | WNV isolate GCTX1-2005, |
| *Euflavivirus* | West Nile virus | *Orthoflavivirus nilense* | HM488131 | WNV isolate WNV-1/US/BID-V4193/2000, |
| *Euflavivirus* | West Nile virus | *Orthoflavivirus nilense* | HM756658 | WNV isolate WNV-1/US/BID-V4686/2003, |
| *Euflavivirus* | West Nile virus | *Orthoflavivirus nilense* | KJ145828 | WNV isolate WNV-1/US/BID-V6057/2010, |
| *Euflavivirus* | West Nile virus | *Orthoflavivirus nilense* | KJ501336 | WNV isolate WNV-1/US/BID-V6517/2001, |
| *Euflavivirus* | West Nile virus | *Orthoflavivirus nilense* | KJ501445 | WNV isolate WNV-1/US/BID-V6381/2001, |
| *Euflavivirus* | West Nile virus | *Orthoflavivirus nilense* | KJ501475 | WNV isolate WNV-1/US/BID-V6471/2002, |
| *Euflavivirus* | West Nile virus | *Orthoflavivirus nilense* | KJ501533 | WNV isolate WNV-1/US/BID-V7817/2012, |
| *Euflavivirus* | West Nile virus | *Orthoflavivirus nilense* | KX547357 | WNV strain WNV-1/Culex/USA/07510530/2007 |
| *Euflavivirus* | West Nile virus | *Orthoflavivirus nilense* | KX547381 | WNV strain WNV-1/Cyanocitta cristata |
| *Euflavivirus* | West Nile virus | *Orthoflavivirus nilense* | KX547392 | WNV strain WNV-1/Culex/USA/07141331/2007 |
| *Euflavivirus* | West Nile virus | *Orthoflavivirus nilense* | KX547522 | WNV strain WNV-1/Corvus |
| *Euflavivirus* | West Nile virus | *Orthoflavivirus nilense* | MN794935 | WNV strain BNI-10/19, |
| *Euflavivirus* | West Nile virus | *Orthoflavivirus nilense* | OL840893 | WNV isolate GR-Serres/S1m/2020, |
| *Euflavivirus* | West Nile virus | *Orthoflavivirus nilense* | OQ053524 | WNV strain GR-Imathia/472m/2022, |
| *Euflavivirus* | West Nile virus | *Orthoflavivirus nilense* | OZ309026 | WNV isolate VV01-25-Q-16825-WNV-M08760-240723 |
| *Euflavivirus* | West Nile virus | *Orthoflavivirus nilense* | OZ310047 | WNV isolate UNMC0685 genome assembly,: |
| *Euflavivirus* | West Nile virus | *Orthoflavivirus nilense* | OZ310094 | WNV isolate UNMC0688 genome assembly,: |
| *Euflavivirus* | West Nile virus | *Orthoflavivirus nilense* | OZ310181 | WNV isolate UNMC0711 genome assembly,: |
| *Euflavivirus* | Ntaya virus | *Orthoflavivirus ntayaense* | JX236040 | Ntaya virus, |
| *Euflavivirus* | Ntaya virus | *Orthoflavivirus ntayaense* | KF917539 | Ntaya virus strain Original, |
| *Euflavivirus* | Omsk haemmorhagic fever virus | *Orthoflavivirus omskense* | AB507800 | OHFV,, strain: |
| *Euflavivirus* | Omsk haemmorhagic fever virus | *Orthoflavivirus omskense* | AY438626 | OHFV strain Kubrin, |
| *Euflavivirus* | Omsk haemmorhagic fever virus | *Orthoflavivirus omskense* | MT354616 | OHFV strain OZ-97/11285, |
| *Euflavivirus* | Omsk haemmorhagic fever virus | *Orthoflavivirus omskense* | MT354619 | OHFV strain Kabyrdak-39/10944 |
| *Euflavivirus* | Omsk haemmorhagic fever virus | *Orthoflavivirus omskense* | MT354621 | OHFV strain Veselovka-753/11084 |
| *Euflavivirus* | Omsk haemmorhagic fever virus | *Orthoflavivirus omskense* | MT354623 | OHFV strain B-37/9866, |
| *Euflavivirus* | Omsk haemmorhagic fever virus | *Orthoflavivirus omskense* | MT354625 | OHFV strain B-1/10186, |
| *Euflavivirus* | Omsk haemmorhagic fever virus | *Orthoflavivirus omskense* | MT354628 | OHFV strain 42M/9722, |
| *Euflavivirus* | Omsk haemmorhagic fever virus | *Orthoflavivirus omskense* | OL689364 | OHFV strain Liv-356-0/13967 |
| *Euflavivirus* | Omsk haemmorhagic fever virus | *Orthoflavivirus omskense* | OL689366 | OHFV strain Br-356-1/13959 |
| *Euflavivirus* | Omsk haemmorhagic fever virus | *Orthoflavivirus omskense* | OL689369 | OHFV strain Kid-362/13944, |
| *Euflavivirus* | Omsk haemmorhagic fever virus | *Orthoflavivirus omskense* | OL689370 | OHFV strain Liv-362/13943, |
| *Euflavivirus* | Omsk haemmorhagic fever virus | *Orthoflavivirus omskense* | OL689376 | OHFV strain Br-356-1/13969 |
| *Euflavivirus* | Omsk haemmorhagic fever virus | *Orthoflavivirus omskense* | OL689378 | OHFV strain Ur-353/13965, |
| *Euflavivirus* | Omsk haemmorhagic fever virus | *Orthoflavivirus omskense* | OL689383 | OHFV strain Br-352/352, |
| *Euflavivirus* | Omsk haemmorhagic fever virus | *Orthoflavivirus omskense* | OL689384 | OHFV strain Br-361/361, |
| *Euflavivirus* | Omsk haemmorhagic fever virus | *Orthoflavivirus omskense* | OL689385 | OHFV strain Br-361/13948, |
| *Euflavivirus* | Omsk haemmorhagic fever virus | *Orthoflavivirus omskense* | OL689386 | OHFV strain Ur-359/13965, |
| *Euflavivirus* | Omsk haemmorhagic fever virus | *Orthoflavivirus omskense* | OL689387 | OHFV strain Br-354/354, |
| *Euflavivirus* | Omsk haemmorhagic fever virus | *Orthoflavivirus omskense* | OP292291 | OHFV strain MO-1007, |
| *Euflavivirus* | Batu Cave virus | *Orthoflavivirus phnompenhense* | KJ469370 | Batu_Cave virus strain P70-1459, |
| *Euflavivirus* | Phnom Penh bat virus | *Orthoflavivirus phnompenhense* | KJ469372 | Phnom_Penh_bat virus strain 30834_A38, |
| *Euflavivirus* | Phnom Penh bat virus | *Orthoflavivirus phnompenhense* | PZ324383 | Phnom_Penh_bat virus strain 30834_A38, |
| *Euflavivirus* | Deer tick virus | *Orthoflavivirus powassanense* | OP037825 | Deer tick virus isolate DTV IPS001 2, |
| *Euflavivirus* | Deer tick virus | *Orthoflavivirus powassanense* | PX666892 | Deer tick virus isolate R59266, |
| *Euflavivirus* | Powassan virus | *Orthoflavivirus powassanense* | EU543649 | Powassan virus strain Partizansk/2006, |
| *Euflavivirus* | Powassan virus | *Orthoflavivirus powassanense* | OL695841 | Powassan virus isolate PV320, |
| *Euflavivirus* | Powassan virus | *Orthoflavivirus powassanense* | OL704153 | Powassan virus isolate P0525, |
| *Euflavivirus* | Powassan virus | *Orthoflavivirus powassanense* | OL704166 | Powassan virus isolate P1113, |
| *Euflavivirus* | Powassan virus | *Orthoflavivirus powassanense* | OL704176 | Powassan virus isolate P1753, |
| *Euflavivirus* | Powassan virus | *Orthoflavivirus powassanense* | OL704184 | Powassan virus isolate 188, |
| *Euflavivirus* | Powassan virus | *Orthoflavivirus powassanense* | OL704188 | Powassan virus isolate 3338, |
| *Euflavivirus* | Powassan virus | *Orthoflavivirus powassanense* | OL704223 | Powassan virus isolate 1341, |
| *Euflavivirus* | Powassan virus | *Orthoflavivirus powassanense* | OL704273 | Powassan virus isolate 17079-117, |
| *Euflavivirus* | Powassan virus | *Orthoflavivirus powassanense* | OL704279 | Powassan virus isolate 14CD13-002, |
| *Euflavivirus* | Powassan virus | *Orthoflavivirus powassanense* | OL704289 | Powassan virus isolate 15CD59-002, |
| *Euflavivirus* | Powassan virus | *Orthoflavivirus powassanense* | OL704290 | Powassan virus isolate 12021-240, |
| *Euflavivirus* | Powassan virus | *Orthoflavivirus powassanense* | OL704295 | Powassan virus isolate 19001-415, |
| *Euflavivirus* | Powassan virus | *Orthoflavivirus powassanense* | OL704320 | Powassan virus isolate 15-S13-03, |
| *Euflavivirus* | Powassan virus | *Orthoflavivirus powassanense* | OL704326 | Powassan virus isolate 19103-1357, |
| *Euflavivirus* | Powassan virus | *Orthoflavivirus powassanense* | OL704339 | Powassan virus isolate 13CD01-002, |
| *Euflavivirus* | Powassan virus | *Orthoflavivirus powassanense* | OL704381 | Powassan virus isolate 19039-373, |
| *Euflavivirus* | Powassan virus | *Orthoflavivirus powassanense* | OP265691 | Powassan virus isolate 63-002, |
| *Euflavivirus* | Karshi virus | *Orthoflavivirus royalense* | AY863002 | Karshi virus strain LEIV 2247, |
| *Euflavivirus* | Karshi virus | *Orthoflavivirus royalense* | DQ462443 | Karshi virus strain LEIV-7192 Tur, |
| *Euflavivirus* | Royal Farm virus | *Orthoflavivirus royalense* | DQ235149 | Royal_Farm virus from Afghanistan polyprotein gene, |
| *Euflavivirus* | Potiskum virus | *Orthoflavivirus saboyaense* | DQ859067 | Potiskum virus strain IBAN 10069 polyprotein gene, |
| *Euflavivirus* | Saboya virus | *Orthoflavivirus saboyaense* | DQ859062 | Saboya virus strain Dak AR D4600 polyprotein gene, |
| *Euflavivirus* | Saumarez Reef virus | *Orthoflavivirus saumarezense* | DQ235150 | Saumarez_Reef virus from Australia polyprotein gene, |
| *Euflavivirus* | Saumarez Reef virus | *Orthoflavivirus saumarezense* | MW959115 | Saumarez_Reef virus isolate W11_2232 pool, |
| *Euflavivirus* | Sepik virus | *Orthoflavivirus sepikense* | DQ837642 | Sepik virus strain MK7148, |
| *Euflavivirus* | Tembusu virus | *Orthoflavivirus tembusu* | JX273153 | Tembusu virus strain JS/2010, |
| *Euflavivirus* | Tembusu virus | *Orthoflavivirus tembusu* | KC990540 | Tembusu virus strain BZ_2010, |
| *Euflavivirus* | Tembusu virus | *Orthoflavivirus tembusu* | KF192951 | Tembusu virus isolate Tembusu virus strain, |
| *Euflavivirus* | Tembusu virus | *Orthoflavivirus tembusu* | KF826767 | Tembusu virus isolate TMUV-SDHS, |
| *Euflavivirus* | Tembusu virus | *Orthoflavivirus tembusu* | KJ782378 | Tembusu virus strain Du/CH/LSD/110128 P50, |

| Genus | Virus_name | Species_name | Accession_number | Description |
|---|---|---|---|---|
| *Euflavivirus* | Tembusu virus | *Orthoflavivirus tembusu* | KM102539 | Tembusu virus strain AH-F10 from China, |
| *Euflavivirus* | Tembusu virus | *Orthoflavivirus tembusu* | KX686573 | Tembusu virus isolate PY-2014, |
| *Euflavivirus* | Tembusu virus | *Orthoflavivirus tembusu* | KX977554 | Tembusu virus strain NCLT230, |
| *Euflavivirus* | Tembusu virus | *Orthoflavivirus tembusu* | MH414569 | Tembusu virus isolate MM1775, |
| *Euflavivirus* | Tembusu virus | *Orthoflavivirus tembusu* | MK907880 | Tembusu virus isolate GA, |
| *Euflavivirus* | Tembusu virus | *Orthoflavivirus tembusu* | MN747003 | Tembusu virus strain TP1906, |
| *Euflavivirus* | Tembusu virus | *Orthoflavivirus tembusu* | MT708901 | Tembusu virus isolate DTMUV/Goose/CHN/2019/AQ-19 |
| *Euflavivirus* | Tembusu virus | *Orthoflavivirus tembusu* | MT951412 | Tembusu virus strain BZ0162, |
| *Euflavivirus* | Tembusu virus | *Orthoflavivirus tembusu* | MW143073 | Tembusu virus isolate SCS01, |
| *Euflavivirus* | Tembusu virus | *Orthoflavivirus tembusu* | MW821486 | Tembusu virus isolate NTUC225/20, |
| *Euflavivirus* | Tembusu virus | *Orthoflavivirus tembusu* | MZ574097 | Tembusu virus isolate P4, |
| *Euflavivirus* | Tembusu virus | *Orthoflavivirus tembusu* | OK315313 | Tembusu virus isolate P4-2a, |
| *Euflavivirus* | Tembusu virus | *Orthoflavivirus tembusu* | PQ154625 | Tembusu virus isolate AVDDD05-CP001/TH2023, |
| *Euflavivirus* | Tembusu virus | *Orthoflavivirus tembusu* | PQ699974 | Tembusu virus isolate HZ2024/19, |
| *Euflavivirus* | Tembusu virus | *Orthoflavivirus tembusu* | PV541690 | Tembusu virus strain AHWH, |
| *Euflavivirus* | Tyuleniy virus | *Orthoflavivirus tyuleniyense* | KF815939 | Tyuleniy virus strain LEIV-6C polyprotein gene, |
| *Euflavivirus* | Uganda S virus | *Orthoflavivirus ugandaense* | DQ859065 | Uganda S virus polyprotein gene, |
| *Euflavivirus* | Usutu virus | *Orthoflavivirus usutuense* | KJ438715 | Usutu virus strain V66, |
| *Euflavivirus* | Usutu virus | *Orthoflavivirus usutuense* | KJ438722 | Usutu virus strain V256, |
| *Euflavivirus* | Usutu virus | *Orthoflavivirus usutuense* | KJ438731 | Usutu virus strain V43, |
| *Euflavivirus* | Usutu virus | *Orthoflavivirus usutuense* | KJ438766 | Usutu virus strain V321, |
| *Euflavivirus* | Usutu virus | *Orthoflavivirus usutuense* | KJ438769 | Usutu virus strain V20, |
| *Euflavivirus* | Usutu virus | *Orthoflavivirus usutuense* | KJ438776 | Usutu virus strain V68, |
| *Euflavivirus* | Usutu virus | *Orthoflavivirus usutuense* | KJ438779 | Usutu virus strain V191, |
| *Euflavivirus* | Usutu virus | *Orthoflavivirus usutuense* | KU664608 | Usutu virus clone ME/2015/ED-A, |
| *Euflavivirus* | Usutu virus | *Orthoflavivirus usutuense* | KX601690 | Usutu virus isolate USUV-HautRhin7315/France/2015, |
| *Euflavivirus* | Usutu virus | *Orthoflavivirus usutuense* | KY426754 | Usutu virus strain V466, |
| *Euflavivirus* | Usutu virus | *Orthoflavivirus usutuense* | KY426767 | Usutu virus strain V540, |
| *Euflavivirus* | Usutu virus | *Orthoflavivirus usutuense* | MF063042 | Usutu virus isolate AT-N211/16, |
| *Euflavivirus* | Usutu virus | *Orthoflavivirus usutuense* | OR141603 | Usutu virus isolate ED-I-178/21, |
| *Euflavivirus* | Usutu virus | *Orthoflavivirus usutuense* | OZ308550 | Usutu virus isolate USUV/NL/2022/AS20221974 genome |
| *Euflavivirus* | Usutu virus | *Orthoflavivirus usutuense* | OZ308552 | Usutu virus isolate USUV/NL/2022/22TV2828 genome |
| *Euflavivirus* | Usutu virus | *Orthoflavivirus usutuense* | OZ308554 | Usutu virus isolate USUV/NL/2017/AS201700198 |
| *Euflavivirus* | Usutu virus | *Orthoflavivirus usutuense* | OZ308618 | Usutu virus isolate USUV/NL/2019/19TV0329_S genome |
| *Euflavivirus* | Usutu virus | *Orthoflavivirus usutuense* | OZ308622 | Usutu virus isolate USUV/NL/2018/18TV0924 genome |
| *Euflavivirus* | Usutu virus | *Orthoflavivirus usutuense* | OZ416062 | Usutu virus isolate USUV/NL/2023/AS20230693 genome |
| *Euflavivirus* | Usutu virus | *Orthoflavivirus usutuense* | PP104387 | Usutu virus isolate TE.25185.1.2, |
| *Euflavivirus* | Sal Vieja virus | *Orthoflavivirus viejaense* | PZ324385 | Sal Vieja virus strain 38TWM-106, |
| *Euflavivirus* | Wesselsbron virus | *Orthoflavivirus wesselsbronense* | JN226796 | Wesselsbron virus from South Africa, |
| *Euflavivirus* | Wesselsbron virus | *Orthoflavivirus wesselsbronense* | KY056256 | Wesselsbron virus strain WSLV-IP248525/SEN/2013, |
| *Euflavivirus* | Wesselsbron virus | *Orthoflavivirus wesselsbronense* | KY056258 | Wesselsbron virus strain WSLV-IP259570/SEN/2013, |
| *Euflavivirus* | Wesselsbron virus | *Orthoflavivirus wesselsbronense* | ON157055 | Wesselsbron virus isolate UVRI5236 polyprotein gene, |
| *Euflavivirus* | Wesselsbron virus | *Orthoflavivirus wesselsbronense* | PP333214 | Wesselsbron virus isolate M5937-UG-2018, |
| *Euflavivirus* | Wesselsbron virus | *Orthoflavivirus wesselsbronense* | PP445079 | Wesselsbron virus isolate HD88963_SN_1988, |
| *Euflavivirus* | Wesselsbron virus | *Orthoflavivirus wesselsbronense* | PP445080 | Wesselsbron virus isolate ArD92269_SN_1988, |
| *Euflavivirus* | Wesselsbron virus | *Orthoflavivirus wesselsbronense* | PP445084 | Wesselsbron virus isolate ArD142157_SN_1995, |
| *Euflavivirus* | Wesselsbron virus | *Orthoflavivirus wesselsbronense* | PP445086 | Wesselsbron virus isolate ArD142143_SN_1995, |
| *Euflavivirus* | Wesselsbron virus | *Orthoflavivirus wesselsbronense* | PP445087 | Wesselsbron virus isolate ArA22079B_CI_1984, |
| *Euflavivirus* | Wesselsbron virus | *Orthoflavivirus wesselsbronense* | PP445088 | Wesselsbron virus isolate ArA23495_CI_1987, |
| *Euflavivirus* | Wesselsbron virus | *Orthoflavivirus wesselsbronense* | PP445089 | Wesselsbron virus isolate ArD65233_SN_1985, |
| *Euflavivirus* | Wesselsbron virus | *Orthoflavivirus wesselsbronense* | PP445090 | Wesselsbron virus isolate ArD90416_SN_1988, |
| *Euflavivirus* | Wesselsbron virus | *Orthoflavivirus wesselsbronense* | PP445091 | Wesselsbron virus isolate ArD90431_SN_1988, |
| *Euflavivirus* | Wesselsbron virus | *Orthoflavivirus wesselsbronense* | PP445092 | Wesselsbron virus isolate ArD90541_SN_1988, |
| *Euflavivirus* | Wesselsbron virus | *Orthoflavivirus wesselsbronense* | PP445097 | Wesselsbron virus isolate ArD140179_MR_1995, |
| *Euflavivirus* | Wesselsbron virus | *Orthoflavivirus wesselsbronense* | PP445098 | Wesselsbron virus isolate ArD140162_MR_1995, |
| *Euflavivirus* | Wesselsbron virus | *Orthoflavivirus wesselsbronense* | PP445099 | Wesselsbron virus isolate ArD140187_MR_1995, |
| *Euflavivirus* | Wesselsbron virus | *Orthoflavivirus wesselsbronense* | PP445101 | Wesselsbron virus isolate ArD142730_SN_1995, |
| *Euflavivirus* | Wesselsbron virus | *Orthoflavivirus wesselsbronense* | PP761170 | Wesselsbron strain RSA/OBP/WBV/LAV |
| *Euflavivirus* | Yaoundé virus | *Orthoflavivirus yaoundeense* | EU082199 | Yaounde virus strain Dak Ar Y276 polyprotein gene, |
| *Euflavivirus* | Yokose virus | *Orthoflavivirus yokoseense* | MH051229 | Yokose virus strain XYBX1332, |
| *Euflavivirus* | Yokose virus | *Orthoflavivirus yokoseense* | PZ324388 | Yokose virus strain Oita 36, |
| *Euflavivirus* | Zika virus | *Orthoflavivirus zikaense* | KU681081 | Zika virus isolate Zika virus/H.sapiens- |
| *Euflavivirus* | Zika virus | *Orthoflavivirus zikaense* | KU681082 | Zika virus isolate Zika virus/H.sapiens-tc/PHL/2012/CPC- |
| *Euflavivirus* | Zika virus | *Orthoflavivirus zikaense* | KX156776 | Zika virus strain ZIKV/Homo sapiens/PAN/CDC- |
| *Euflavivirus* | Zika virus | *Orthoflavivirus zikaense* | KX262887 | Zika virus isolate 103451, |
| *Euflavivirus* | Zika virus | *Orthoflavivirus zikaense* | KX601167 | Zika virus strain ZIKV/Aedes sp./MYS/P6-740/1966, |
| *Euflavivirus* | Zika virus | *Orthoflavivirus zikaense* | KX673530 | Zika virus isolate PHE_semen_Guadeloupe, |
| *Euflavivirus* | Zika virus | *Orthoflavivirus zikaense* | MG758785 | Zika virus isolate 41525, |
| *Euflavivirus* | Zika virus | *Orthoflavivirus zikaense* | MH061884 | Zika virus strain ZIKV/Macaca |
| *Euflavivirus* | Zika virus | *Orthoflavivirus zikaense* | MH061899 | Zika virus strain ZIKV/Macaca |
| *Euflavivirus* | Zika virus | *Orthoflavivirus zikaense* | MH061901 | Zika virus strain ZIKV/Macaca |
| *Euflavivirus* | Zika virus | *Orthoflavivirus zikaense* | MH675619 | Zika virus isolate CTS-30-16p, |
| *Euflavivirus* | Zika virus | *Orthoflavivirus zikaense* | MT377503 | Zika virus isolate 15-1144, |
| *Euflavivirus* | Zika virus | *Orthoflavivirus zikaense* | MT507050 | Zika virus isolate S-542/Yucatan/2017 clone 6A, |
| *Euflavivirus* | Zika virus | *Orthoflavivirus zikaense* | MT636065 | Zika virus isolate ARCB116141, |
| *Euflavivirus* | Zika virus | *Orthoflavivirus zikaense* | MW143020 | Zika virus strain MR766, |
| *Euflavivirus* | Zika virus | *Orthoflavivirus zikaense* | ON142673 | Zika virus isolate AB3234, |
| *Euflavivirus* | Zika virus | *Orthoflavivirus zikaense* | PP316140 | Zika virus isolate Zika virus/human/Haiti/36/2021, |
| *Euflavivirus* | Zika virus | *Orthoflavivirus zikaense* | PP626403 | Zika virus isolate Z5-ERC-R2, |
| *Euflavivirus* | Zika virus | *Orthoflavivirus zikaense* | PQ129535 | Zika virus isolate ZIKV/H.sapiens/Brazil/PE243/2015 |
| *Euflavivirus* | Zika virus | *Orthoflavivirus zikaense* | PQ869256 | Zika virus isolate Sen/Thai-NS3, |

| Genus | Virus_name | Species_name | Accession_number | Description |
|---|---|---|---|---|
| *Euflavivirus* | Tick-borne encephalitis virus | *Orthoflavivirus zilberi* | JQ825159 | Tick-borne encephalitis virus strain Primorye-274, |
| *Euflavivirus* | Tick-borne encephalitis virus | *Orthoflavivirus zilberi* | JQ825151 | Tick-borne encephalitis virus strain Spassk-72, |
| *Euflavivirus* | Tick-borne encephalitis virus | *Orthoflavivirus zilberi* | KP844726 | Tick-borne encephalitis virus strain Birobidzhan 1354 |
| *Euflavivirus* | Tick-borne encephalitis virus | *Orthoflavivirus zilberi* | AB753012 | Tick-borne encephalitis virus RNA, strain: Oshima |
| *Euflavivirus* | Tick-borne encephalitis virus | *Orthoflavivirus zilberi* | JQ825147 | Tick-borne encephalitis virus strain Shkotovo-94, |
| *Euflavivirus* | Tick-borne encephalitis virus | *Orthoflavivirus zilberi* | KU761574 | Tick-borne encephalitis virus isolate Primorye-1284 |
| *Euflavivirus* | Tick-borne encephalitis virus | *Orthoflavivirus zilberi* | KJ914683 | Tick-borne encephalitis virus strain Tomsk-M202 |
| *Euflavivirus* | Tick-borne encephalitis virus | *Orthoflavivirus zilberi* | KJ739731 | Tick-borne encephalitis virus strain Tomsk-M83 |
| *Euflavivirus* | Tick-borne encephalitis virus | *Orthoflavivirus zilberi* | JQ650522 | Tick-borne encephalitis virus isolate MDJ01 |
| *Euflavivirus* | Tick-borne encephalitis virus | *Orthoflavivirus zilberi* | PV183805 | Tick-borne encephalitis virus isolate |
| *Euflavivirus* | Tick-borne encephalitis virus | *Orthoflavivirus zilberi* | JQ825148 | Tick-borne encephalitis virus strain Primorye-82, |
| *Euflavivirus* | Tick-borne encephalitis virus | *Orthoflavivirus zilberi* | MT681749 | Tick-borne encephalitis virus strain M442, |
| *Euflavivirus* | Tick-borne encephalitis virus | *Orthoflavivirus zilberi* | PX467451 | Tick-borne encephalitis virus isolate Krasnyi Yar-9 |
| *Euflavivirus* | Tick-borne encephalitis virus | *Orthoflavivirus zilberi* | KC422667 | Tick-borne encephalitis virus strain Zabaikalye 30-00 |
| *Euflavivirus* | Tick-borne encephalitis virus | *Orthoflavivirus zilberi* | PX794532 | Tick-borne encephalitis virus strain NMGHMY15, |
| *Euflavivirus* | Tick-borne encephalitis virus | *Orthoflavivirus zilberi* | JQ825162 | Tick-borne encephalitis virus strain Primorye-437, |
| *Euflavivirus* | Tick-borne encephalitis virus | *Orthoflavivirus zilberi* | JX498939 | Tick-borne encephalitis virus strain 205 |
| *Euflavivirus* | Tick-borne encephalitis virus | *Orthoflavivirus zilberi* | JQ825163 | Tick-borne encephalitis virus strain Primorye-750, |
| *Euflavivirus* | Tick-borne encephalitis virus | *Orthoflavivirus zilberi* | KT001073 | Tick-borne encephalitis virus strain Lazo MP36 |
| *Euflavivirus* | Tick-borne encephalitis virus | *Orthoflavivirus zilberi* | KP844725 | Tick-borne encephalitis virus strain Chichagovka 1223 |
| *Classified fusiviruses* | | | | |
| *Fusivirus* | Cell fusing agent virus | *Orthoflavivirus junctionis* | KJ741267 | Cell fusing agent virus strain Galveston |
| *Fusivirus* | Cell fusing agent virus | *Orthoflavivirus junctionis* | LC496857 | Cell fusing agent virus 16GH83 RNA |
| *Fusivirus* | Cell fusing agent virus | *Orthoflavivirus junctionis* | M91671 | Flavivirus cell fusing agent polyprotein gene |
| *Fusivirus* | Cell fusing agent virus | *Orthoflavivirus junctionis* | MH237596 | Cell fusing agent virus strain 2 |
| *Fusivirus* | Cell fusing agent virus | *Orthoflavivirus junctionis* | MK860761 | Cell fusing agent virus isolate KPP |
| *Fusivirus* | Cell fusing agent virus | *Orthoflavivirus junctionis* | MZ972992 | Cell fusing agent virus strain Macapa_01 |
| *Fusivirus* | Cell fusing agent virus | *Orthoflavivirus junctionis* | MZ972993 | Cell fusing agent virus strain Macapa_04 |
| *Fusivirus* | Cell fusing agent virus | *Orthoflavivirus junctionis* | MZ972994 | Cell fusing agent virus strain Macapa_02 |
| *Fusivirus* | Cell fusing agent virus | *Orthoflavivirus junctionis* | LR596014 | Cell fusing agent virus gene for , |
| *Fusivirus* | Cell fusing agent virus | *Orthoflavivirus junctionis* | OR479690 | Cell fusing agent virus strain Ae11 |
| *Fusivirus* | Cell fusing agent virus | *Orthoflavivirus junctionis* | OR479689 | Cell fusing agent virus strain Ae10 |
| *Fusivirus* | Cell fusing agent virus | *Orthoflavivirus junctionis* | LR694077 | Cell fusing agent virus |
| *Fusivirus* | Cell fusing agent virus | *Orthoflavivirus junctionis* | OR479687 | Cell fusing agent virus strain Ae6 |
| *Fusivirus* | Cell fusing agent virus | *Orthoflavivirus junctionis* | KJ476731 | Cell fusing agent virus isolate Mex AR269 |
| *Fusivirus* | Cell fusing agent virus | *Orthoflavivirus junctionis* | KU936054 | Cell fusing agent virus strain Aag2 Bristol |
| *Fusivirus* | Cell fusing agent virus | *Orthoflavivirus junctionis* | LR694072 | Cell fusing agent virus |
| *Fusivirus* | Cell fusing agent virus | *Orthoflavivirus junctionis* | LR694075 | Cell fusing agent virus gene for , |
| *Fusivirus* | Cell fusing agent virus | *Orthoflavivirus junctionis* | MH310082 | Cell fusing agent virus isolate CC_A_CFAV |
| *Fusivirus* | Cell fusing agent virus | *Orthoflavivirus junctionis* | OQ305237 | Cell fusing agent virus isolate KLF47 |
| *Fusivirus* | Cell fusing agent virus | *Orthoflavivirus junctionis* | LR694074 | Cell fusing agent virus gene for , |
| *Other classified subgenera* | | | | |
| *Crangovirus* | Crangon crangon flavivirus | *Orthoflavivirus aphei* | MK473878 | Crangon crangon flavivirus |
| *Crangovirus* | Crangon crangon flavivirus | *Orthoflavivirus aphei* | PP215320 | Crangon crangon flavivirus isolate 02/HH/201811 |
| *Crangovirus* | Crangon crangon flavivirus | *Orthoflavivirus aphei* | PP215319 | Crangon crangon flavivirus isolate 01/HH/201811 |
| *Unclassified euflaviviruses (ICTV)* | | | | |
| *Euflavivirus* | Barkedji virus | Unclass. | KC496020 | Barkedji virus isolate 363/11 |
| *Euflavivirus* | Barkedji virus | Unclass. | LC497470 | Barkedji virus zmq17L31 RNA |
| *Euflavivirus* | Barkedji virus | Unclass. | MG214905 | Barkedji virus strain SQU29/Oman/2016 |
| *Euflavivirus* | Barkedji virus | Unclass. | MG214906 | Barkedji virus strain SQU38/Oman/2016 |
| *Euflavivirus* | Chaoyang virus | Unclass. | FJ883471 | Chaoyang virus strain Deming polyprotein |
| *Euflavivirus* | Donggang virus | Unclass. | JQ086551 | Donggang virus isolate DG0909 |
| *Euflavivirus* | Donggang virus | Unclass. | MZ357894 | Donggang virus isolate 5 |
| *Euflavivirus* | Donggang virus | Unclass. | MZ357895 | Donggang virus isolate 3 |
| *Euflavivirus* | Donggang virus | Unclass. | MZ357896 | Donggang virus isolate 5-3 |
| *Euflavivirus* | Donggang virus | Unclass. | MZ357897 | Donggang virus isolate 5.1-81 |
| *Euflavivirus* | Donggang virus | Unclass. | MZ357898 | Donggang virus isolate 5.82-107 |
| *Euflavivirus* | Donggang virus | Unclass. | MZ357899 | Donggang virus isolate 3.1-102 |
| *Euflavivirus* | Fitzroy river virus | Unclass. | KM361634 | Fitzroy River Virus strain K73884 |
| *Euflavivirus* | Ilomantsi virus | Unclass. | KC692067 | Ilomantsi virus isolate M0724 |
| *Euflavivirus* | Kampung Karu virus | Unclass. | KY320648 | Kampung Karu virus isolate SWK_P44 |
| *Euflavivirus* | Lammi virus | Unclass. | FJ606789 | Lammi virus |
| *Euflavivirus* | Lammi virus | Unclass. | KC692068 | Lammi virus isolate M0719 |
| *Euflavivirus* | Lammi virus | Unclass. | PP329310 | Lammi virus isolate KHAM-T22912 |
| *Euflavivirus* | Long Pine virus | Unclass. | KY290256 | Long Pine Key virus strain EVG 5-72 |
| *Euflavivirus* | Marisma mosquito virus | Unclass. | KY347801 | Marisma mosquito virus isolate HU 4528/07 |
| *Euflavivirus* | Marisma mosquito virus | Unclass. | MF139576 | Marisma mosquito virus strain HU4528/07 |
| *Euflavivirus* | Nanay virus | Unclass. | MF139575 | Nanay virus strain PRD316/PER/09 |
| *Euflavivirus* | Nhumirim virus | Unclass. | KJ210048 | Nhumirim virus strain BrMS-MQ10 |
| *Euflavivirus* | Nounané virus | Unclass. | EU159426 | Nounane virus polyprotein mRNA |
| *Euflavivirus* | Paraiso Escondido virus | Unclass. | KJ152564 | Paraiso Escondido virus isolate Ecuador2012 |
| *Euflavivirus* | Spondweni virus | Unclass. | DQ859064 | Spondweni virus strain SM-6 V-1 |
| *Euflavivirus* | T'hó virus | Unclass. | EU879061 | T'Ho virus strain T'Ho-Mex07 |
| More divergent dengue viruses | | | | |
| *Euflavivirus* | DENV 1 | Divergent | KR919820 | Dengue virus isolate Brun2014 |
| *Euflavivirus* | DENV 1 | Sylvatic | EF457905 | Dengue virus type 1 isolate P72-1244 |
| *Euflavivirus* | DENV 2 | Divergent | FJ467493 | Dengue virus 2 isolate DKD811 |
| *Euflavivirus* | DENV 2 | Divergent | KX274130 | Dengue virus type 2 strain QML22 |
| *Euflavivirus* | DENV 2 | ylvatic | EF105379 | Dengue virus type 2 isolate P8-1407 polyprotein gene, |

| Genus | Virus_name | Species_name | Accession_number | Description |
|---|---|---|---|---|
| *Euflavivirus* | DENV 4 | ylvatic | EF457906 | Dengue virus type 4 isolate P75-215 |
| *Euflavivirus* | DENV 4 | ylvatic | JF262779 | Dengue virus 4 isolate P75-514 |
| *Euflavivirus* | DENV 4 | ylvatic | JF262780 | Dengue virus 4 isolate P73-1120 |
| *Euflavivirus* | DENV 4 | Divergent | OK605599 | Dengue virus 4 isolate DKE-121 |
| *Unclassified fusiviruses (ICTV)* | | | | |
| *Fusivirus* | Anopheles flavivirus | Unclass. | KX148546 | Ano Anopheles flavivirus variant 1 |
| *Fusivirus* | Anopheles flavivirus | Unclass. | KX148547 | A Anopheles flavivirus variant 2 |
| *Fusivirus* | Aedes flavivirus | Unclass. | AB488408 | Aed Aedes flavivirus, strain: Narita-21 |
| *Fusivirus* | Aedes flavivirus | Unclass. | KC181923 | Aedes flavivirus strain AEFV-SPFLD-MO-2011-MP6 |
| *Fusivirus* | Aedes flavivirus | Unclass. | KJ741266 | Aedes flavivirus strain Bangkok |
| *Fusivirus* | Aedes flavivirus | Unclass. | LC770118 | Aedes flavivirus AeFV/AAL/Gunma-JP/2019 RNA |
| *Fusivirus* | Aedes flavivirus | Unclass. | MT254427 | Aedes flavivirus strain TC4A8_18-9L-Y-T-Aea-B-1-1 |
| *Fusivirus* | Aedes flavivirus | Unclass. | MT577804 | Aedes flavivirus strain AEFV/Ticino/2019 |
| *Fusivirus* | Aedes flavivirus | Unclass. | OR098296 | Aedes flavivirus strain Sasso Marconi 2011 |
| *Fusivirus* | Aedes flavivirus | Unclass. | PQ223729 | Aedes flavivirus isolate AF-hnwzs |
| *Fusivirus* | Aedes flavivirus | Unclass. | PV165522 | Aedes flavivirus isolate JH2YW |
| *Fusivirus* | Aedes flavivirus | Unclass. | PZ303603 | Aedes flavivirus strain AEFV-SPFLD-MO-2011-MP6 |
| *Fusivirus* | Cuacua virus | Unclass. | KX245154 | Cuacua virus isolate MZ2014-Cv polyprotein |
| *Fusivirus* | Culex theileri flavivirus | Unclass. | HE574574 | Culex theileri flavivirus RP-2011 |
| *Fusivirus* | Culex theileri flavivirus | Unclass. | MF153378 | Culex theileri flavivirus |
| *Fusivirus* | Culex flavivirus | Unclass. | AB377213 | Culex flavivirus, strain: NIID-21-2 |
| *Fusivirus* | Culex flavivirus | Unclass. | AB701766 | Culex flavivirus RNA, strain: Toyama71 |
| *Fusivirus* | Culex flavivirus | Unclass. | AB701770 | Culex flavivirus RNA, strain: Toyama41 |
| *Fusivirus* | Culex flavivirus | Unclass. | AB701772 | Culex flavivirus RNA, strain: Toyama740 |
| *Fusivirus* | Culex flavivirus | Unclass. | AB701775 | Culex flavivirus RNA, strain: Toyama1431 |
| *Fusivirus* | Culex flavivirus | Unclass. | AB701776 | Culex flavivirus RNA, strain: Toyama2627 |
| *Fusivirus* | Culex flavivirus | Unclass. | EU879060 | Culex flavivirus strain CxFV-Mex07 |
| *Fusivirus* | Culex flavivirus | Unclass. | FJ502995 | Culex flavivirus strain HOU24518 |
| *Fusivirus* | Culex flavivirus | Unclass. | FJ663034 | Culex flavivirus isolate Iowa07 |
| *Fusivirus* | Culex flavivirus | Unclass. | GQ165808 | Culex flavivirus strain Uganda08 |
| *Fusivirus* | Culex flavivirus | Unclass. | JQ308186 | Culex flavivirus isolate HLD102 |
| *Fusivirus* | Culex flavivirus | Unclass. | JQ308187 | Culex flavivirus isolate HLD105 |
| *Fusivirus* | Culex flavivirus | Unclass. | LC504568 | Culex flavivirus 16GH36 |
| *Fusivirus* | Culex flavivirus | Unclass. | MH188006 | Culex flavivirus strain CFV/San Francisco |
| *Fusivirus* | Culex flavivirus | Unclass. | MH719098 | Culex flavivirus isolate CxFV-T1123 |
| *Fusivirus* | Culex flavivirus | Unclass. | MN294938 | Culex flavivirus isolate SA-JD-AJ-AJ-16-1-C7 |
| *Fusivirus* | Culex flavivirus | Unclass. | MT197496 | UNVERIFIED: Culex flavivirus strain 111 |
| *Fusivirus* | Culex flavivirus | Unclass. | OK413947 | Culex flavivirus strain CxFV_Mygg_341 |
| *Fusivirus* | Culex flavivirus | Unclass. | OM914715 | Culex flavivirus isolate NX1908 |
| *Fusivirus* | Culex flavivirus | Unclass. | PV165384 | Culex flavivirus isolate 1_RNA/Culex/China/PEKW/2022 |
| *Fusivirus* | Culex flavivirus | Unclass. | PZ305743 | Culex flavivirus strain HOU24518 |
| *Fusivirus* | Culiseta flavivirus | Unclass. | KT599442 | Culiseta flavivirus strain 502-13 |
| *Fusivirus* | Hanko virus | Unclass. | JQ268258 | Hanko virus polyprotein gene |
| *Fusivirus* | Hanko virus | Unclass. | ON949927 | MAG: Hanko virus isolate FIN/U-2018/94 |
| *Fusivirus* | Hanko virus | Unclass. | ON949928 | MAG: Hanko virus isolate FIN/U-2018/95 |
| *Fusivirus* | Hanko virus | Unclass. | ON949929 | MAG: Hanko virus isolate FIN/U-2018/96 |
| *Fusivirus* | Hanko virus | Unclass. | ON949930 | MAG: Hanko virus isolate FIN/U-2018/97 |
| *Fusivirus* | Kamiti River virus | Unclass. | AY149905 | Kamiti River virus isolate SR-82 |
| *Fusivirus* | La Tina virus | Unclass. | KY320649 | La Tina virus isolate 49_LT96 |
| *Fusivirus* | La Tina virus | Unclass. | PV165523 | La Tina virus isolate LHYW |
| *Fusivirus* | Mercadeo virus | Unclass. | KP688057 | Mercadeo virus isolate ER-M31 |
| *Fusivirus* | Mercadeo virus | Unclass. | KP688058 | Mercadeo virus isolate ER-M10 |
| *Fusivirus* | Mosquito 2 flavivirus | Unclass. | AB981186 | Mosquito flavivirus |
| *Fusivirus* | Mosquito 1 flavivirus | Unclass. | KC464457 | Mosquito flavivirus isolate LSFlaviV-A20-09 |
| *Fusivirus* | Mosquito 1 flavivirus | Unclass. | KX652375 | Mosquito flavivirus isolate 94_1 |
| *Fusivirus* | Mosquito 1 flavivirus | Unclass. | KX652377 | Mosquito flavivirus isolate 65_02 |
| *Fusivirus* | Mosquito 2 flavivirus | Unclass. | KX652378 | Mosquito flavivirus isolate 599 |
| *Fusivirus* | Mosquito 2 flavivirus | Unclass. | MZ821064 | Mosquito flavivirus strain YN15-283-02 |
| *Fusivirus* | Nakiwogo virus | Unclass. | GQ165809 | Nakiwogo virus strain Uganda08 |
| *Fusivirus* | Nienokoue virus | Unclass. | JQ957875 | Nienokoue virus isolate B51/CI/2004 |
| *Fusivirus* | Palm Creek virus | Unclass. | KC505248 | Palm Creek virus isolate 56 |
| *Fusivirus* | Parramatta River virus | Unclass. | KT192549 | Parramatta River virus isolate 92-B115745 |
| *Fusivirus* | Quang Binh virus | Unclass. | FJ644291 | Quang Binh virus isolate VN180 |
| *Fusivirus* | Xīshuāngbǎnnà aedes flavivirus | Unclass. | KU201526 | Xishuangbanna aedes flavivirus |
| *Unclassified euflaviviruses (NCBI)* | | | | |
| *Euflavivirus* | Alpine chamois encephalitis virus | Unclass. | PP782054 | Alpine chamois encephalitis virus strain cds |
| *Euflavivirus* | Alpine chamois encephalitis virus | Unclass. | PP782055 | Alpine chamois encephalitis virus strain cds |
| *Euflavivirus* | Alpine chamois encephalitis virus | Unclass. | PP795456 | Alpine chamois encephalitis virus isolate Tick\63100- |
| *Euflavivirus* | Alpine chamois encephalitis virus | Unclass. | PP795457 | Alpine chamois encephalitis virus isolate |
| *Euflavivirus* | Alpine chamois encephalitis virus | Unclass. | KU308380 | Bamaga virus isolate CY4270 |
| *Euflavivirus* | Alpine chamois encephalitis virus | Unclass. | MG587038 | Binjari virus isolate BFTA20 polyprotein gene, |
| *Euflavivirus* | Alpine chamois encephalitis virus | Unclass. | MW032264 | Chinese soft-shelled turtle flavivirus isolate HZ-2017 |
| *Euflavivirus* | Binjari virus | Unclass. | PQ150542 | Hebei orthoflavivirus isolate Hebei_067 |
| *Euflavivirus* | Chinese soft-shelled turtle | Unclass. | PV778045 | Hebei orthoflavivirus isolate AH-2023-A7 polyprotein |
| *Euflavivirus* | Héběi orthoflavivirus | Unclass. | PZ000270 | Hebei orthoflavivirus isolate 2_29574 |
| *Euflavivirus* | Héběi orthoflavivirus | Unclass. | PZ000271 | Hebei orthoflavivirus isolate JZ-3_2838 |
| *Euflavivirus* | Héběi orthoflavivirus | Unclass. | KF815940 | Kama virus strain LEIV-20776Tat, |
| *Euflavivirus* | Héběi orthoflavivirus | Unclass. | MT812695 | Kama virus strain KAMV/M16-Omsk-1983 |
| *Euflavivirus* | Kama virus | Unclass. | MT812696 | Kama virus strain KAMV/M48-Omsk-1983 |

| Genus | Virus_name | Species_name | Accession_number | Description |
|---|---|---|---|---|
| *Euflavivirus* | Kama virus | Unclass. | PV021569 | Lunay virus strain AR826399/LUNYV, |
| *Euflavivirus* | Kama virus | Unclass. | LC582740 | Mpulungu flavivirus ZT17-1078 RNA, |
| *Euflavivirus* | Lunay virus | Unclass. | LC567151 | Psorophora flavivirus bvmq18-51 RNA, |
| *Euflavivirus* | Mpulungu flavivirus | Unclass. | LC902872 | Psorophora flavivirus P12_37-degree-adapted RNA, |
| *Euflavivirus* | Psorophora flavivirus | Unclass. | LC657549 | Saruyama virus 18HKR14, |
| *Euflavivirus* | Psorophora flavivirus | Unclass. | OP503546 | Trionyx sinensis flavivirus polyprotein (YH2016000001) |
| *Euflavivirus* | Saruyama virus | Unclass. | OQ238819 | Trionyx sinensis flavivirus isolate YH2018, |
| *Euflavivirus* | Trionyx sinensis flavivirus | Unclass. | OP699738 | Xinyang flavivirus strain HUNAU202107, |
| *Euflavivirus* | Trionyx sinensis flavivirus | Unclass. | PV690395 | Yanbian tick orthoflavivirus isolate Tick_HLJ21, |
| *Unclassified fusiviruses (NCBI)* | | | | |
| *Fusivirus* | Accesaty virus | Unclass. | OQ915774 | Accesaty virus isolate F1416 |
| *Fusivirus* | Aedes flavivirus | Unclass. | LC536088 | Aedes flavivirus AEFV/MQ/29/Bogor/2017 |
| *Fusivirus* | Aedes flavivirus | Unclass. | MK251047 | Aedes flavivirus strain KRD32 |
| *Fusivirus* | Aedes flavivirus | Unclass. | OQ915866 | Aedes flavivirus isolate F1161 |
| *Fusivirus* | Aedes flavivirus | Unclass. | LC775064 | Aedes flavivirus AeAl-2/AEFV |
| *Fusivirus* | Aedes notoscriptus flavivirus | Unclass. | MZ358857 | Aedes notoscriptus flavivirus isolate 207252, |
| *Fusivirus* | Aedes notoscriptus flavivirus | Unclass. | MZ358856 | Aedes notoscriptus flavivirus isolate 207218, |
| *Fusivirus* | Aguas Brancas virus | Unclass. | OQ749789 | Aguas Brancas virus isolate Ar839941 |
| *Fusivirus* | Aguas Brancas virus | Unclass. | OQ749788 | Aguas Brancas virus isolate Ar812610/02 |
| *Fusivirus* | Aguas Brancas virus | Unclass. | OQ749787 | Aguas Brancas virus isolate Ar812610/01 |
| *Fusivirus* | Calbertado virus | Unclass. | MH188007 | Culex Flavi-like virus strain CFVL/San Francisco |
| *Fusivirus* | Calbertado virus | Unclass. | KX669685 | Calbertado virus isolate SK-90 |
| *Fusivirus* | Calbertado virus | Unclass. | MW434106 | Calbertado virus isolate CMS001_032_ALCO |
| *Fusivirus* | Calbertado virus | Unclass. | KX669682 | Calbertado virus isolate Cal_mos_2 |
| *Fusivirus* | Calbertado virus | Unclass. | KX669684 | Calbertado virus isolate SK-43 |
| *Fusivirus* | Calbertado virus | Unclass. | KX669683 | Calbertado virus isolate 42-81 |
| *Fusivirus* | Calbertado virus | Unclass. | KX669689 | Calbertado virus isolate CALper |
| *Fusivirus* | Calbertado virus | Unclass. | KX669687 | Calbertado virus isolate 4a2 |
| *Fusivirus* | Calbertado virus | Unclass. | KX669688 | Calbertado virus isolate 44-13 |
| *Fusivirus* | Calbertado virus | Unclass. | KX669686 | Calbertado virus isolate 42-15 |
| *Fusivirus* | Culex theileri flavivirus | Unclass. | HE574573 | Culex theileri flavivirus RP-2011 |
| *Fusivirus* | Falli virus | Unclass. | MN567479 | Falli virus isolate Ferlo |
| *Fusivirus* | Flavivirus virus | Unclass. | OQ588812 | Flavivirus sp. isolate M138 |
| *Fusivirus* | Haslams Creek virus | Unclass. | MF352617 | Haslams Creek virus isolate HaCV 115734 |
| *Fusivirus* | Karumba virus | Unclass. | KY460523 | Karumba virus isolate KRBV WBay2 |
| *Fusivirus* | Karumba virus | Unclass. | MF352615 | Karumba virus isolate KRBV Kim1 |
| *Fusivirus* | Karumba virus | Unclass. | KY460522 | Karumba virus isolate KRBV 1892 |
| *Fusivirus* | Kilpisjarvi flavivirus | Unclass. | ON949931 | Kilpisjarvi flavivirus isolate FIN/L-2018/90 |
| *Fusivirus* | Mac Peak virus | Unclass. | MF352616 | Mac Peak virus isolate McPV 150840 |
| *Fusivirus* | Mansonia flavivirus | Unclass. | LC567153 | Mansonia flavivirus bvmq18-1 RNA, |
| *Fusivirus* | Menghai flavivirus | Unclass. | KX907452 | Menghai flavivirus isolate MHAedFV1 |
| *Fusivirus* | Mosquito flavivirus | Unclass. | OL700075 | Mosquito flavivirus isolate XY107097 |
| *Fusivirus* | Mosquito flavivirus | Unclass. | LC513840 | Mosquito flavivirus 17CxNGK-Ctr-C6P3 |
| *Fusivirus* | Nienokoue virus | Unclass. | KX879625 | Nienokoue virus strain B21/CI/2004 |
| *Fusivirus* | Nienokoue virus | Unclass. | KX879626 | Nienokoue virus strain C64/CI/2004 |
| *Fusivirus* | Ochlerotatus caspius flavivirus | Unclass. | HF548540 | Ochlerotatus caspius flavivirus |
| *Fusivirus* | Ochlerotatus caspius flavivirus | Unclass. | KY345399 | Ochlerotatus caspius flavivirus isolate 1608 |
| *Fusivirus* | Ochlerotatus scapularis flavivirus | Unclass. | LC567152 | Ochlerotatus scapularis flavivirus bvmq18-25 RNA, |
| *Fusivirus* | Orthoflavivirus | Unclass. | PZ161043 | Orthoflavivirus sp. isolate Mos_EL_116 |
| *Fusivirus* | Orthoflavivirus | Unclass. | PZ161033 | Orthoflavivirus sp. isolate Mos_EL_107 |
| *Fusivirus* | Orthoflavivirus | Unclass. | PZ161045 | Orthoflavivirus sp. isolate Mos_EL_112 |
| *Fusivirus* | Orthoflavivirus | Unclass. | PZ161035 | Orthoflavivirus sp. isolate Mos_EL_110 |
| *Fusivirus* | Orthoflavivirus | Unclass. | PZ161042 | Orthoflavivirus sp. isolate Mos_EL_109 |
| *Fusivirus* | Orthoflavivirus | Unclass. | PZ161036 | Orthoflavivirus sp. isolate Mos_EL_105 |
| *Fusivirus* | Orthoflavivirus | Unclass. | PZ161040 | Orthoflavivirus sp. isolate Mos_EL_78 |
| *Fusivirus* | Orthoflavivirus | Unclass. | PZ161034 | Orthoflavivirus sp. isolate Mos_EL_75 |
| *Fusivirus* | Orthoflavivirus | Unclass. | PZ161037 | Orthoflavivirus sp. isolate Mos_EL_73 |
| *Fusivirus* | Orthoflavivirus | Unclass. | PZ161038 | Orthoflavivirus sp. isolate Mos_EL_72 |
| *Fusivirus* | Orthoflavivirus | Unclass. | PZ161039 | Orthoflavivirus sp. isolate Mos_EL_114 |
| *Fusivirus* | Orthoflavivirus | Unclass. | PZ161041 | Orthoflavivirus sp. isolate Mos_EL_106 |
| *Fusivirus* | Palm Creek virus | Unclass. | OQ915762 | Palm Creek virus isolate F1616 |
| *Fusivirus* | Palm Creek virus | Unclass. | OQ915761 | Palm Creek virus isolate F1392F1551 |
| *Fusivirus* | Palm Creek virus | Unclass. | OQ915759 | Palm Creek virus isolate F1392F1225 |
| *Fusivirus* | Palm Creek virus | Unclass. | OQ915760 | Palm Creek virus isolate F1392 |
| *Fusivirus* | Palm Creek virus | Unclass. | KT192550 | Palm Creek virus isolate K71061 |
| *Fusivirus* | Parramatta River virus | Unclass. | PP066163 | Parramatta River virus isolate mosPPEE |
| *Fusivirus* | Psorophora ferox flavivirus | Unclass. | PP946240 | Psorophora ferox flavivirus isolate Flavi |
| *Fusivirus* | Quang Binh virus | Unclass. | OQ915806 | Quang Binh virus isolate F1449F1466 |
| *Fusivirus* | Quang Binh virus | Unclass. | MH827524 | Quang Binh virus isolate JM17156/China/2017 |
| *Fusivirus* | Quang Binh virus | Unclass. | PQ376979 | Quang Binh virus isolate Ganzhou |
| *Fusivirus* | Quang Binh virus | Unclass. | OP087419 | Quang Binh virus isolate 21-7-SY-CXT-18 |
| *Fusivirus* | Quang Binh virus | Unclass. | OQ915805 | Quang Binh virus isolate F1449 |
| *Fusivirus* | Quang Binh virus | Unclass. | OQ915803 | Quang Binh virus isolate F1449F1108 |
| *Fusivirus* | Quang Binh virus | Unclass. | MH827523 | Quang Binh virus isolate LZ17046/China/2017 |
| *Fusivirus* | Quang Binh virus | Unclass. | OR479686 | Quang Binh virus strain Cg2 |
| *Fusivirus* | Quang Binh virus | Unclass. | PX205256 | Quang Binh virus isolate 2024-08 |
| *Fusivirus* | Quang Binh virus | Unclass. | MW452275 | Quang Binh virus strain YX594 |
| *Fusivirus* | Quang Binh virus | Unclass. | MG719525 | Quang Binh virus isolate DD1716 |
| *Fusivirus* | Quang Binh virus | Unclass. | OQ139647 | Quang Binh virus isolate GZ21m120 |
| *Fusivirus* | Quang Binh virus | Unclass. | PV010630 | Quang Binh virus isolate 490 Cx. ge 9/2/21 T6 |

| Genus | Virus_name | Species_name | Accession_number | Description |
|---|---|---|---|---|
| *Fusivirus* | Quang Binh virus | Unclass. | OQ139648 | Quang Binh virus isolate GZ21m167 |
| *Fusivirus* | Quang Binh virus | Unclass. | PV010628 | Quang Binh virus isolate 490 Cx. ge 9/2/21 T1 |
| *Fusivirus* | Quang Binh virus | Unclass. | OP087420 | Quang Binh virus isolate 21-9-YY-CXT-3 |
| *Fusivirus* | Quang Binh virus | Unclass. | MW246771 | Quang Binh virus strain JS,JA_H6_18-10E-JS-Cxt-C-2-1 |
| *Fusivirus* | Quang Binh virus | Unclass. | OQ139646 | Quang Binh virus isolate GZ21m081 |
| *Fusivirus* | Quang Binh virus | Unclass. | PQ376980 | Quang Binh virus isolate Jiujiang |
| *Fusivirus* | Quang Binh virus | Unclass. | OQ915802 | Quang Binh virus isolate F1407 |
| *Fusivirus* | Sabethes flavivirus | Unclass. | MH899446 | Sabethes flavivirus isolate Ms42 |
| *Fusivirus* | Serra gelada orthoflavivirus | Unclass. | PQ673810 | Serra pelada orthoflavivirus isolate AR862689/ORFV |
| *Fusivirus* | Tafomo virus | Unclass. | MZ202252 | Tafomo virus strain TV/B22-CI-2004 |
| *Other unclassified potential subgenera* | | | | |
| Unclass. | Cnidaria flavivirus | Unclass. | OX394137 | Cnidaria flavivirus polyprotein, |
| *Unclass.* | Hangzhou flavivirus | Unclass. | MZ209680 | Hangzhou flavivirus 3 isolate YWLFY92, |
| *Unclass.* | Harrimaniidae flavivirus | Unclass. | OX394149 | Harrimaniidae flavivirus polyprotein, |
| *Unclass.* | Sea-firefly flavivirus | Unclass. | OX394161 | Sea-firefly flavivirus |
| *Unclass.* | Tabanus flavivirus | Unclass. | LC540441 | Tabanus rufidens flavivirus 18HF18 |

**Table S3.** Publications describing currently unclassified orthoflaviviruses (download from PubMed, 27/08/2026; unformatted texr)

25. Pabbaraju K, Ho KC, Wong S, Fox JD, Kaplen B, Tyler S, Drebot M, Tilley PA. 2009. Surveillance of mosquito-borne viruses in Alberta using reverse transcription polymerase chain reaction with generic primers. J Med Entomol 46:640-8.
26. Huhtamo E, Putkuri N, Kurkela S, Manni T, Vaheri A, Vapalahti O, Uzcátegui NY. 2009. Characterization of a novel flavivirus from mosquitoes in northern europe that is related to mosquito-borne flaviviruses of the tropics. J Virol 83:9532-40.
27. Roiz D, Vázquez A, Seco MP, Tenorio A, Rizzoli A. 2009. Detection of novel insect flavivirus sequences integrated in Aedes albopictus (Diptera: Culicidae) in Northern Italy. Virol J 6:93.
28. Hoshino K, Isawa H, Tsuda Y, Sawabe K, Kobayashi M. 2009. Isolation and characterization of a new insect flavivirus from Aedes albopictus and Aedes flavopictus mosquitoes in Japan. Virology 391:119-29.
29. Blitvich BJ, Lin M, Dorman KS, Soto V, Hovav E, Tucker BJ, Staley M, Platt KB, Bartholomay LC. 2009. Genomic sequence and phylogenetic analysis of Culex flavivirus, an insect-specific flavivirus, isolated from Culex pipiens (Diptera: Culicidae) in Iowa. J Med Entomol 46:934-41.
30. Cook S, Moureau G, Harbach RE, Mukwaya L, Goodger K, Ssenfuka F, Gould E, Holmes EC, de Lamballerie X. 2009. Isolation of a novel species of flavivirus and a new strain of Culex flavivirus (Flaviviridae) from a natural mosquito population in Uganda. J Gen Virol 90:2669-2678.
31. Grard G, Moureau G, Charrel RN, Holmes EC, Gould EA, de Lamballerie X. 2010. Genomics and evolution of Aedes-borne flaviviruses. J Gen Virol 91:87-94.
32. Schubert AM, Putonti C. 2010. Evolution of the sequence composition of Flaviviruses. Infect Genet Evol 10:129-36.
33. Firth AE, Blitvich BJ, Wills NM, Miller CL, Atkins JF. 2010. Evidence for ribosomal frameshifting and a novel overlapping gene in the genomes of insect-specific flaviviruses. Virology 399:153-166.
34. Farfan-Ale JA, Loroño-Pino MA, Garcia-Rejon JE, Soto V, Lin M, Staley M, Dorman KS, Bartholomay LC, Hovav E, Blitvich BJ. 2010. Detection of flaviviruses and orthobunyaviruses in mosquitoes in the Yucatan Peninsula of Mexico in 2008. Vector Borne Zoonotic Dis 10:777-83.
35. Calzolari M, Bonilauri P, Bellini R, Caimi M, Defilippo F, Maioli G, Albieri A, Medici A, Veronesi R, Pilani R, Gelati A, Angelini P, Parco V, Fabbi M, Barbieri I, Lelli D, Lavazza A, Cordioli P, Dottori M. 2010. Arboviral survey of mosquitoes in two northern Italian regions in 2007 and 2008. Vector Borne Zoonotic Dis 10:875-84.
36. Saiyasombat R, Dorman KS, Garcia-Rejon JE, Loroño-Pino MA, Farfan-Ale JA, Blitvich BJ. 2010. Isolation and sequence analysis of Culex flavivirus from Culex interrogator and Culex quinquefasciatus in the Yucatan Peninsula of Mexico. Arch Virol 155:983-6.
37. Grant-Klein RJ, Baldwin CD, Turell MJ, Rossi CA, Li F, Lovari R, Crowder CD, Matthews HE, Rounds MA, Eshoo MW, Blyn LB, Ecker DJ, Sampath R, Whitehouse CA. 2010. Rapid identification of vector-borne flaviviruses by mass spectrometry. Mol Cell Probes 24:219-28.
38. Kent RJ, Crabtree MB, Miller BR. 2010. Transmission of West Nile virus by Culex quinquefasciatus say infected with Culex Flavivirus Izabal. PLoS Negl Trop Dis 4:e671.
39. Newman CM, Cerutti F, Anderson TK, Hamer GL, Walker ED, Kitron UD, Ruiz MO, Brawn JD, Goldberg TL. 2011. Culex flavivirus and West Nile virus mosquito coinfection and positive ecological association in Chicago, United States. Vector Borne Zoonotic Dis 11:1099-105.
40. Tyler S, Bolling BG, Blair CD, Brault AC, Pabbaraju K, Armijos MV, Clark DC, Calisher CH, Drebot MA. 2011. Distribution and phylogenetic comparisons of a novel mosquito flavivirus sequence present in Culex tarsalis Mosquitoes from western Canada with viruses isolated in California and Colorado. Am J Trop Med Hyg 85:162-8.
41. Bolling BG, Eisen L, Moore CG, Blair CD. 2011. Insect-specific flaviviruses from Culex mosquitoes in Colorado, with evidence of vertical transmission. Am J Trop Med Hyg 85:169-77.
42. Saiyasombat R, Bolling BG, Brault AC, Bartholomay LC, Blitvich BJ. 2011. Evidence of efficient transovarial transmission of Culex flavivirus by Culex pipiens (Diptera: Culicidae). J Med Entomol 48:1031-8.
43. Vázquez A, Sánchez-Seco MP, Palacios G, Molero F, Reyes N, Ruiz S, Aranda C, Marqués E, Escosa R, Moreno J, Figuerola J, Tenorio A. 2012. Novel flaviviruses detected in different species of mosquitoes in Spain. Vector Borne Zoonotic Dis 12:223-9.
44. Isawa H, Kuwata R, Tajima S, Hoshino K, Sasaki T, Takasaki T, Kobayashi M, Sawabe K. 2012. Construction of an infectious cDNA clone of Culex flavivirus, an insect-specific flavivirus from Culex mosquitoes. Arch Virol 157:975-9.
45. Crockett RK, Burkhalter K, Mead D, Kelly R, Brown J, Varnado W, Roy A, Horiuchi K, Biggerstaff BJ, Miller B, Nasci R. 2012. Culex flavivirus and West Nile virus in Culex quinquefasciatus populations in the southeastern United States. J Med Entomol 49:165-74.
46. Hoshino K, Takahashi-Nakaguchi A, Isawa H, Sasaki T, Higa Y, Kasai S, Tsuda Y, Sawabe K, Kobayashi M. 2012. Entomological surveillance for flaviviruses at migratory bird stopover sites in Hokkaido, Japan, and a new insect flavivirus detected in Aedes galloisi (Diptera: Culicidae). J Med Entomol 49:175-82.
47. Calzolari M, Zé-Zé L, Růžek D, Vázquez A, Jeffries C, Defilippo F, Osório HC, Kilian P, Ruíz S, Fooks AR, Maioli G, Amaro F, Tlustý M, Figuerola J, Medlock JM, Bonilauri P, Alves MJ, Šebesta O, Tenorio A, Vaux AGC, Bellini R, Gelbič I, Sánchez-Seco MP, Johnson N, Dottori M. 2012. Detection of mosquito-only flaviviruses in Europe. J Gen Virol 93:1215-1225.
48. Bolling BG, Olea-Popelka FJ, Eisen L, Moore CG, Blair CD. 2012. Transmission dynamics of an insect-specific flavivirus in a naturally infected Culex pipiens laboratory colony and effects of co-infection on vector

competence for West Nile virus. Virology 427:90-7.
49. Huanyu W, Haiyan W, Shihong F, Guifang L, Hong L, Xiaoyan G, Lizhi S, Rayner S, Aiqiang X, Guodong L. 2012. Isolation and identification of a distinct strain of Culex Flavivirus from mosquitoes collected in Mainland China. Virol J 9:73.
50. Parreira R, Cook S, Lopes Â, de Matos AP, de Almeida AP, Piedade J, Esteves A. 2012. Genetic characterization of an insect-specific flavivirus isolated from Culex theileri mosquitoes collected in southern Portugal. Virus Res 167:152-61.
51. Machado DC, Mondini A, dos Santos Santana V, Yonamine PT, Chiaravalloti Neto F, Zanotto PM, Nogueira ML. 2012. First identification of Culex flavivirus (Flaviviridae) in Brazil. Intervirology 55:475-83.
52. Huhtamo E, Moureau G, Cook S, Julkunen O, Putkuri N, Kurkela S, Uzcátegui NY, Harbach RE, Gould EA, Vapalahti O, de Lamballerie X. 2012. Novel insect-specific flavivirus isolated from northern Europe. Virology 433:471-8.
53. Roiz D, Vázquez A, Rosso F, Arnoldi D, Girardi M, Cuevas L, Perez-Pastrana E, Sánchez-Seco MP, Tenorio A, Rizzoli A. 2012. Detection of a new insect flavivirus and isolation of Aedes flavivirus in Northern Italy. Parasit Vectors 5:223.
54. Lee JS, Grubaugh ND, Kondig JP, Turell MJ, Kim HC, Klein TA, O'Guinn ML. 2013. Isolation and genomic characterization of Chaoyang virus strain ROK144 from Aedes vexans nipponii from the Republic of Korea. Virology 435:220-4.
55. An SY, Liu JS, Ren Y, Wang ZS, Han Y, Ding J, Guo JQ. 2012. [Isolation of the Culex flavivirus from mosquitoes in Liaoning Province, China]. Bing Du Xue Bao 28:511-6.
56. Grubaugh ND, Petz LN, Melanson VR, McMenamy SS, Turell MJ, Long LS, Pisarcik SE, Kengluecha A, Jaichapor B, O'Guinn ML, Lee JS. 2013. Evaluation of a field-portable DNA microarray platform and nucleic acid amplification strategies for the detection of arboviruses, arthropods, and bloodmeals. Am J Trop Med Hyg 88:245-53.
57. Hobson-Peters J, Yam AW, Lu JW, Setoh YX, May FJ, Kurucz N, Walsh S, Prow NA, Davis SS, Weir R, Melville L, Hunt N, Webb RI, Blitvich BJ, Whelan P, Hall RA. 2013. A new insect-specific flavivirus from northern Australia suppresses replication of West Nile virus and Murray Valley encephalitis virus in co-infected mosquito cells. PLoS One 8:e56534.
58. Chen YY, Lin JW, Fan YC, Tu WC, Chang GJ, Chiou SS. 2013. First detection of the Africa/Caribbean/Latin American subtype of Culex flavivirus in Asian country, Taiwan. Comp Immunol Microbiol Infect Dis 36:387-96.
59. Evangelista J, Cruz C, Guevara C, Astete H, Carey C, Kochel TJ, Morrison AC, Williams M, Halsey ES, Forshey BM. 2013. Characterization of a novel flavivirus isolated from Culex (Melanoconion) ocossa mosquitoes from Iquitos, Peru. J Gen Virol 94:1266-1272.
60. Obara-Nagoya M, Yamauchi T, Watanabe M, Hasegawa S, Iwai-Itamochi M, Horimoto E, Takizawa T, Takashima I, Kariwa H. 2013. Ecological and genetic analyses of the complete genomes of Culex flavivirus strains isolated from Culex tritaeniorhynchus and Culex pipiens (Diptera: Culicidae) group mosquitoes. J Med Entomol 50:300-9.
61. Haddow AD, Guzman H, Popov VL, Wood TG, Widen SG, Haddow AD, Tesh RB, Weaver SC. 2013. First isolation of Aedes flavivirus in the Western Hemisphere and evidence of vertical transmission in the mosquito Aedes (Stegomyia) albopictus (Diptera: Culicidae). Virology 440:134-9.
62. Ferreira DD, Cook S, Lopes Â, de Matos AP, Esteves A, Abecasis A, de Almeida AP, Piedade J, Parreira R. 2013. Characterization of an insect-specific flavivirus (OCFVPT) co-isolated from Ochlerotatus caspius collected in southern Portugal along with a putative new Negev-like virus. Virus Genes 47:532-45.
63. Kolodziejek J, Pachler K, Bin H, Mendelson E, Shulman L, Orshan L, Nowotny N. 2013. Barkedji virus, a novel mosquito-borne flavivirus identified in Culex perexiguus mosquitoes, Israel, 2011. J Gen Virol 94:2449-2457.
64. Grubaugh ND, McMenamy SS, Turell MJ, Lee JS. 2013. Multi-gene detection and identification of mosquito-borne RNA viruses using an oligonucleotide microarray. PLoS Negl Trop Dis 7:e2349.
65. Zuo S, Zhao Q, Guo X, Zhou H, Cao W, Zhang J. 2014. Detection of Quang Binh virus from mosquitoes in China. Virus Res 180:31-8.
66. Takhampunya R, Kim HC, Tippayachai B, Lee DK, Lee WJ, Chong ST, Kim MS, Lee JS, Klein TA. 2014. Distribution and mosquito hosts of Chaoyang virus, a newly reported flavivirus from the Republic of Korea, 2008-2011. J Med Entomol 51:464-74.
67. Papa A, Papadopoulou E, Kalaitzopoulou S, Tsioka K, Mourelatos S. 2014. Detection of West Nile virus and insect-specific flavivirus RNA in Culex mosquitoes, central Macedonia, Greece. Trans R Soc Trop Med Hyg 108:555-9.
68. L'Vov D K, Al'khovskiĭ SV, Shchelkanov M, Shchetinin AM, Deriabin PG, Gitel'man AK, Samokhvalov EI, Botikov AG. 2014. [Taxonomy of the Sokuluk virus (SOKV) (Flaviviridae, Flavivirus, Entebbe bat virus group) isolated from bats (Vespertilio pipistrellus Schreber, 1774), ticks (Argasidae Koch, 1844), and birds in Kyrgyzstan]. Vopr Virusol 59:30-4.
69. Huhtamo E, Cook S, Moureau G, Uzcátegui NY, Sironen T, Kuivanen S, Putkuri N, Kurkela S, Harbach RE, Firth AE, Vapalahti O, Gould EA, de Lamballerie X. 2014. Novel flaviviruses from mosquitoes: mosquito-specific evolutionary lineages within the phylogenetic group of mosquito-borne flaviviruses. Virology 464-465:320-329.
70. Goenaga S, Fabbri CM, García JB, Rondán JC, Gardenal N, Calderón GE, Enria DA, Levis SM. 2014. New strains

of Culex flavivirus isolated in Argentina. J Med Entomol 51:900-6.
71. Kenney JL, Solberg OD, Langevin SA, Brault AC. 2014. Characterization of a novel insect-specific flavivirus from Brazil: potential for inhibition of infection of arthropod cells with medically important flaviviruses. J Gen Virol 95:2796-2808.
72. Saiyasombat R, Carrillo-Tripp J, Miller WA, Bredenbeek PJ, Blitvich BJ. 2014. Substitution of the premembrane and envelope protein genes of Modoc virus with the homologous sequences of West Nile virus generates a chimeric virus that replicates in vertebrate but not mosquito cells. Virol J 11:150.
73. Rizzo F, Cerutti F, Ballardini M, Mosca A, Vitale N, Radaelli MC, Desiato R, Prearo M, Pautasso A, Casalone C, Acutis P, Peletto S, Mandola ML. 2014. Molecular characterization of flaviviruses from field-collected mosquitoes in northwestern Italy, 2011-2012. Parasit Vectors 7:395.
74. Pauvolid-Corrêa A, Solberg O, Couto-Lima D, Kenney J, Serra-Freire N, Brault A, Nogueira R, Langevin S, Komar N. 2015. Nhumirim virus, a novel flavivirus isolated from mosquitoes from the Pantanal, Brazil. Arch Virol 160:21-7.
75. Grisenti M, Vázquez A, Herrero L, Cuevas L, Perez-Pastrana E, Arnoldi D, Rosà R, Capelli G, Tenorio A, Sánchez-Seco MP, Rizzoli A. 2015. Wide detection of Aedes flavivirus in north-eastern Italy--a European hotspot of emerging mosquito-borne diseases. J Gen Virol 96:420-430.
76. Blitvich BJ, Firth AE. 2015. Insect-specific flaviviruses: a systematic review of their discovery, host range, mode of transmission, superinfection exclusion potential and genomic organization. Viruses 7:1927-59.
77. Zheng Y, Cao Y, Fu S, Cheng J, Zhao J, Dai P, Kong X, Liang G. 2015. [Isolation and identification of mosquito-borne arboviruses in Yuncheng city, Shanxi province, 2012]. Zhonghua Liu Xing Bing Xue Za Zhi 36:368-73.
78. Datta S, Gopalakrishnan R, Chatterjee S, Veer V. 2015. Phylogenetic Characterization of a Novel Insect-Specific Flavivirus Detected in a Culex Pool, Collected from Assam, India. Intervirology 58:149-54.
79. Kuwata R, Sugiyama H, Yonemitsu K, Van Dung N, Terada Y, Taniguchi M, Shimoda H, Takano A, Maeda K. 2015. Isolation of Japanese encephalitis virus and a novel insect-specific flavivirus from mosquitoes collected in a cowshed in Japan. Arch Virol 160:2151-9.
80. Liang W, He X, Liu G, Zhang S, Fu S, Wang M, Chen W, He Y, Tao X, Jiang H, Lin X, Gao X, Hu W, Liu Y, Feng L, Cao Y, Yang G, Jing C, Liang G, Wang H. 2015. Distribution and phylogenetic analysis of Culex flavivirus in mosquitoes in China. Arch Virol 160:2259-68.
81. Calzolari M, Zé-Zé L, Vázquez A, Sánchez Seco MP, Amaro F, Dottori M. 2016. Insect-specific flaviviruses, a worldwide widespread group of viruses only detected in insects. Infect Genet Evol 40:381-388.
82. Turell MJ. 2015. Experimental Transmission of Karshi (Mammalian Tick-Borne Flavivirus Group) Virus by Ornithodoros Ticks >2,900 Days after Initial Virus Exposure Supports the Role of Soft Ticks as a Long-Term Maintenance Mechanism for Certain Flaviviruses. PLoS Negl Trop Dis 9:e0004012.
83. Carrera JP, Guzman H, Beltrán D, Díaz Y, López-Vergès S, Torres-Cosme R, Popov V, Widen SG, Wood TG, Weaver SC, Cáceres-Carrera L, Vasilakis N, Tesh RB. 2015. Mercadeo Virus: A Novel Mosquito-Specific Flavivirus from Panama. Am J Trop Med Hyg 93:1014-9.
84. Kuwata R, Isawa H, Hoshino K, Sasaki T, Kobayashi M, Maeda K, Sawabe K. 2015. Analysis of Mosquito-Borne Flavivirus Superinfection in Culex tritaeniorhynchus (Diptera: Culicidae) Cells Persistently Infected with Culex Flavivirus (Flaviviridae). J Med Entomol 52:222-9.
85. Alkan C, Zapata S, Bichaud L, Moureau G, Lemey P, Firth AE, Gritsun TS, Gould EA, de Lamballerie X, Depaquit J, Charrel RN. 2015. Ecuador Paraiso Escondido Virus, a New Flavivirus Isolated from New World Sand Flies in Ecuador, Is the First Representative of a Novel Clade in the Genus Flavivirus. J Virol 89:11773-85.
86. Newman CM, Anderson TK, Goldberg TL. 2016. Decreased Flight Activity in Culex pipiens (Diptera: Culicidae) Naturally Infected With Culex flavivirus. J Med Entomol 53:233-6.
87. Goenaga S, Kenney JL, Duggal NK, Delorey M, Ebel GD, Zhang B, Levis SC, Enria DA, Brault AC. 2015. Potential for Co-Infection of a Mosquito-Specific Flavivirus, Nhumirim Virus, to Block West Nile Virus Transmission in Mosquitoes. Viruses 7:5801-12.
88. Cao YX, He XX, Fu SH, He Y, Li H, Gao XY, Liang GD, Wang HY. 2015. Real-time RT-PCR Assay for the Detection of Culex flavivirus. Biomed Environ Sci 28:917-9.
89. Misencik MJ, Grubaugh ND, Andreadis TG, Ebel GD, Armstrong PM. 2016. Isolation of a Novel Insect-Specific Flavivirus from Culiseta melanura in the Northeastern United States. Vector Borne Zoonotic Dis 16:181-90.
90. Fernandes LN, de Paula MB, Araújo AB, Gonçalves EF, Romano CM, Natal D, Malafronte Rdos S, Marrelli MT, Levi JE. 2016. Detection of Culex flavivirus and Aedes flavivirus nucleotide sequences in mosquitoes from parks in the city of São Paulo, Brazil. Acta Trop 157:73-83.
91. Zhu Z, Chan JF, Tee KM, Choi GK, Lau SK, Woo PC, Tse H, Yuen KY. 2016. Comparative genomic analysis of pre-epidemic and epidemic Zika virus strains for virological factors potentially associated with the rapidly expanding epidemic. Emerg Microbes Infect 5:e22.
92. Fan H, Zhao Q, Guo X, Sun Q, Zuo S, Wu C, Zhou H, An X, Pei G, Tong Y, Zhang J, Shi T. 2016. Complete genome sequence of Xishuangbanna flavivirus, a novel mosquito-specific flavivirus from China. Arch Virol 161:1723-7.
93. Papa A, Papadopoulou E, Paliwal R, Kalaitzopoulou S, Mourelatos S, Niedrig M. 2016. Insect-specific flaviviruses in Aedes mosquitoes in Greece. Arch Virol 161:2183-8.
94. Grant A, Ponia SS, Tripathi S, Balasubramaniam V, Miorin L, Sourisseau M, Schwarz MC, Sánchez-Seco MP, Evans MJ, Best SM, García-Sastre A. 2016. Zika Virus Targets Human STAT2 to Inhibit Type I Interferon Signaling. Cell Host Microbe 19:882-90.

119. Nouri S, Matsumura EE, Kuo YW, Falk BW. 2018. Insect-specific viruses: from discovery to potential translational applications. Curr Opin Virol 33:33-41.
120. White SK, Lednicky JA, Okech BA, Morris JG, Jr., Dunford JC. 2018. Spondweni Virus in Field-Caught Culex quinquefasciatus Mosquitoes, Haiti, 2016. Emerg Infect Dis 24:1765-1767.
121. Charles J, Tangudu CS, Hurt SL, Tumescheit C, Firth AE, Garcia-Rejon JE, Machain-Williams C, Blitvich BJ. 2018. Detection of novel and recognized RNA viruses in mosquitoes from the Yucatan Peninsula of Mexico using metagenomics and characterization of their in vitro host ranges. J Gen Virol 99:1729-1738.
122. Romo H, Kenney JL, Blitvich BJ, Brault AC. 2018. Restriction of Zika virus infection and transmission in Aedes aegypti mediated by an insect-specific flavivirus. Emerg Microbes Infect 7:181.
123. Ajamma YU, Onchuru TO, Ouso DO, Omondi D, Masiga DK, Villinger J. 2018. Vertical transmission of naturally occurring Bunyamwera and insect-specific flavivirus infections in mosquitoes from islands and mainland shores of Lakes Victoria and Baringo in Kenya. PLoS Negl Trop Dis 12:e0006949.
124. Fernandes LN, Coletti TM, Monteiro FJC, Rego M, Ribeiro ESD, Ribeiro GO, Marinho R, Komninakis SV, Witkin SS, Deng X, Delwart E, Sabino EC, Leal É, Costa ACD. 2018. A Novel Highly Divergent Strain of Cell Fusing Agent Virus (CFAV) in Mosquitoes from the Brazilian Amazon Region. Viruses 10.
125. Ndiaye EH, Diallo D, Fall G, Ba Y, Faye O, Dia I, Diallo M. 2018. Arboviruses isolated from the Barkedji mosquito-based surveillance system, 2012-2013. BMC Infect Dis 18:642.
126. Miranda J, Mattar S, Gonzalez M, Hoyos-López R, Aleman A, Aponte J. 2019. First report of Culex flavivirus infection from Culex coronator (Diptera: Culicidae), Colombia. Virol J 16:1.
127. Tangudu CS, Charles J, Hurt SL, Dunphy BM, Smith RC, Bartholomay LC, Blitvich BJ. 2019. Skunk River virus, a novel orbivirus isolated from Aedes trivittatus in the United States. J Gen Virol 100:295-300.
128. Hannon ER, Jackson KC, Biggerstaff BJ, Raman V, Komar N. 2019. Bloodmeal Host Selection of Culex quinquefasciatus (Diptera: Culicidae) in Las Vegas, Nevada, United States. J Med Entomol 56:603-608.
129. Salazar V, Jagger BW, Mongkolsapaya J, Burgomaster KE, Dejnirattisai W, Winkler ES, Fernandez E, Nelson CA, Fremont DH, Pierson TC, Crowe JE, Jr., Screaton GR, Diamond MS. 2019. Dengue and Zika Virus Cross-Reactive Human Monoclonal Antibodies Protect against Spondweni Virus Infection and Pathogenesis in Mice. Cell Rep 26:1585-1597.e4.
130. Rademan R, Markotter W, Paweska JT, Jansen van Vuren P. 2019. Multiplex real-time RT-PCR for detection and distinction of Spondweni and Zika virus. J Virol Methods 266:72-76.
131. García-Nicolás O, Lewandowska M, Ricklin ME, Summerfield A. 2019. Monocyte-Derived Dendritic Cells as Model to Evaluate Species Tropism of Mosquito-Borne Flaviviruses. Front Cell Infect Microbiol 9:5.
132. Moraes OS, Cardoso BF, Pacheco TA, Pinto AZL, Carvalho MS, Hahn RC, Burlamaqui TCT, Oliveira LF, Oliveira RS, Vasconcelos JM, Lemos PS, Nunes MRT, Slhessarenko RD. 2019. Natural infection by Culex flavivirus in Culex quinquefasciatus mosquitoes captured in Cuiabá, Mato Grosso Mid-Western Brazil. Med Vet Entomol 33:397-406.
133. Akıner MM, Öztürk M, Başer AB, Günay F, Hacıoğlu S, Brinkmann A, Emanet N, Alten B, Özkul A, Nitsche A, Linton YM, Ergünay K. 2019. Arboviral screening of invasive Aedes species in northeastern Turkey: West Nile virus circulation and detection of insect-only viruses. PLoS Negl Trop Dis 13:e0007334.
134. Martin E, Borucki MK, Thissen J, Garcia-Luna S, Hwang M, Wise de Valdez M, Jaing CJ, Hamer GL, Frank M. 2019. Mosquito-Borne Viruses and Insect-Specific Viruses Revealed in Field-Collected Mosquitoes by a Monitoring Tool Adapted from a Microbial Detection Array. Appl Environ Microbiol 85.
135. Piyasena TBH, Newton ND, Hobson-Peters J, Vet LJ, Setoh YX, Bielefeldt-Ohmann H, Khromykh AA, Hall RA. 2019. Chimeric viruses of the insect-specific flavivirus Palm Creek with structural proteins of vertebrate-infecting flaviviruses identify barriers to replication of insect-specific flaviviruses in vertebrate cells. J Gen Virol 100:1580-1586.
136. Birnberg L, Talavera S, Aranda C, Núñez AI, Napp S, Busquets N. 2019. Field-captured Aedes vexans (Meigen, 1830) is a competent vector for Rift Valley fever phlebovirus in Europe. Parasit Vectors 12:484.
137. Jegal S, Jun H, Kim-Jeon MD, Park SH, Ahn SK, Lee J, Gong YW, Joo K, Kwon MJ, Roh JY, Lee WG, Lee W, Bahk YY, Kim TS. 2020. Three-year surveillance of culicine mosquitoes (Diptera: Culicidae) for flavivirus infections in Incheon Metropolitan City and Hwaseong-si of Gyeonggi-do Province, Republic of Korea. Acta Trop 202:105258.
138. Goenaga S, Goenaga J, Boaglio ER, Enria DA, Levis SDC. 2020. Superinfection exclusion studies using West Nile virus and Culex flavivirus strains from Argentina. Mem Inst Oswaldo Cruz 115:e200012.
139. Jaeger AS, Weiler AM, Moriarty RV, Rybarczyk S, O'Connor SL, O'Connor DH, Seelig DM, Fritsch MK, Friedrich TC, Aliota MT. 2020. Spondweni virus causes fetal harm in Ifnar1(-/-) mice and is transmitted by Aedes aegypti mosquitoes. Virology 547:35-46.
140. Yap G, Mailepessov D, Lim XF, Chan S, How CB, Humaidi M, Yeo G, Chong CS, Lam-Phua SG, Lee R, Okumura C, Vythilingam I, Ng LC. 2020. Detection of Japanese Encephalitis Virus in Culex Mosquitoes in Singapore. Am J Trop Med Hyg 103:1234-1240.
141. Kubacki J, Flacio E, Qi W, Guidi V, Tonolla M, Fraefel C. 2020. Viral Metagenomic Analysis of Aedes albopictus Mosquitos from Southern Switzerland. Viruses 12.
142. Wastika CE, Harima H, Sasaki M, Hang'ombe BM, Eshita Y, Qiu Y, Hall WW, Wolfinger MT, Sawa H, Orba Y. 2020. Discoveries of Exoribonuclease-Resistant Structures of Insect-Specific Flaviviruses Isolated in Zambia. Viruses 12.
143. Atoni E, Zhao L, Hu C, Ren N, Wang X, Liang M, Mwaliko C, Yuan Z, Xia H. 2020. A dataset of distribution and diversity of mosquito-associated viruses and their mosquito vectors in China. Sci Data 7:342.

144. Bahk YY, Park SH, Kim-Jeon MD, Oh SS, Jung H, Jun H, Kim KA, Park JM, Ahn SK, Lee J, Choi EJ, Moon BS, Gong YW, Kwon MJ, Kim TS. 2020. Monitoring Culicine Mosquitoes (Diptera: Culicidae) as a Vector of Flavivirus in Incheon Metropolitan City and Hwaseong-Si, Gyeonggi-Do, Korea, during 2019. Korean J Parasitol 58:551-558.
145. McLean BJ, Hall-Mendelin S, Webb CE, Bielefeldt-Ohmann H, Ritchie SA, Hobson-Peters J, Hall RA, van den Hurk AF. 2021. The Insect-Specific Parramatta River Virus Is Vertically Transmitted by Aedes vigilax Mosquitoes and Suppresses Replication of Pathogenic Flaviviruses In Vitro. Vector Borne Zoonotic Dis 21:208-215.
146. Renner M, Dejnirattisai W, Carrique L, Martin IS, Karia D, Ilca SL, Ho SF, Kotecha A, Keown JR, Mongkolsapaya J, Screaton GR, Grimes JM. 2021. Flavivirus maturation leads to the formation of an occupied lipid pocket in the surface glycoproteins. Nat Commun 12:1238.
147. Klein A, Strube C, Becker SC, Naccache F. 2021. Screening for Viruses and Lemur-Associated Filara in Wild-Caught Mosquitoes From Madagascar. J Med Entomol 58:983-989.
148. Fang Y, Tambo E, Xue JB, Zhang Y, Zhou XN, Khater EIM. 2021. Detection of DENV-2 and Insect-Specific Flaviviruses in Mosquitoes Collected From Jeddah, Saudi Arabia. Front Cell Infect Microbiol 11:626368.
149. Tangudu CS, Charles J, Nunez-Avellaneda D, Hargett AM, Brault AC, Blitvich BJ. 2021. Chimeric Zika viruses containing structural protein genes of insect-specific flaviviruses cannot replicate in vertebrate cells due to entry and post-translational restrictions. Virology 559:30-39.
150. Fang Y, Li XS, Zhang W, Xue JB, Wang JZ, Yin SQ, Li SG, Li XH, Zhang Y. 2021. Molecular epidemiology of mosquito-borne viruses at the China-Myanmar border: discovery of a potential epidemic focus of Japanese encephalitis. Infect Dis Poverty 10:57.
151. Akiyama BM, Graham ME, Z OD, Beckham JD, Kieft JS. 2021. Three-dimensional structure of a flavivirus dumbbell RNA reveals molecular details of an RNA regulator of replication. Nucleic Acids Res 49:7122-7138.
152. Chatterjee S, Kim CM, Yun NR, Kim DM, Song HJ, Chung KA. 2021. Molecular detection and identification of Culex flavivirus in mosquito species from Jeju, Republic of Korea. Virol J 18:150.
153. Guarido MM, Govender K, Riddin MA, Schrama M, Gorsich EE, Brooke BD, Almeida APG, Venter M. 2021. Detection of Insect-Specific Flaviviruses in Mosquitoes (Diptera: Culicidae) in Northeastern Regions of South Africa. Viruses 13.
154. Öhlund P, Hayer J, Hesson JC, Blomström AL. 2021. Small RNA Response to Infection of the Insect-Specific Lammi Virus and Hanko Virus in an Aedes albopictus Cell Line. Viruses 13.
155. Öhlund P, Delhomme N, Hayer J, Hesson JC, Blomström AL. 2022. Transcriptome Analysis of an Aedes albopictus Cell Line Single- and Dual-Infected with Lammi Virus and WNV. Int J Mol Sci 23.
156. Peinado SA, Aliota MT, Blitvich BJ, Bartholomay LC. 2022. Biology and Transmission Dynamics of Aedes flavivirus. J Med Entomol 59:659-666.
157. Zhang Y, Liang D, Yuan F, Yan Y, Wang Z, Liu P, Yu Q, Zhang X, Wang X, Zheng A. 2022. Replication is the key barrier during the dual-host adaptation of mosquito-borne flaviviruses. Proc Natl Acad Sci U S A 119:e2110491119.
158. Habarugira G, Moran J, Harrison JJ, Isberg SR, Hobson-Peters J, Hall RA, Bielefeldt-Ohmann H. 2022. Evidence of Infection with Zoonotic Mosquito-Borne Flaviviruses in Saltwater Crocodiles (Crocodylus porosus) in Northern Australia. Viruses 14.
159. Bai Y, Zhang Y, Su Z, Tang S, Wang J, Wu Q, Yang J, Moming A, Zhang Y, Bell-Sakyi L, Sun S, Shen S, Deng F. 2022. Discovery of Tick-Borne Karshi Virus Implies Misinterpretation of the Tick-Borne Encephalitis Virus Seroprevalence in Northwest China. Front Microbiol 13:872067.
160. Huang Y, Zhang H, Li X, Zhao L, Cai D, Wang S, Ren N, Ma H, Huang D, Wang F, Yuan Z, Zhang B, Xia H. 2022. In Vitro and In Vivo Characterization of a New Strain of Mosquito Flavivirus Derived from Culicoides. Viruses 14.
161. Tangudu CS, Hargett AM, Blitvich BJ. 2022. Evidence that untranslated genomic sequences are key determinants of insect-specific flavivirus host restriction. Virology 574:102-114.
162. Kullappan M, Benedict BA, Rajajagadeesan A, Baskaran P, Periadurai ND, Ambrose JM, Gandhamaneni SH, Nakkella AK, Agarwal A, Veeraraghavan VP, Surapaneni KM. 2022. Ellagic Acid as a Potential Inhibitor against the Nonstructural Protein NS3 Helicase of Zika Virus: A Molecular Modelling Study. Biomed Res Int 2022:2044577.
163. Colmant AMG, Warrilow D, Hall-Mendelin S, Onn M, Hobson-Peters J, Huang B, Kurucz N, Warchot A, Primmer BR, Isberg S, Bielefeldt-Ohmann H, Hall RA. 2022. Arthropod-Borne Virus Surveillance as a Tool to Study the Australian Mosquito Virome. Viruses 14.
164. Coletti TM, Romano CM, Urbinatti PR, Teixeira RS, Pedrosa LWA, Nardi MS, Natal D, Costa ACD, Jesus JG, Claro IM, Sabino EC, Witkin SS, Marrelli MT, Fernandes LN. 2022. Characterization of insect-specific Culex flavivirus (Flaviviridae) nucleotide sequences in mosquitoes from urban parks in São Paulo, Brazil. Rev Soc Bras Med Trop 55:e0067.
165. Wen D, Ding LS, Zhang Y, Li X, Zhang X, Yuan F, Zhao T, Zheng A. 2022. Suppression of flavivirus transmission from animal hosts to mosquitoes with a mosquito-delivered vaccine. Nat Commun 13:7780.
166. Walsh E, Torres TZB, Rückert C. 2022. Culex Mosquito Piwi4 Is Antiviral against Two Negative-Sense RNA Viruses. Viruses 14.
167. Fang Y, Hang T, Yang LM, Xue JB, Fujita R, Feng XS, Jiang TG, Zhang Y, Li SZ, Zhou XN. 2023. Long-distance spread of Tembusu virus, and its dispersal in local mosquitoes and domestic poultry in Chongming Island, China. Infect Dis Poverty 12:52.

168. Jaeger AS, Crooks CM, Weiler AM, Bliss MI, Rybarczyk S, Richardson A, Einwalter M, Peterson E, Capuano S, 3rd, Barkhymer A, Becker JT, Greene JT, Freedman TS, Langlois RA, Friedrich TC, Aliota MT. 2023. Primary infection with Zika virus provides one-way heterologous protection against Spondweni virus infection in rhesus macaques. Sci Adv 9:eadg3444.
169. Anakha A, Shah HK, Fathima PA, Aiswarya RS, Ajithlal PM, Kumar NP, Kumar A, Saini P. 2023. Identification and molecular characterization of Culex flavivirus in Culex quinquefasciatus (Diptera: Culicidae): first report from India. Trans R Soc Trop Med Hyg 117:839-843.
170. Tangudu CS, Hargett AM, Mitrisin BC, Laredo-Tiscareño SV, Blitvich BJ. 2023. Production of a chimeric flavivirus that contains the major structural glycoprotein genes of T'Ho virus in the genetic background of Zika virus. Virol J 20:197.
171. Tang X, Li R, Qi Y, Li W, Liu Z, Wu J. 2023. The identification and genetic characteristics of Quang Binh virus from field-captured Culex tritaeniorhynchus (Diptera: Culicidae) from Guizhou Province, China. Parasit Vectors 16:318.
172. Lwande OW, Näslund J, Sjödin A, Lantto R, Luande VN, Bucht G, Ahlm C, Agwanda B, Obanda V, Evander M. 2024. Novel strains of Culex flavivirus and Hubei chryso-like virus 1 from the Anopheles mosquito in western Kenya. Virus Res 339:199266.
173. Yuan H, Rao J, Zhang J, Ye J, Cao S, Chen H, Song Y. 2024. Japanese encephalitis virus inhibits superinfection of Zika virus in cells by the NS2B protein. J Virol 98:e0185923.
174. Kholodilov IS, Aibulatov SV, Khalin AV, Polienko AE, Klimentov AS, Belova OA, Rogova AA, Medvedev SG, Karganova GG. 2024. Orthoflavivirus Lammi in Russia: Possible Transovarial Transmission and Trans-Stadial Survival in Aedes cinereus (Diptera, Culicidae). Viruses 16.
175. Wang LL, Cheng Q, Newton ND, Wolfinger MT, Morgan MS, Slonchak A, Khromykh AA, Cheng TY, Parry RH. 2024. Xinyang flavivirus, from Haemaphysalis flava ticks in Henan Province, China, defines a basal, likely tick-only Orthoflavivirus clade. J Gen Virol 105.
176. Besson B, Overheul GJ, Wolfinger MT, Junglen S, van Rij RP. 2024. Pan-flavivirus analysis reveals sfRNA-independent, 3' UTR-biased siRNA production from an insect-specific flavivirus. J Virol 98:e0121524.
177. Nelson AN, Ploss A. 2024. Emerging mosquito-borne flaviviruses. mBio 15:e0294624.
178. Dingle AO, Adelman ZN. 2024. Evaluating the specificity of flavivirus proteases in Aedes aegypti cells for dengue virus 2-derived cleavage sites. PLoS One 19:e0309095.
179. Fu Y, Zhao W, Wu S, Li J, Liu Q, Jiang F, Lu H, Kang L, Xia Q, Cui F. 2025. Risk assessment of an Aedes flavivirus and its effect on pathogenic flavivirus replication in mosquitoes. Parasit Vectors 18:88.
180. Khasa R, Ogden SC, Wang Y, Mou Z, Metzler AD, Xie X, Dai X, Tang H. 2025. A single mutation in the PrM gene of Zika virus determines AXL dependency for infection of human neural cells. J Virol 99:e0187324.
181. Sewade W, Polat C, Kasap OE. 2025. Molecular evidence of Wolbachia and Orthoflavivirus infection in field-collected mosquitoes in three provinces of Türkiye. Trop Med Int Health 30:498-508.
182. Aranda-Coello JM, Machain-Williams C, Weber M, Dzul Rosado AR, Simpkins TR, Blitvich BJ. 2025. Serologic Surveillance for Orthoflaviviruses and Chikungunya Virus in Bats and Opossums in Chiapas, Mexico. Viruses 17.
183. Miao Q, Deng L, Le X, Li Q, Ning Y, Duan Y, Liu Q, Tao Y, Wang B, Xia X. 2025. Genetic and Antiviral Potential Characterization of Four Insect-Specific Viruses Identified and Isolated from Mosquitoes in Yunnan Province. Viruses 17.
184. Ang CCW, Ang ATY, Yam J, Lee RX, Lim MJ, Loh ZY, Torno M, Hakim L, Wong JCC, Harrison JJ, Hobson-Peters J, Tan CH, Setoh YX. 2025. Spatiotemporal prevalence and characterization of the lineage I insect-specific flavivirus, Quang Binh virus, isolated from Culex gelidus mosquitoes in Singapore. J Gen Virol 106.
185. Tangudu CS, Dankaona W, Blitvich BJ. 2025. Exchanging the stem-loop A promoter of Zika virus with the corresponding region of tick-borne encephalitis virus generates a chimeric virus that replicates in vertebrate and mosquito cells. Arch Virol 170:236.
186. Ran X, Xiao D, Wu Y, Shi Y, Liu S, Bai Y, Zhang Q, Liu L, Liu Q, Li J, Zhao M. 2026. Differences in metavirome among Aedes albopictus, Culex tritaeniorhynchus, and Anopheles sinensis in Jiangxi Province, China. Parasit Vectors 19:14.
187. Chan JJX, Zhao Z, Vieira C, Goh JZH, Slonchak A. 2026. Current Insights into Superinfection Exclusion in Insect-Specific Orthoflaviviruses. Viruses 18.
188. Kobayashi H, Tabata K, Sasaki M, Hall WW, Sawa H, Orba Y. 2026. NS4B and NS5 mutations expand the temperature range permissive for the replication of dual-host affiliated insect-specific flaviviruses in mosquito cells. Arch Virol 171.
189. Haldar I, Arif W, Devaraju P, Sihag KK, Srirama S, Balakrishnan V, Srinivasan P, Ramasamy A, Rahi M. 2026. First report of Culex flavivirus and its association with Wolbachia in Culex quinquefasciatus from Puducherry, India. Trans R Soc Trop Med Hyg 120:851-861.
190. Öhlund P, Pinto MC, Blomström AL. 2026. Transcriptomic and RNAi-mediated antiviral response to Kamiti River virus infection in Aedes albopictus U4.4 cells. Sci Rep 16.

**Table S4**. Within-strain amino acid p-distances of named orthoflaviviruses (n = 44)

| **Virus** | **Species** | **Count** | ***PP* AA distance[1]** | **NS5 AA distance[1]** | ***E* AA distance[1]** |
|---|---|---|---|---|---|
| Aroa virus | *Orthoflavivirus aroaense* | 1 | 0.003 (0.003-0.003) | 0.000 (0.000-0.000) | 0.000 (0.000-0.000) |
| Dakar_bat virus | *Orthoflavivirus dakarense* | 1 | 0.024 (0.024-0.024) | 0.000 (0.000-0.000) | 0.000 (0.000-0.000) |
| Jutiapa virus | *Orthoflavivirus jutiapaense* | 1 | 0.011 (0.011-0.011) | 0.000 (0.000-0.000) | 0.000 (0.000-0.000) |
| Ntaya virus | *Orthoflavivirus ntayaense* | 1 | 0.020 (0.020-0.020) | 0.000 (0.000-0.000) | 0.000 (0.000-0.000) |
| Deer_tick virus | *Orthoflavivirus powassanense* | 1 | 0.002 (0.002-0.002) | 0.000 (0.000-0.000) | 0.001 (0.001-0.001) |
| Banzi virus | *Orthoflavivirus banziense* | 1 | 0.020 (0.020-0.020) | 0.001 (0.001-0.001) | 0.000 (0.000-0.000) |
| Phnom_Penh_bat virus | *Orthoflavivirus phnompenhense* | 1 | 0.055 (0.055-0.055) | 0.001 (0.001-0.001) | 0.001 (0.001-0.001) |
| Kedougou virus | *Orthoflavivirus kedougouense* | 1 | 0.014 (0.014-0.014) | 0.002 (0.002-0.002) | 0.002 (0.002-0.002) |
| Ilheus virus | *Orthoflavivirus ilheusense* | 153 | 0.007 (0.001-0.019) | 0.002 (0.009-0.009) | 0.030 (0.000-0.056) |
| Stratford virus | *Orthoflavivirus kokoberaorum* | 10 | 0.007 (0.000-0.012) | 0.004 (0.007-0.007) | 0.014 (0.000-0.024) |
| Sokuluk virus | *Orthoflavivirus entebbeense* | 1 | 0.016 (0.016-0.016) | 0.004 (0.004-0.004) | 0.003 (0.003-0.003) |
| Usutu virus | *Orthoflavivirus usutuense* | 190 | 0.007 (0.000-0.018) | 0.005 (0.014-0.014) | 0.018 (0.001-0.041) |
| Langat virus | *Orthoflavivirus langatense* | 15 | 0.012 (0.000-0.025) | 0.005 (0.014-0.014) | 0.015 (0.000-0.046) |
| Kyasanur Forest disease | *Orthoflavivirus kyasanurense* | 91 | 0.012 (0.001-0.025) | 0.005 (0.010-0.010) | 0.021 (0.000-0.034) |
| Wesselsbron virus | *Orthoflavivirus wesselsbronense* | 190 | 0.009 (0.000-0.019) | 0.005 (0.013-0.013) | 0.038 (0.000-0.068) |
| Rocio virus | *Orthoflavivirus ilheusense* | 1 | 0.009 (0.009-0.009) | 0.006 (0.006-0.006) | 0.003 (0.003-0.003) |
| Alkhumra virus | *Orthoflavivirus kyasanurense* | 15 | 0.011 (0.001-0.017) | 0.006 (0.009-0.009) | 0.007 (0.000-0.009) |
| ITMV | *Orthoflavivirus israelense* | 15 | 0.009 (0.000-0.014) | 0.008 (0.013-0.013) | 0.033 (0.000-0.055) |
| Rio_Bravo virus | *Orthoflavivirus bravoense* | 1 | 0.026 (0.026-0.026) | 0.009 (0.009-0.009) | 0.008 (0.008-0.008) |
| Iguape virus | *Orthoflavivirus aroaense* | 1 | 0.013 (0.013-0.013) | 0.010 (0.010-0.010) | 0.007 (0.007-0.007) |
| Koutango virus | *Orthoflavivirus koutangoense* | 6 | 0.013 (0.005-0.017) | 0.011 (0.016-0.016) | 0.080 (0.016-0.112) |
| Powassan virus | *Orthoflavivirus powassanense* | 153 | 0.018 (0.002-0.055) | 0.011 (0.038-0.038) | 0.038 (0.002-0.139) |
| St Louis encephalitis virus | *Orthoflavivirus louisense* | 190 | 0.010 (0.000-0.019) | 0.011 (0.025-0.025) | 0.038 (0.000-0.080) |
| Tembusu virus | *Orthoflavivirus tembusu* | 190 | 0.022 (0.002-0.043) | 0.012 (0.024-0.024) | 0.066 (0.000-0.130) |

| | | | | | |
|---|---|---|---|---|---|
| Bagaza virus | *Orthoflavivirus bagazaense* | 171 | 0.018 (0.000-0.038) | 0.013 (0.046-0.046) | 0.044 (0.000-0.079) |
| Louping ill virus | *Orthoflavivirus loupingi* | 190 | 0.017 (0.002-0.038) | 0.013 (0.047-0.047) | 0.032 (0.000-0.083) |
| MVEV encephalitis | *Orthoflavivirus murrayense* | 171 | 0.019 (0.000-0.042) | 0.014 (0.031-0.031) | 0.057 (0.000-0.124) |
| OHFV | *Orthoflavivirus omskense* | 190 | 0.019 (0.001-0.047) | 0.015 (0.035-0.035) | 0.044 (0.000-0.110) |
| Dengue virus type 3 | *Orthoflavivirus denguei* | 190 | 0.015 (0.002-0.026) | 0.015 (0.031-0.031) | 0.046 (0.001-0.087) |
| Entebbe bat virus | *Orthoflavivirus entebbeense* | 3 | 0.030 (0.016-0.038) | 0.016 (0.023-0.023) | 0.035 (0.003-0.051) |
| Dengue virus type 1 | *Orthoflavivirus denguei* | 190 | 0.022 (0.003-0.037) | 0.016 (0.031-0.031) | 0.060 (0.001-0.092) |
| West Nile virus | *Orthoflavivirus nilense* | 190 | 0.019 (0.000-0.064) | 0.017 (0.061-0.061) | 0.064 (0.001-0.213) |
| Dengue virus type 4 | *Orthoflavivirus denguei* | 190 | 0.020 (0.000-0.043) | 0.020 (0.053-0.053) | 0.055 (0.002-0.095) |
| Kokobera virus | *Orthoflavivirus kokoberaorum* | 1 | 0.019 (0.019-0.019) | 0.020 (0.020-0.020) | 0.046 (0.046-0.046) |
| Yellow fever virus | *Orthoflavivirus flavi* | 190 | 0.022 (0.000-0.073) | 0.021 (0.067-0.067) | 0.059 (0.000-0.161) |
| Japanese encephalitis viru | *Orthoflavivirus japonicum* | 253 | 0.025 (0.000-0.057) | 0.022 (0.061-0.061) | 0.091 (0.000-0.162) |
| Zika virus | *Orthoflavivirus zikaense* | 190 | 0.019 (0.001-0.037) | 0.022 (0.044-0.044) | 0.064 (0.000-0.123) |
| Cell_fusing_agent virus | *Orthoflavivirus iunctionis* | 190 | 0.059 (0.005-0.116) | 0.022 (0.043-0.043) | 0.042 (0.000-0.064) |
| Dengue virus type 2 | *Orthoflavivirus denguei* | 190 | 0.023 (0.001-0.037) | 0.025 (0.043-0.043) | 0.066 (0.002-0.108) |
| Modoc virus | *Orthoflavivirus modocense* | 3 | 0.046 (0.036-0.054) | 0.026 (0.039-0.039) | 0.081 (0.003-0.122) |
| Karshi virus | *Orthoflavivirus royalense* | 3 | 0.051 (0.018-0.072) | 0.029 (0.043-0.043) | 0.099 (0.002-0.148) |
| TBEV | *Orthoflavivirus neudoerflense* | 190 | 0.044 (0.004-0.074) | 0.039 (0.072-0.072) | 0.108 (0.001-0.162) |
| Saumarez_Reef virus | *Orthoflavivirus saumarezense* | 1 | 0.061 (0.061-0.061) | 0.042 (0.042-0.042) | 0.104 (0.104-0.104) |
| Yokose virus | *Orthoflavivirus yokoseense* | 1 | 0.180 (0.180-0.180) | 0.143 (0.143-0.143) | 0.259 (0.259-0.259) |

[1]Mean distance (minimum – Maximum) values

**Table S5**. Sequence distances of selected isolates of sylvatic and divergent DENV isolates with human DENV strains

| | | | Sequence distances (mean, range) | | |
|---|---|---|---|---|---|
| **Type** | **Accession** | **Isolate** | **Polyprotein** | ***NS5*** | ***E*** |
| *Sylvatic DENV isolates* | | | | | |
| DENV1 | EF457905 | P72-1244 | 0.022 (0.019 - 0.025) | 0.019 (0.015 - 0.025) | 0.023 (0.016 - 0.030) |
| DENV2 | EF105379 | P8-1407 | 0.066 (0.062 - 0.071) | 0.073 (0.064 - 0.081) | 0.047 (0.040 - 0.055) |
| DENV4 | EF457906 | P75-215 | 0.046 (0.041 - 0.060) | 0.045 (0.039 - 0.071) | 0.045 (0.038 - 0.053) |
| DENV4 | JF262779 | P75-514 | 0.046 (0.041 - 0.059) | 0.043 (0.037 - 0.069) | 0.045 (0.038 - 0.053) |
| DENV4 | JF262780 | P73-1120 | 0.046 (0.041 - 0.059) | 0.043 (0.037 - 0.069) | 0.045 (0.038 - 0.053) |
| *Divergent DENV isolates* | | | | | |
| DENV1 | KR919820 | Brun2014 | 0.066 (0.063 - 0.070) | 0.050 (0.047 - 0.056) | 0.066 (0.059 - 0.071) |
| DENV2 | FJ467493 | DKD811 | 0.068 (0.063 - 0.073) | 0.073 (0.063 - 0.082) | 0.050 (0.042 - 0.059) |
| DENV2 | KX274130 | QML22 | 0.140 (0.138 - 0.143) | 0.132 (0.128 - 0.136) | 0.095 (0.089 - 0.105) |
| DENV4 | OK605599 | DKE-121 | 0.149 (0.146 - 0.160) | 0.157 (0.152 - 0.176) | 0.111 (0.103 - 0.119) |

**Table S6**. Between-strain sequence divergence of named orthoflaviviruses assigned to the same family

| ***Species*** | **Virus_1** | **Virus_2** | **Count** | ***PP* AA distance[1]** | ***NS5* AA distance[1]** | ***E* AA distance[1]** |
|---|---|---|---|---|---|---|
| *Orthoflavivirus aroaense* | Aroa | Iguape | 4 | 0.110 (0.106-0.114) | 0.087 (0.084-0.090) | 0.094 (.089-0.099) |
| | | Bussuquara | 2 | 0.383 (0.383-0.383) | 0.277 (0.276-0.277) | 0.415 (.413-0.417) |
| | | Naranjal | 2 | 0.384 (0.384-0.384) | 0.274 (0.274-0.275) | 0.425 (.425-0.425) |
| | Naranjal | Bussuquara | 1 | 0.132 (0.132-0.132) | 0.102 (0.102-0.102) | 0.144 (.144-0.144) |
| | | Iguape | 2 | 0.383 (0.381-0.386) | 0.271 (0.268-0.275) | 0.431 (.431-0.431) |
| | Iguape | Bussuquara | 2 | 0.387 (0.384-0.389) | 0.269 (0.266-0.272) | 0.421 (.421-0.421) |
| *Orthoflavivirus denguei* | DENV_3 | DENV_1 | 400 | 0.217 (0.213-0.225) | 0.184 (0.176-0.191) | 0.220 (.215-0.231) |
| | | DENV_2 | 400 | 0.277 (0.272-0.282) | 0.209 (0.201-0.219) | 0.315 (.308-0.323) |
| | | DENV_4 | 400 | 0.299 (0.294-0.310) | 0.232 (0.226-0.252) | 0.368 (.360-0.373) |
| | DENV_2 | DENV_1 | 400 | 0.277 (0.273-0.286) | 0.205 (0.198-0.211) | 0.307 (.304-0.311) |
| | | DENV_4 | 400 | 0.305 (0.301-0.318) | 0.252 (0.243-0.275) | 0.363 (.354-0.376) |
| | DENV_4 | DENV_1 | 400 | 0.309 (0.305-0.329) | 0.242 (0.238-0.262) | 0.361 (.354-0.372) |
| *Orthoflavivirus kokoberaorum* | Kokobera | Bainyik | 2 | 0.077 (0.073-0.082) | 0.083 (0.081-0.086) | 0.061 (.056-0.066) |
| | | New_Mapoon | 2 | 0.291 (0.289-0.293) | 0.251 (0.250-0.251) | 0.242 (.242-0.242) |
| | | Stratford | 10 | 0.155 (0.150-0.161) | 0.159 (0.156-0.163) | 0.154 (.148-0.160) |
| | | Torres | 2 | 0.144 (0.140-0.147) | 0.137 (0.136-0.138) | 0.113 (.108-0.119) |
| | Stratford | Torres | 5 | 0.110 (0.109-0.113) | 0.113 (0.112-0.115) | 0.093 (.092-0.094) |
| | | Bainyik | 5 | 0.148 (0.147-0.150) | 0.153 (0.153-0.153) | 0.139 (.138-0.140) |
| | | New_Mapoon | 5 | 0.289 (0.288-0.291) | 0.239 (0.237-0.241) | 0.238 (.238-0.238) |
| | Torres | Bainyik | 1 | 0.132 (0.132-0.132) | 0.120 (0.120-0.120) | 0.096 (.096-0.096) |
| | | New_Mapoon | 1 | 0.282 (0.282-0.282) | 0.239 (0.239-0.239) | 0.247 (.246-0.248) |
| | New_Mapoon | Bainyik | 1 | 0.283 (0.283-0.283) | 0.241 (0.241-0.241) | 0.237 (.237-0.237) |

| ***Species*** | **Virus_1** | **Virus_2** | **Count** | ***PP* AA distance[1]** | ***NS5* AA distance[1]** | ***E* AA distance[1]** |
|---|---|---|---|---|---|---|
| *Orthoflavivirus entebbeense* | Sokuluk | Entebbe_bat | 6 | 0.216 (0.210-0.223) | 0.177 (0.175-0.180) | 0.176 (.173-0.180) |
| *Orthoflavivirus ilheusense* | Rocio | Ilheus | 36 | 0.218 (0.216-0.225) | 0.195 (0.191-0.201) | 0.205 (.198-0.214) |
| *Orthoflavivirus kyasanurense* | KFDV | Alkhumra | 84 | 0.037 (0.029-0.043) | 0.024 (0.021-0.027) | 0.017 (.012-0.022) |
| *Orthoflavivirus murrayense* | MVEV | Alfuy | 19 | 0.163 (0.160-0.167) | 0.136 (0.133-0.139) | 0.151 (.144-0.156) |
| *Orthoflavivirus phnompenhense* | Phnom_Penh_bat | Batu_Cave | 2 | 0.065 (0.063-0.067) | 0.023 (0.022-0.024) | 0.033 (.031-0.035) |
| *Orthoflavivirus powassanense* | Powassan | Deer_tick | 36 | 0.015 (0.002-0.056) | 0.008 (0.001-0.038) | 0.007 (.000-0.042) |
| *Orthoflavivirus royalense* | Royal_Farm | Karshi | 3 | 0.294 (0.292-0.295) | 0.183 (0.181-0.187) | 0.314 (.308-0.320) |
| *Orthoflavivirus saboyaense* | Saboya | Potiskum | 1 | 0.010 (0.010-0.010) | 0.003 (0.003-0.003) | 0.008 (.008-0.008) |

[1]Mean distance (minimum – Maximum) values

SUPPLEMENTARY FIGURES

**Fig. S1.** Labelled phylogenetic tree of *Orthoflavivirus NS5* gene sequences (Fig. 1 and Suppl. file (NS5.NWK)

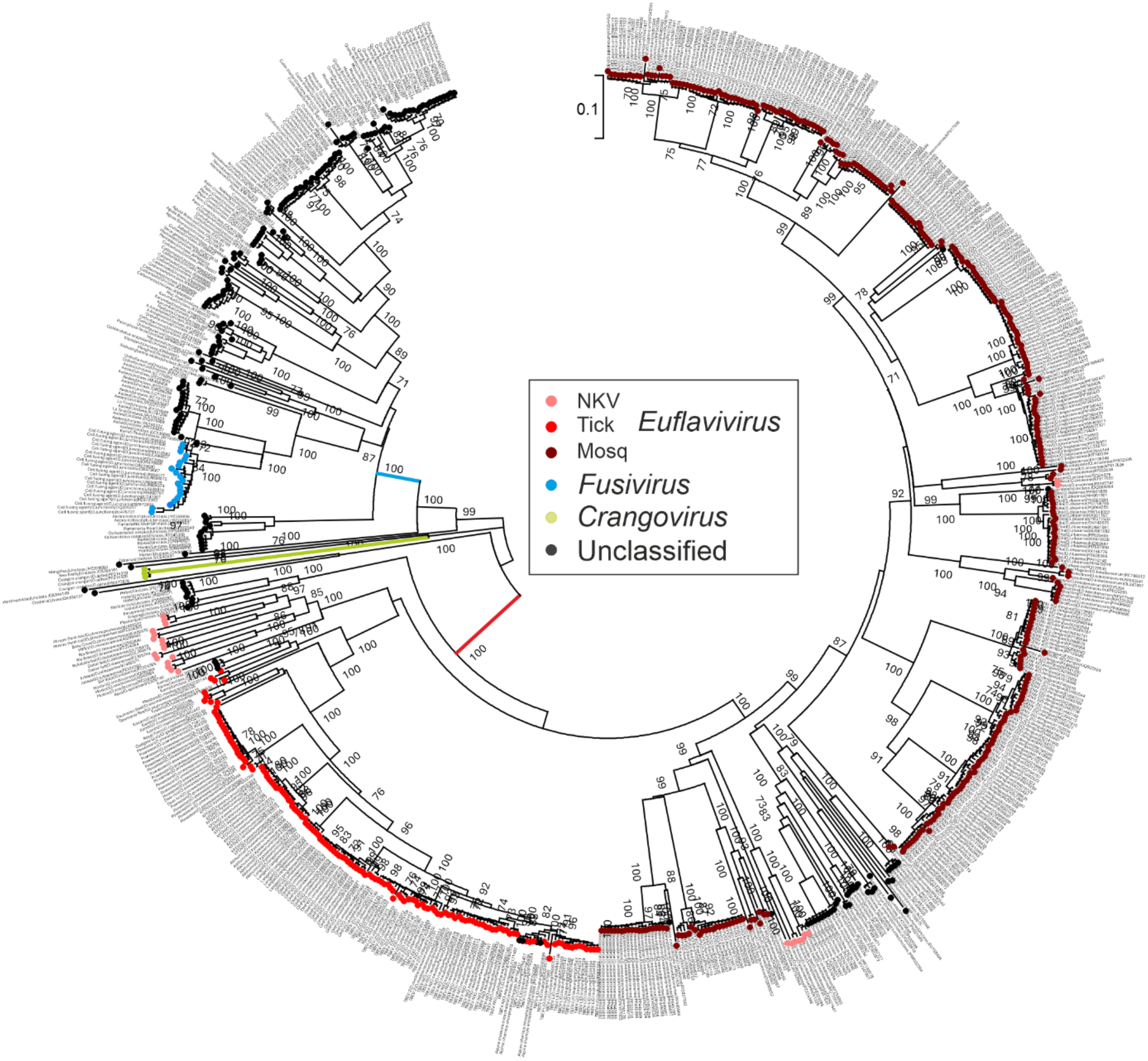

**Fig. S2**. Distributions of *E* gene (A) and (B) polyprotein p-distances between (left) and within (right) species of currently classified orthoflaviviruses. Assignment thresholds of 0.10 and 0.135 for *E* and polyprotein respectively splits within and between-species distances equivalently to the 0.10 threshold for *NS5* distances (Fig., 2). Grey and blue shading of bars as in Fig. 2

A) *E* gene

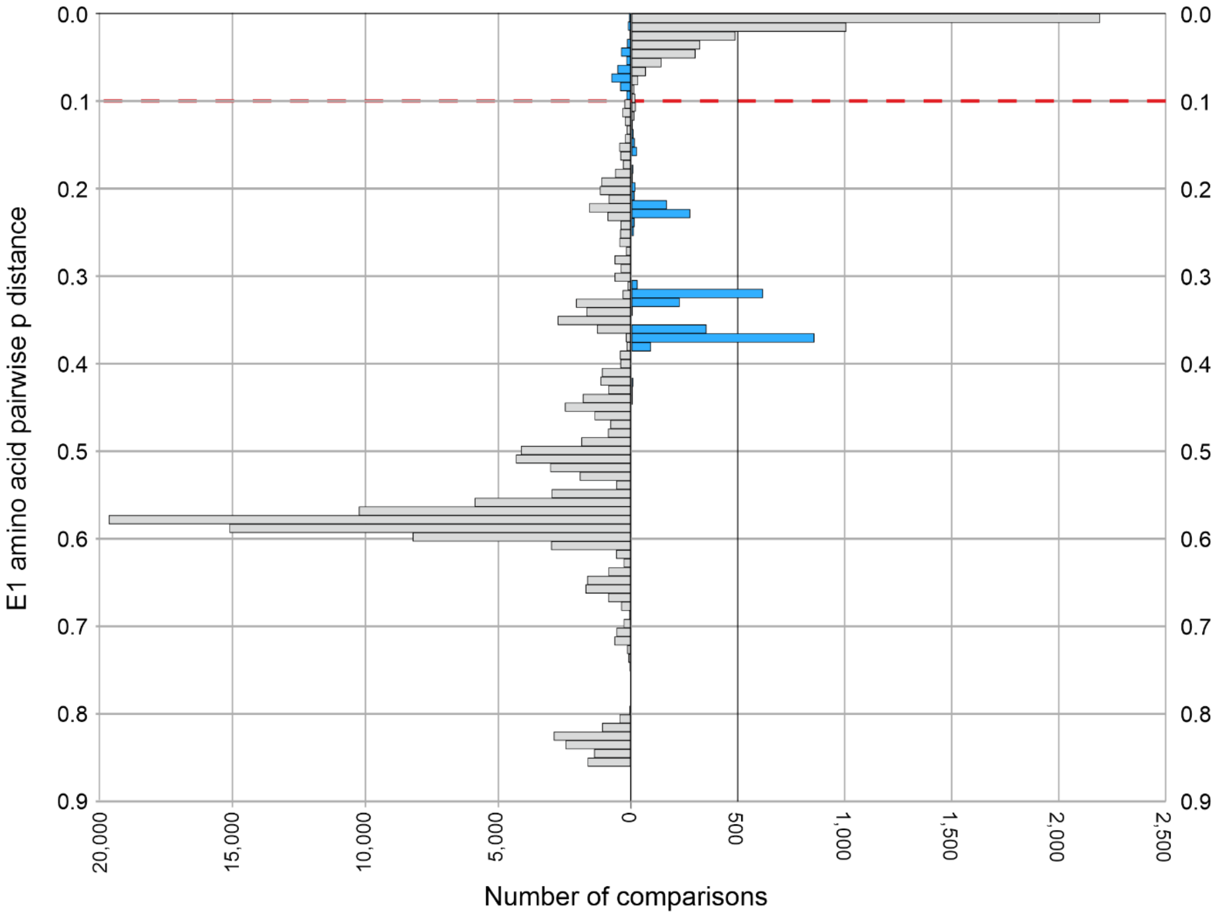

B) Complete polyprotein

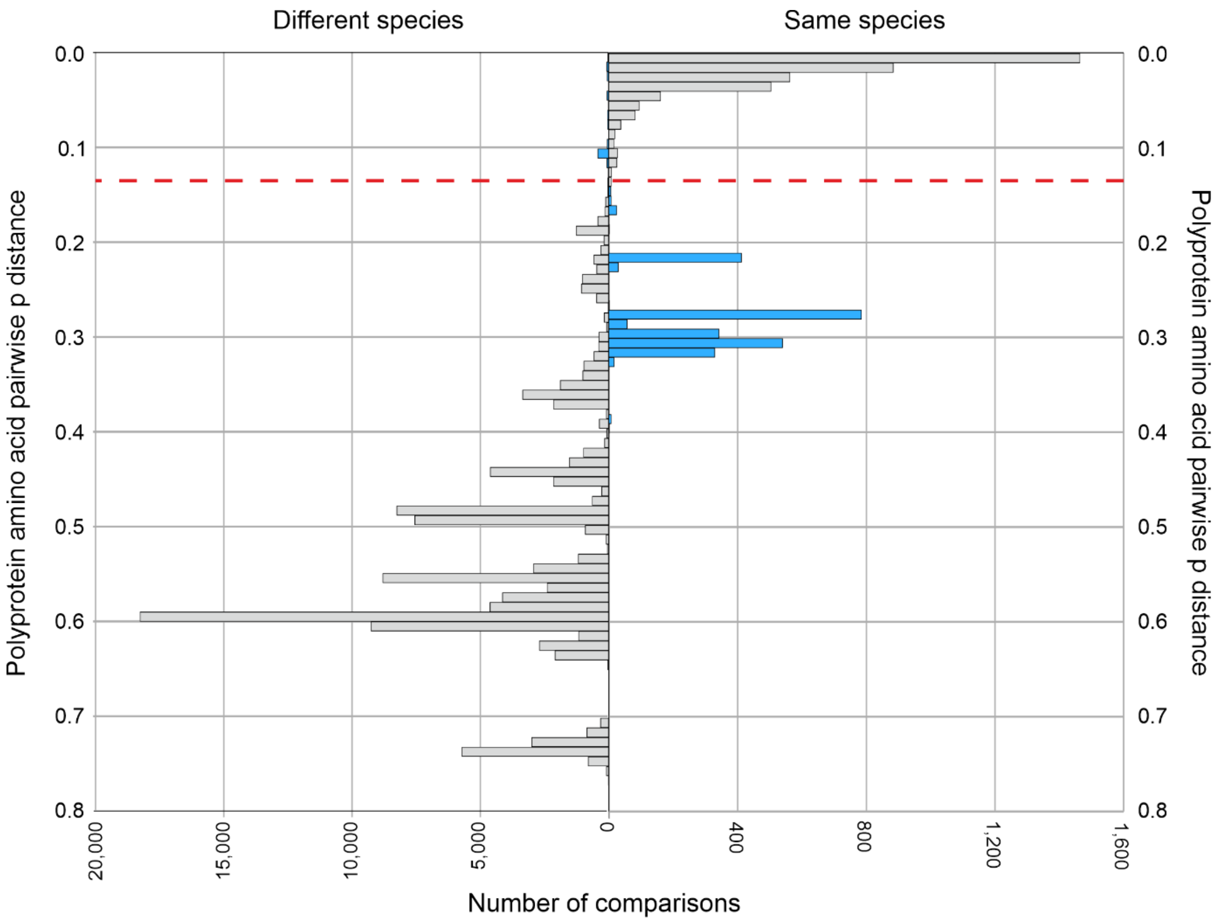
